# Identification of biomarkers, pathways and potential therapeutic targets for heart failure using bioinformatics analysis

**DOI:** 10.1101/2021.08.05.455244

**Authors:** Basavaraj Vastrad, Chanabasayya Vastrad

## Abstract

Heart failure (HF) is a complex cardiovascular diseases associated with high mortality. To discover key molecular changes in HF, we analyzed next-generation sequencing (NGS) data of HF. In this investigation, differentially expressed genes (DEGs) were analyzed using limma in R package from GSE161472 of the Gene Expression Omnibus (GEO). Then, gene enrichment analysis, protein-protein interaction (PPI) network, miRNA-hub gene regulatory network and TF-hub gene regulatory network construction, and topological analysis were performed on the DEGs by the Gene Ontology (GO), REACTOME pathway, STRING, HiPPIE, miRNet, NetworkAnalyst and Cytoscape. Finally, we performed receiver operating characteristic curve (ROC) analysis of hub genes. A total of 930 DEGs 9464 up regulated genes and 466 down regulated genes) were identified in HF. GO and REACTOME pathway enrichment results showed that DEGs mainly enriched in localization, small molecule metabolic process, SARS-CoV infections and the citric acid (TCA) cycle and respiratory electron transport. Subsequently, the PPI network, miRNA-hub gene regulatory network and TF-hub gene regulatory network were constructed, and 10 hub genes in these network were focused on by centrality analysis and module analysis. Furthermore, data showed that HSP90AA1, ARRB2, MYH9, HSP90AB1, FLNA, EGFR, PIK3R1, CUL4A, YEATS4 and KAT2B were good diagnostic values. In summary, this study suggests that HSP90AA1, ARRB2, MYH9, HSP90AB1, FLNA, EGFR, PIK3R1, CUL4A, YEATS4 and KAT2B may act as the key genes in HF.

## Introduction

Heart failure (HF) is one of the chronic cardiovascular diseases, affecting 1% to 2% of the adult population worldwide [1]. HF is said to be inefficiency of the heart to supply the peripheral tissues with the appropriate amount of blood and oxygen to meet their metabolic requirement and is linked with a high risk for subsequent mortality and morbidity [2]. Multiple risk factors might cause HF, including diabetics [3], hypertension [4], obesity [5], genetics [6], environmental triggers [7], and immunity, inflammation, and oxidative stress [8]. Although there are extensive investigation available regarding the etiologies and mechanisms underlying HF, the precise molecular mechanisms remain unclear [9–10]. Therefore, essential molecular markers of HF that are identifiable with more powerful technologies are urgently required.

Understanding the status of various genes and signaling pathway in early diagnosis of HF could improve the effect of initial treatment. COL1A1 [11], CXCL14 [12], MECP2 [13], RBM20 [14], PGC-1 [15], Wnt signaling pathway [16], TGF□β1/Smad3 signaling pathway [17], AT1-CARP signaling pathway [18], Akt signaling pathway [19] and neuregulin-1/ErbB signaling [20] were responsible for progression of HF. Therefore, we aimed to further explore the molecular pathogenesis of HF and identify specific molecular targets. However, these data still demand further clinical interpretation.

Next-generation sequencing (NGS) technology plays a crucial role in the analysis of gene expression, which served as important tools in cardiovascular research with great clinical application [21]. Recently, a large number of gene expression profiling studies have been reported with the use of NGS technology. The integrated bioinformatics analysis will be more positive and provide valuable novel molecular targets to foster the advancement of specific diagnosis and new therapeutic strategies.

In this investigation, NGS dataset (GSE161472) was downloaded from the GEO database (http://www.ncbi.nlm.nih.gov/geo/) [22], and crucial genes identified by combining bioinformatics analyses in HF. Gene ontology (GO) terms and REACTOME pathways associated with HF were investigated, and the hub genes associated with HF were identified by protein–protein interaction (PPI) network, modules, miRNA-hub gene regulatory network and TF-hub gene regulatory network construction and analysis. Subsequently, we validated the hub genes by receiver operating characteristic curve (ROC) analysis. Furthermore, we investigated the potential candidate molecular markers for their utility in diagnosis, prognosis, and drug targeting in HF.

## Material and methods

### Data resources

This study investigated DEGs in HF versus normal samples by analyzing GSE161472 GEO expression profiling by high throughput sequencing data downloaded from the GEO database. GEO serves as a public repository for experimental high-throughput raw NGS data. Expression profiling by high throughput sequencing profile was generated with the GPL11154 Illumina HiSeq 2000 (Homo sapiens). The GSE161472 dataset included 84 samples, containing 47 HF and 37 normal control samples.

### Identification of DEGs

The analysis of screening DEGs between HF and normal control samples was analyzed by limma in R package [23]. Moreover, the threshold for the DEGs was set as P-value <0 .05, and |log2foldchange (FC)| > 0.22 for up regulated genes and|log2foldchange (FC)| < -0.18 for down regulated genes. The heat map and volcano plot of the DEGs were plotted using gplots and ggplot2, respectively.

### GO and REACTOME pathway enrichment analysis of DEGs

The GO terms (http://www.geneontology.org) database primarily adds three categories: biological process (BP), cellular component (CC), and molecular function (MF) [24]. The REACTOME pathway (https://reactome.org/) [25] database compiles genomic, chemical, and systematic functional information. The g:Profiler (http://biit.cs.ut.ee/gprofiler/) [26] online tool implements methods to analyze and anticipate functional profiles of gene and gene clusters. In this investigation, GO terms and REACTOME pathways were analyzed using the g:Profiler with the enrichment threshold of P < 0.05.

### Construction of the PPI network and module analysis

The Human Integrated Protein-Protein Interaction rEference (HiPPIE) interactome (http://cbdm-01.zdv.uni-mainz.de/~mschaefer/hippie/) [27] database provides a significant association of protein□protein interaction (PPI). Cytoscape 3.8.2 (http://www.cytoscape.org/) [28] is used for the visual exploration of interaction networks. In this investigation, DEGs PPI networks were analyzed by the HiPPIE database and subsequently visualized by using Cytoscape. In addition, the node degree [29], betweenness centrality [30], stress centrality [31] and closeness centrality [32] of each protein node in the PPI network was calculated using plug-in Network Analyzer of the Cytoscape software. PEWCC1 (http://apps.cytoscape.org/apps/PEWCC1) [33] plug-in of the Cytoscape software was then used to screen out modules of PPI networks, and the degree cutoff = 2, node score cutoff = 0.2, k core = 2, and max depth = 100.

### MiRNA-hub gene regulatory network construction

The miRNet database (https://www.mirnet.ca/) [34], a web biological database for prediction of known and unknown miRNA and hub genes relationships, was used to construct the miRNA-hub gene regulatory network, which was visualized in Cytoscape 3.8.2 [28].

### TF-hub gene regulatory network construction

TF-hub gene regulatory network analysis is useful to analyze the interactions between hub genes and TF which might provide insights into the mechanisms of generation or development of diseases. NetworkAnalyst database (https://www.networkanalyst.ca/) [35] and Cytoscape 3.8.2 [28] software were used to build the TF-hub gene regulatory network.

### Validation of hub genes by receiver operating characteristic curve (ROC) analysis

Then ROC curve analysis was implemented to calculate the sensitivity (true positive rate) and specificity (true negative rate) of the hub gens for HF diagnosis and we investigated how large the area under the curve (AUC) was by using pROC package in R statistical software [36]. The diagnostic values of the hub genes were predicted based on the ROC curve analysis.

## Results

### Identification of DEGs

Dataset GSE161472 was downloaded from the GEO database and analyzed using R packages (limma). Volcano plot was constructed to visualize fold changes of the DEGs (Fig. 1). A total of 930 DEGs were identified in GSE161472, among which a total of 464 were up regulated genes and 466 were down regulated genes and are listed in Table 1. The DEGs in the NSG data are shown as volcano plots in Fig.1. The 464 up regulated genes and 466 down regulated genes are shown in heatmap and were shown in Fig. 2.

**Fig. 1.**
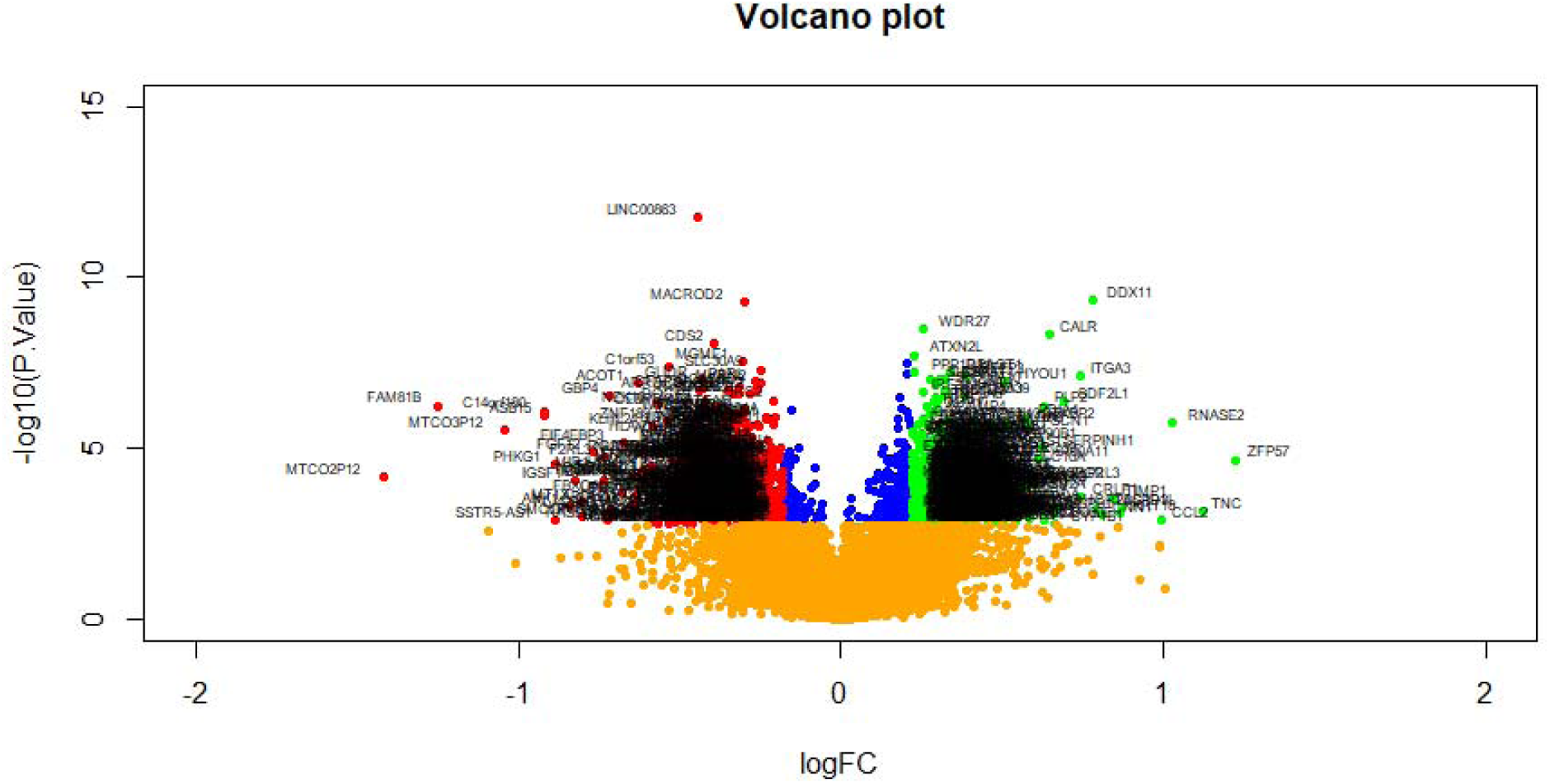
Volcano plot of differentially expressed genes. Genes with a significant change of more than two-fold were selected. Green dot represented up regulated significant genes and red dot represented down regulated significant genes.

**Fig. 2.**
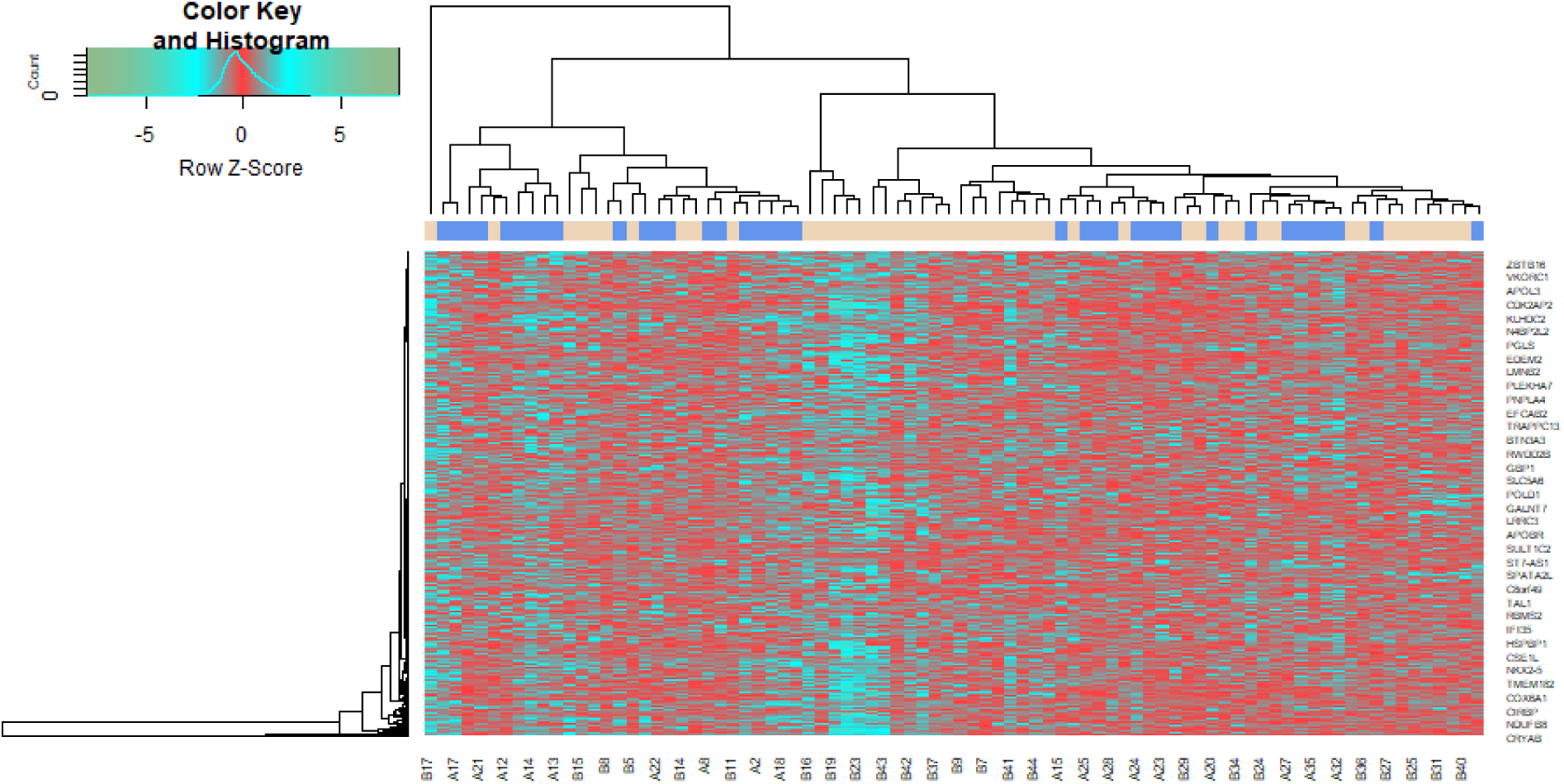
Heat map of differentially expressed genes. Legend on the top left indicate log fold change of genes. (A1 – A37 = normal control samples; B1 – B47 = HF samples)

**Table 1.**
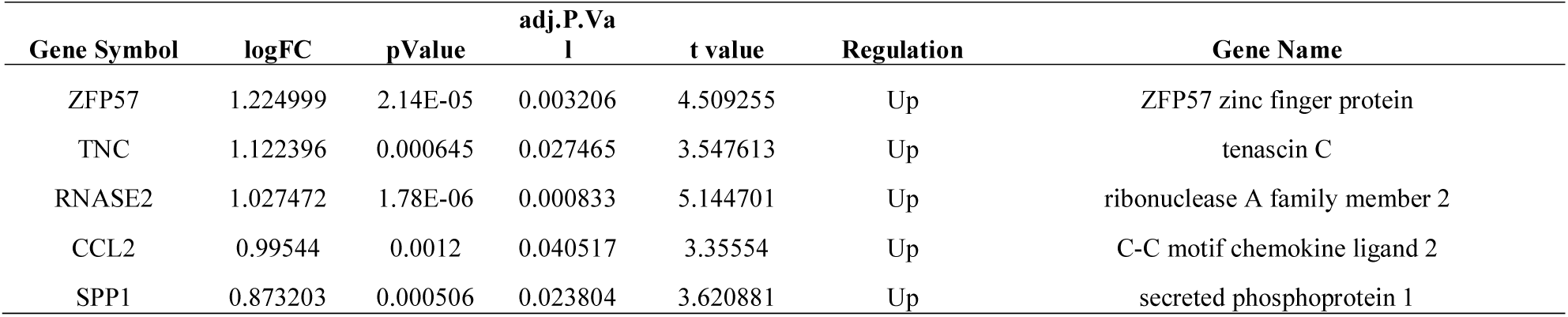
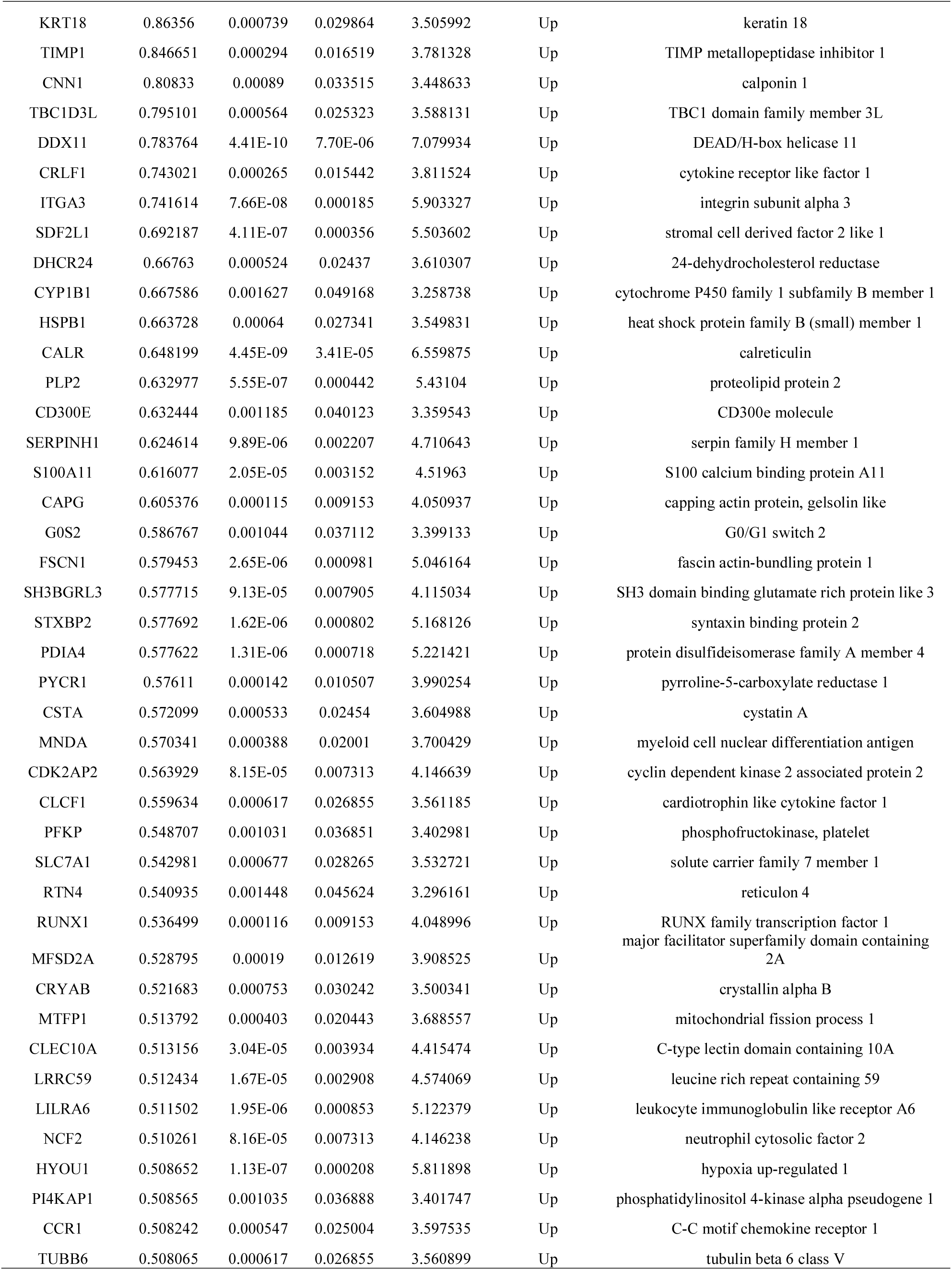
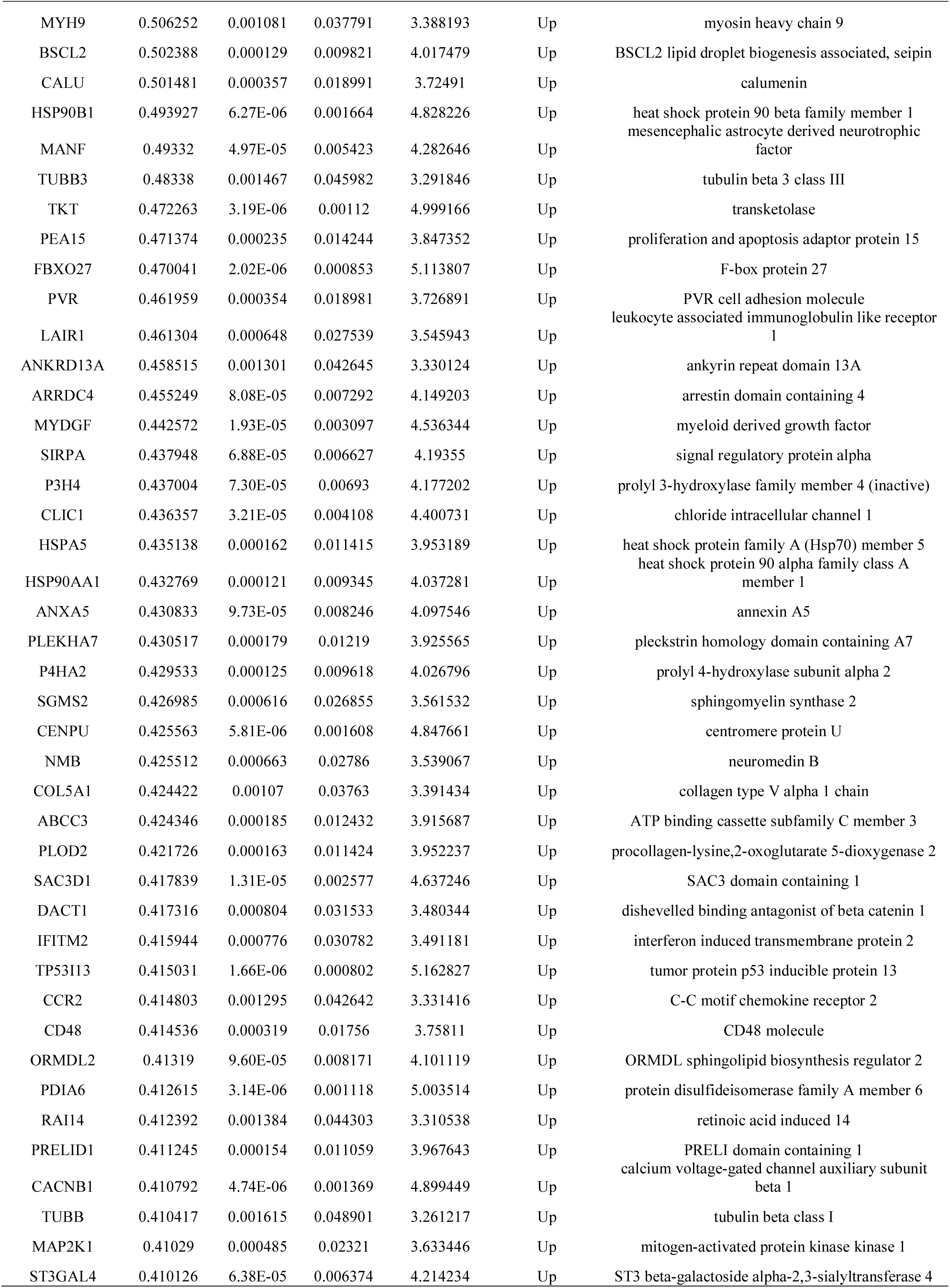
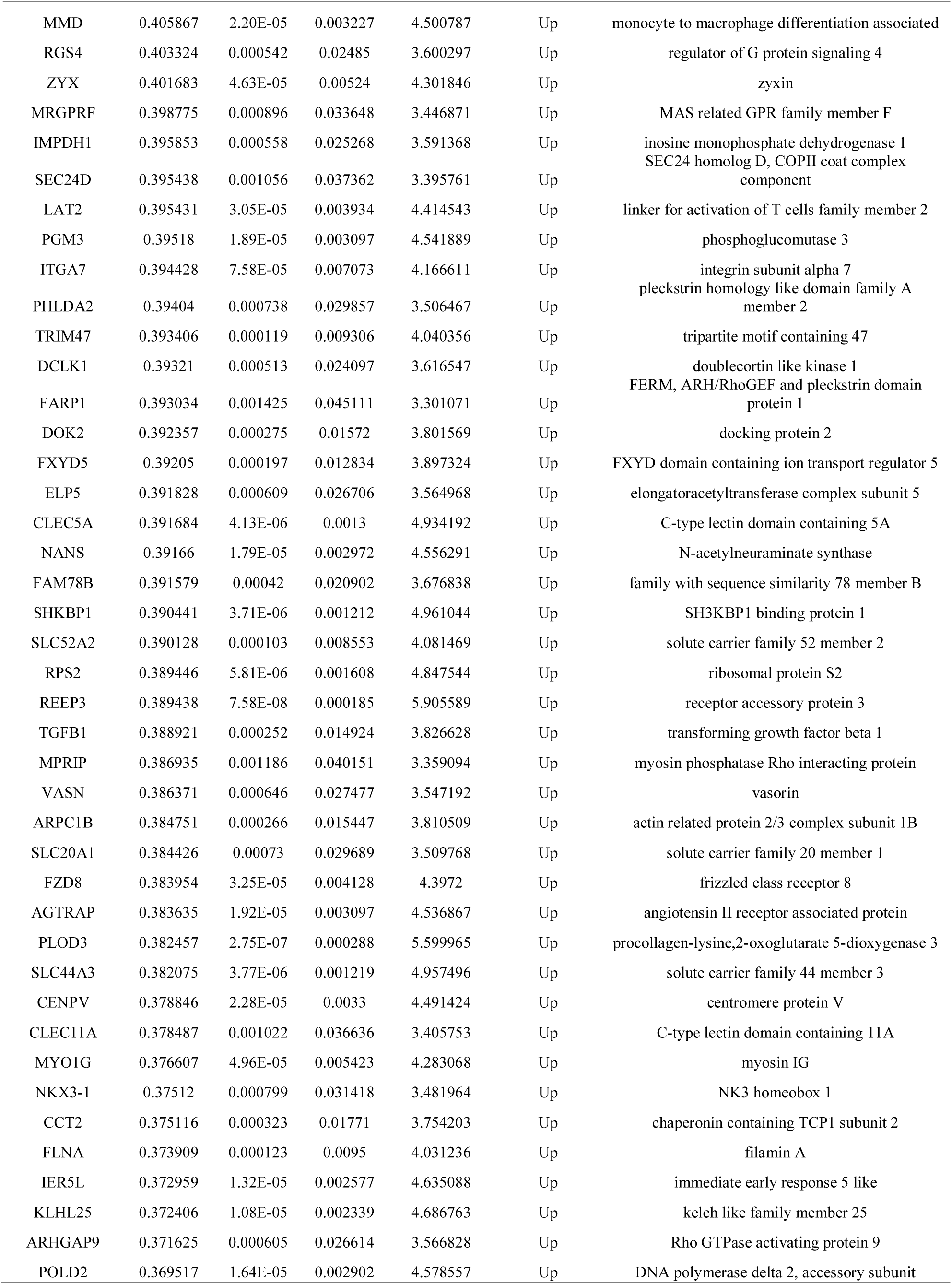
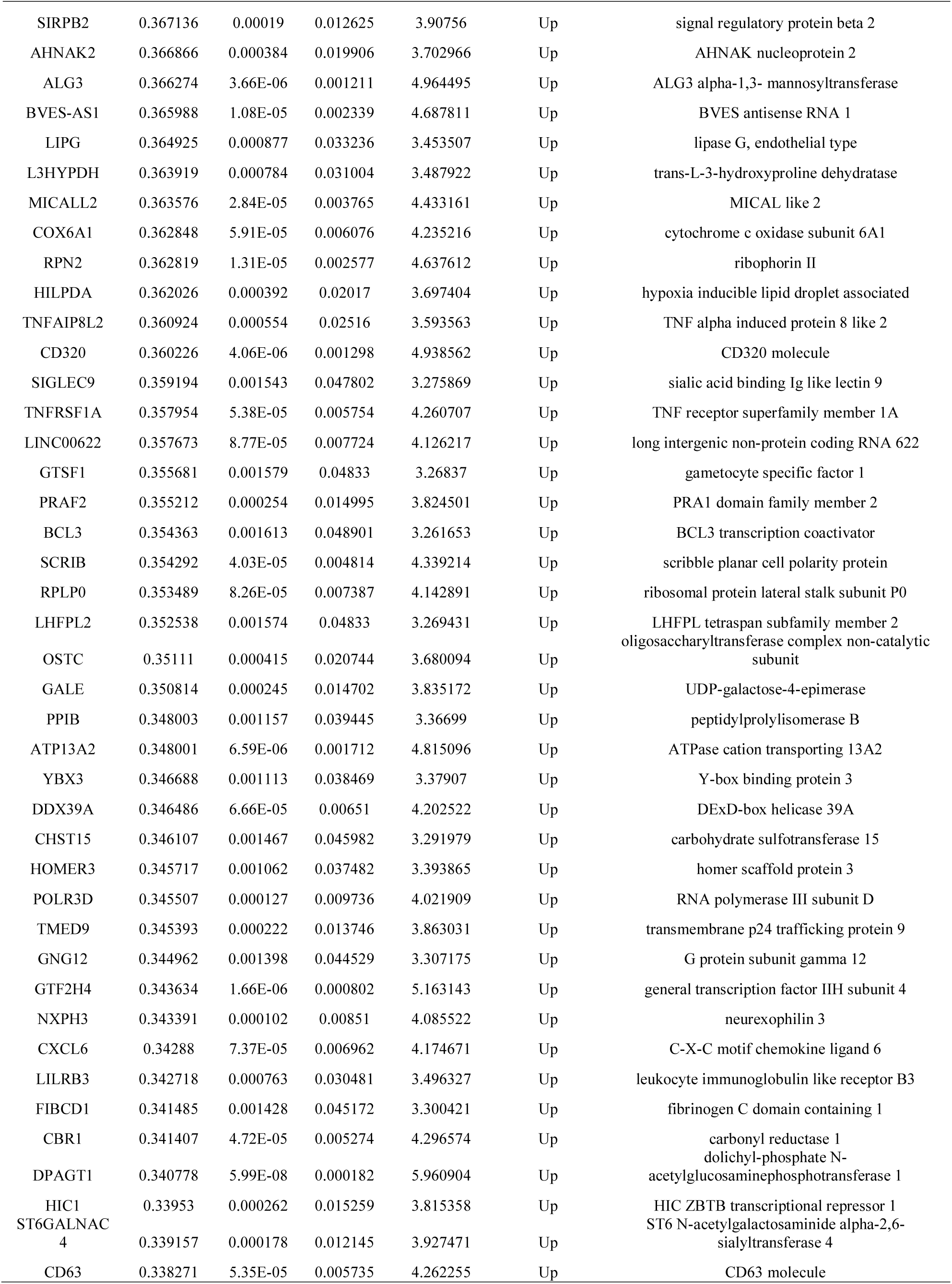
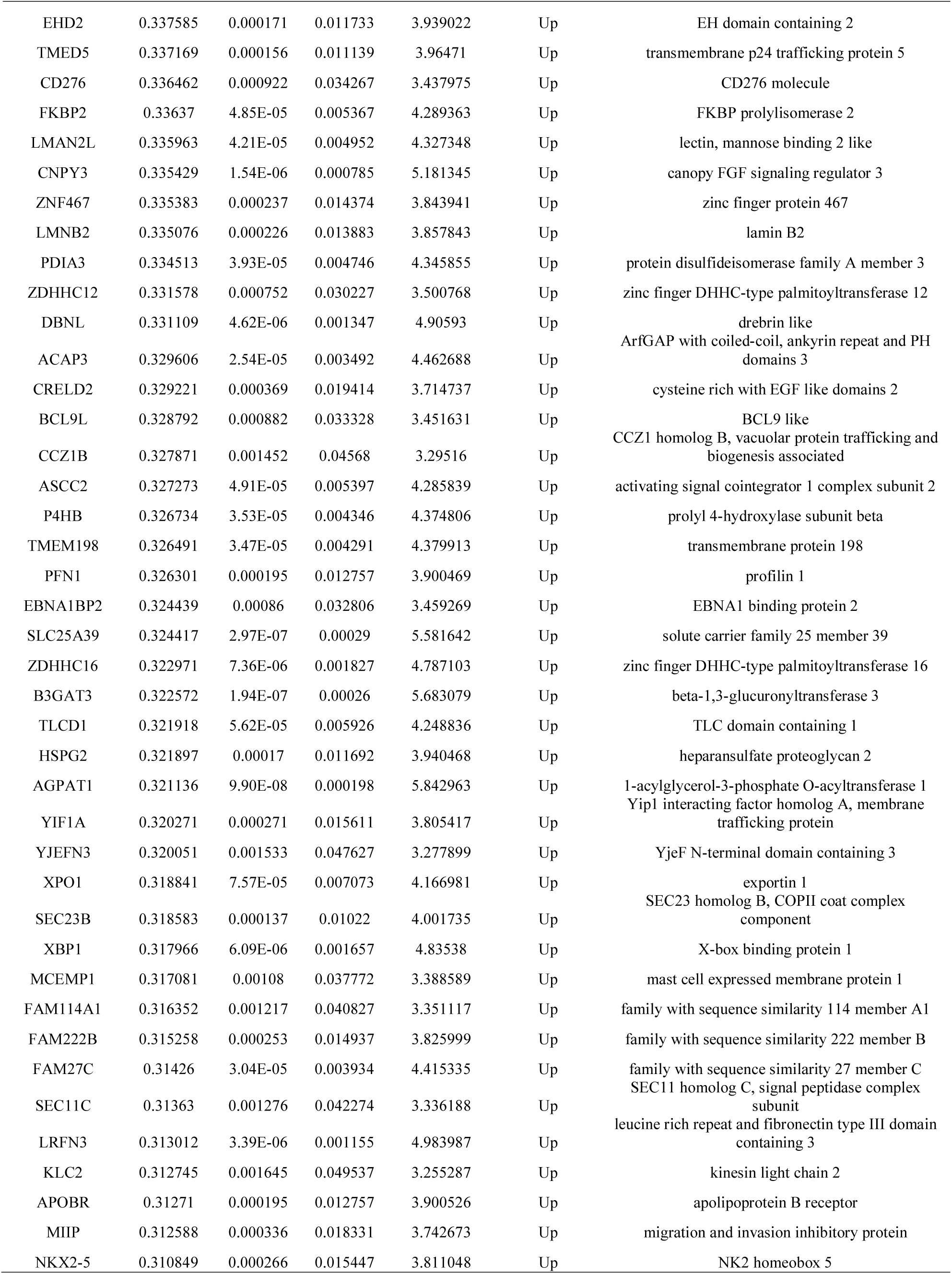
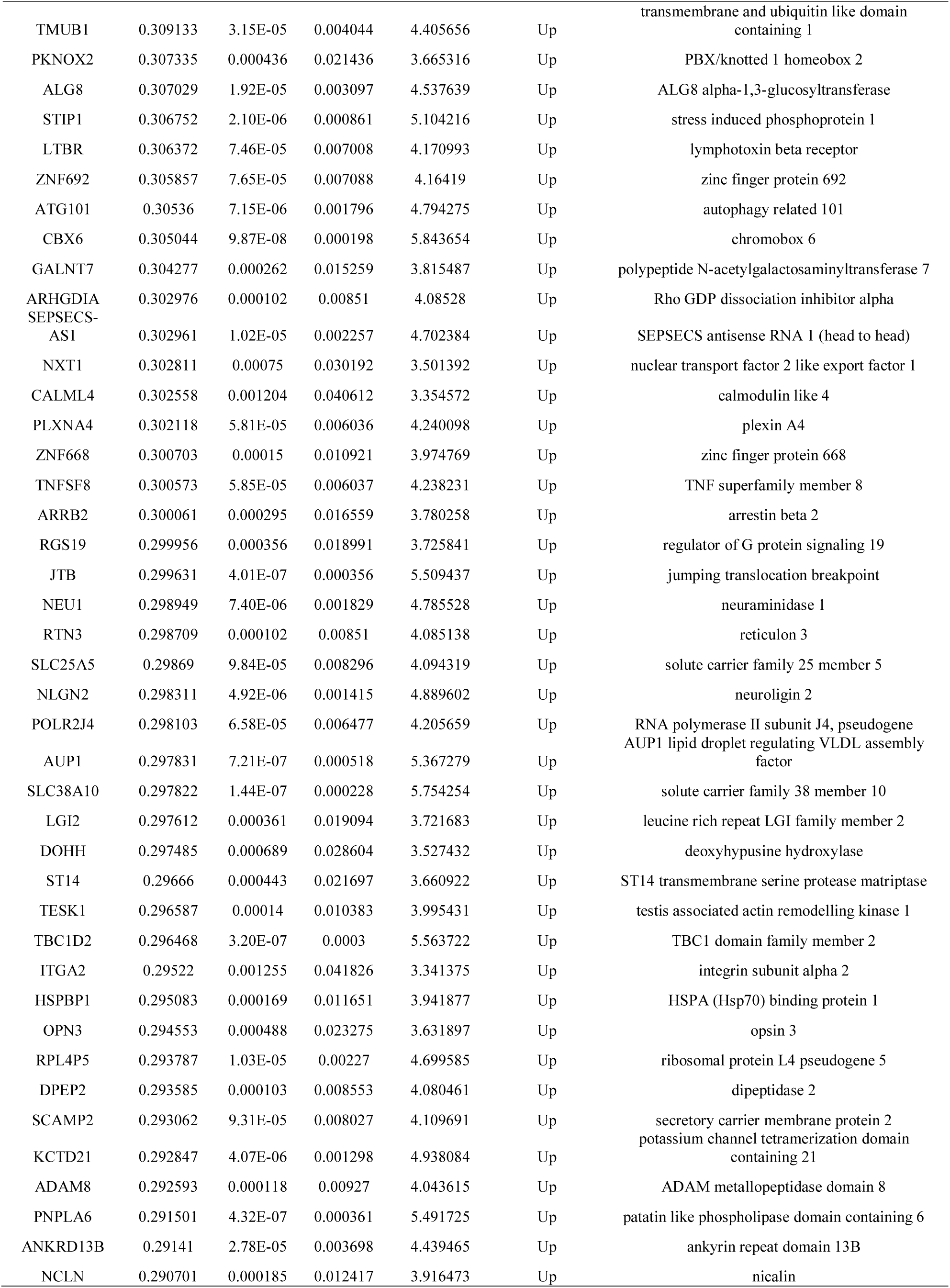
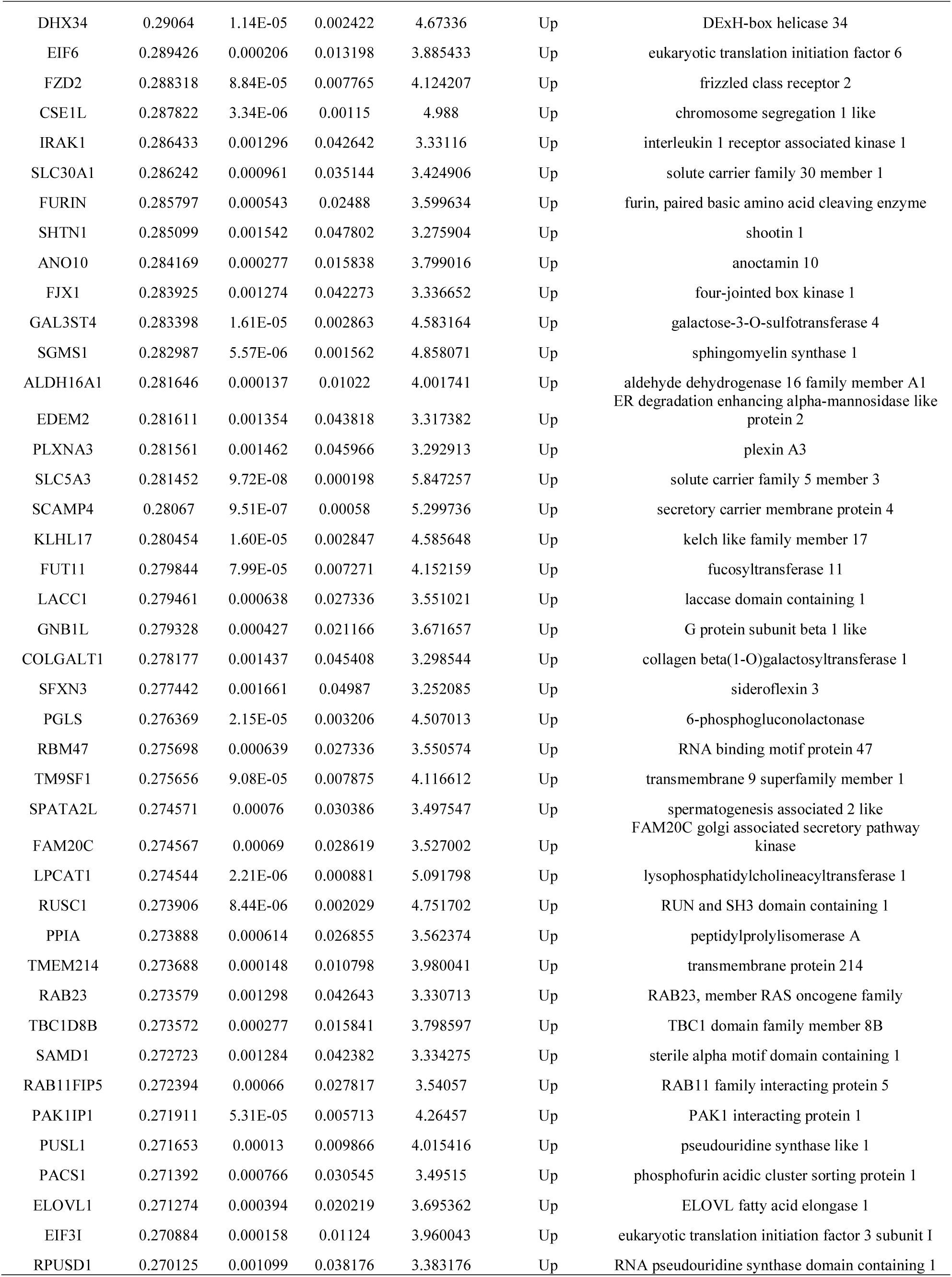
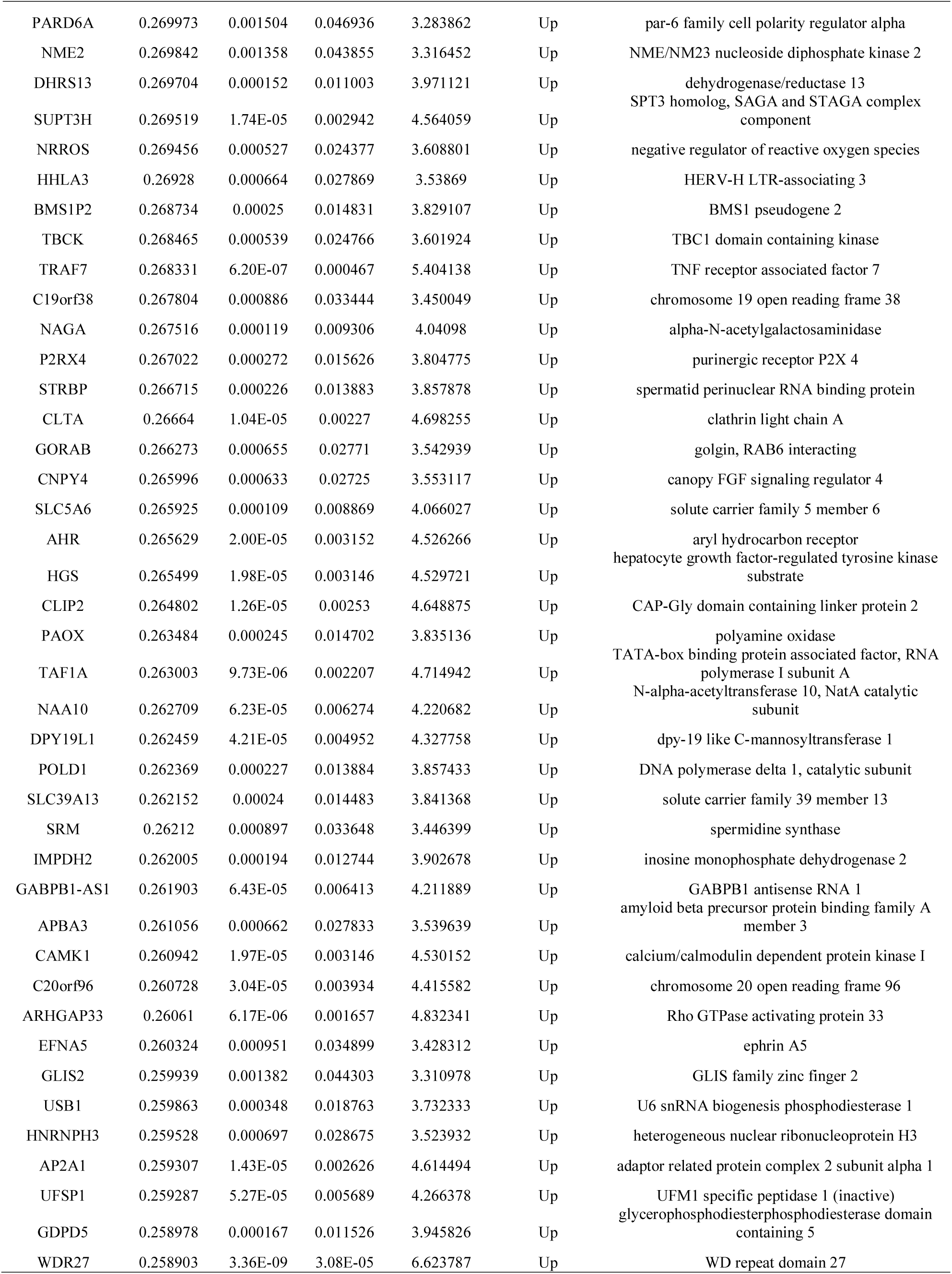
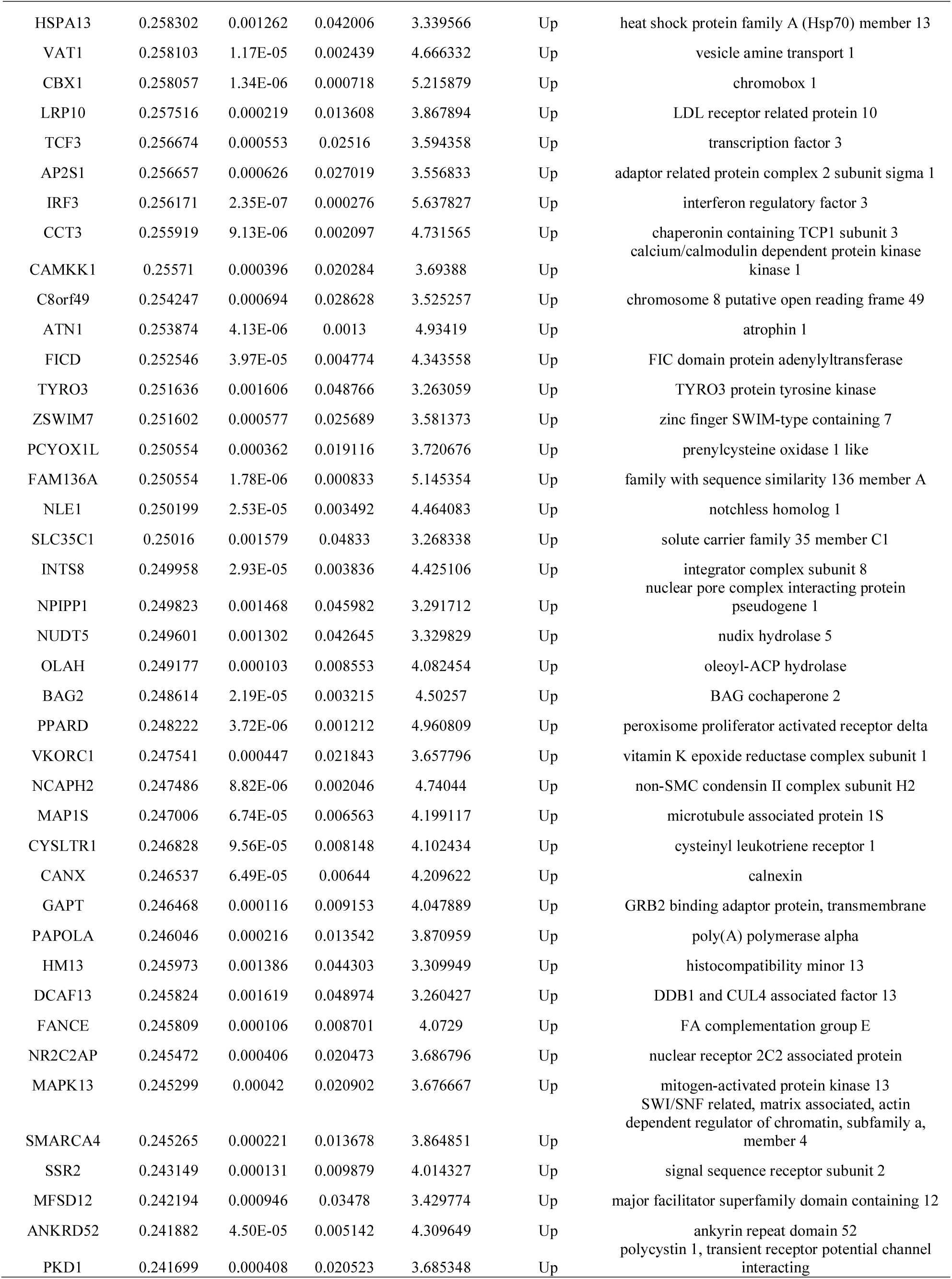
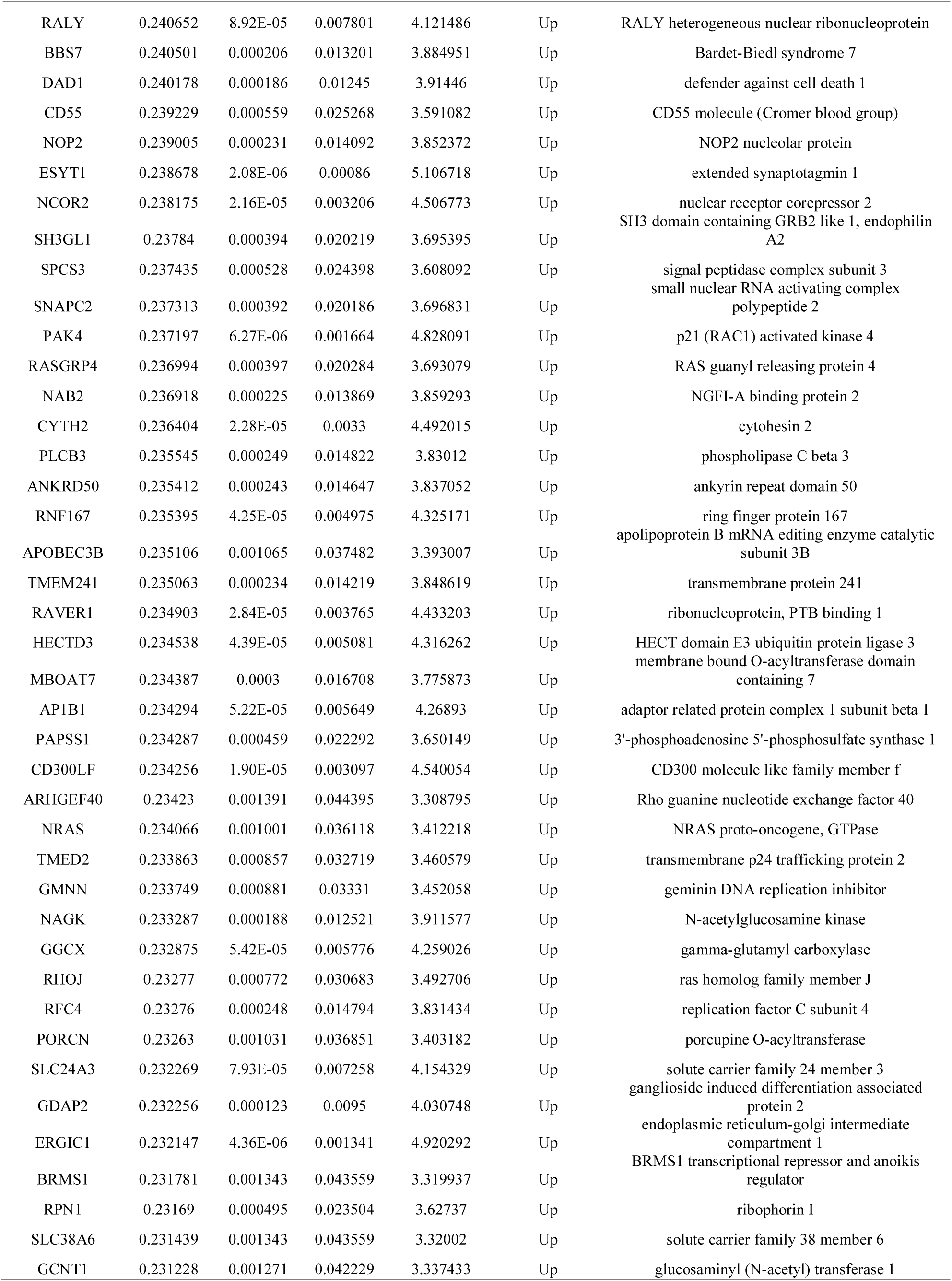
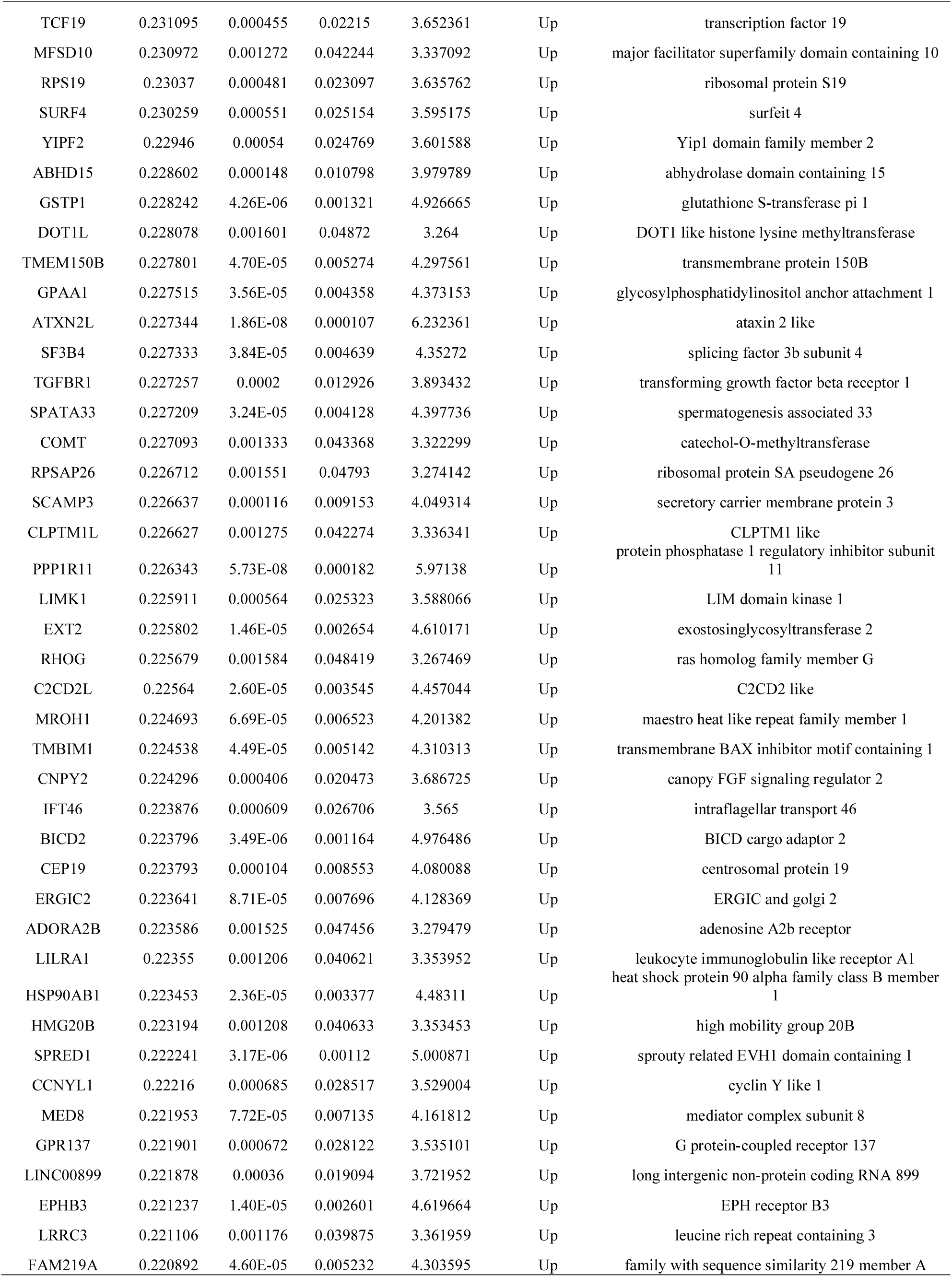
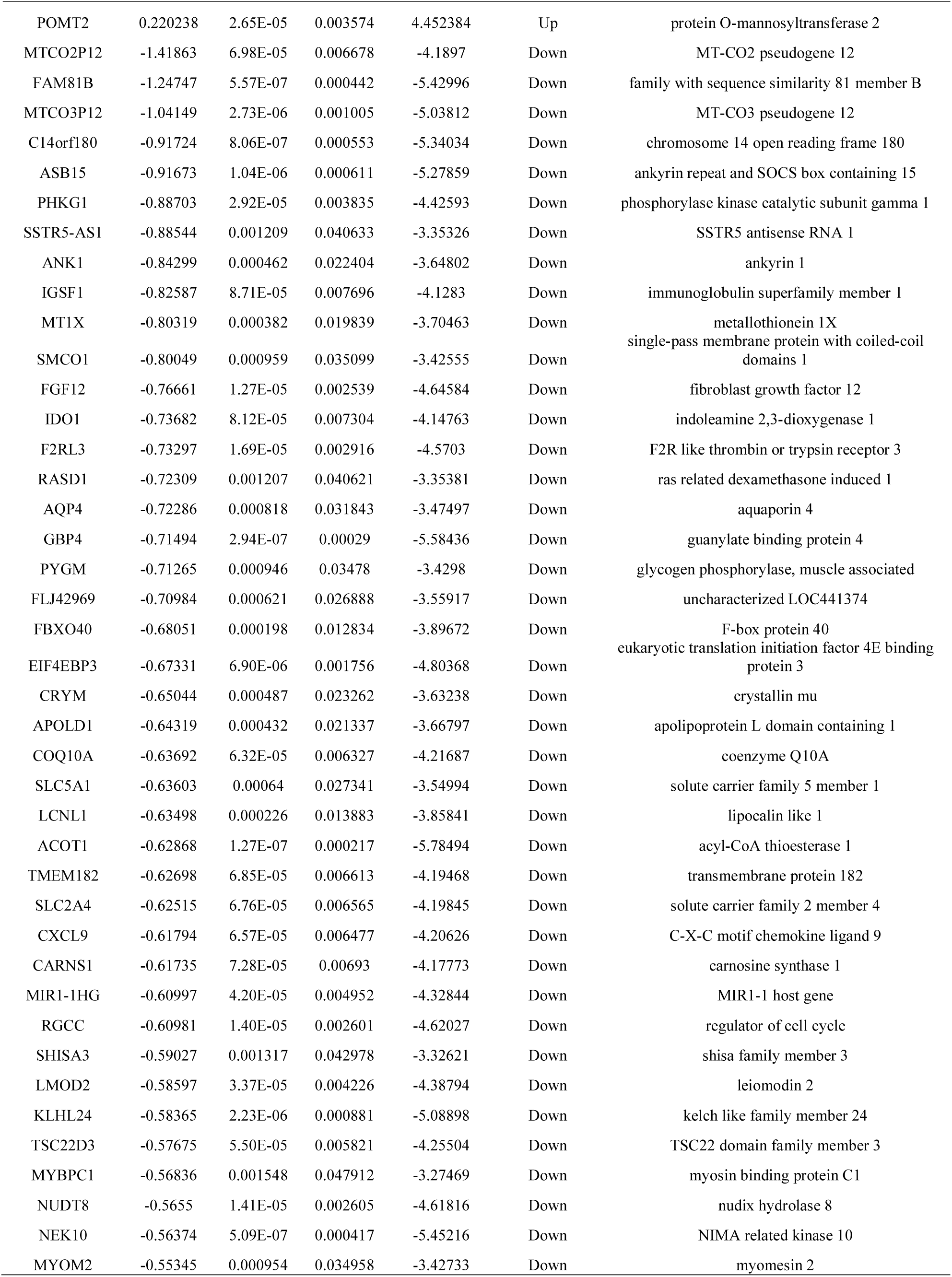
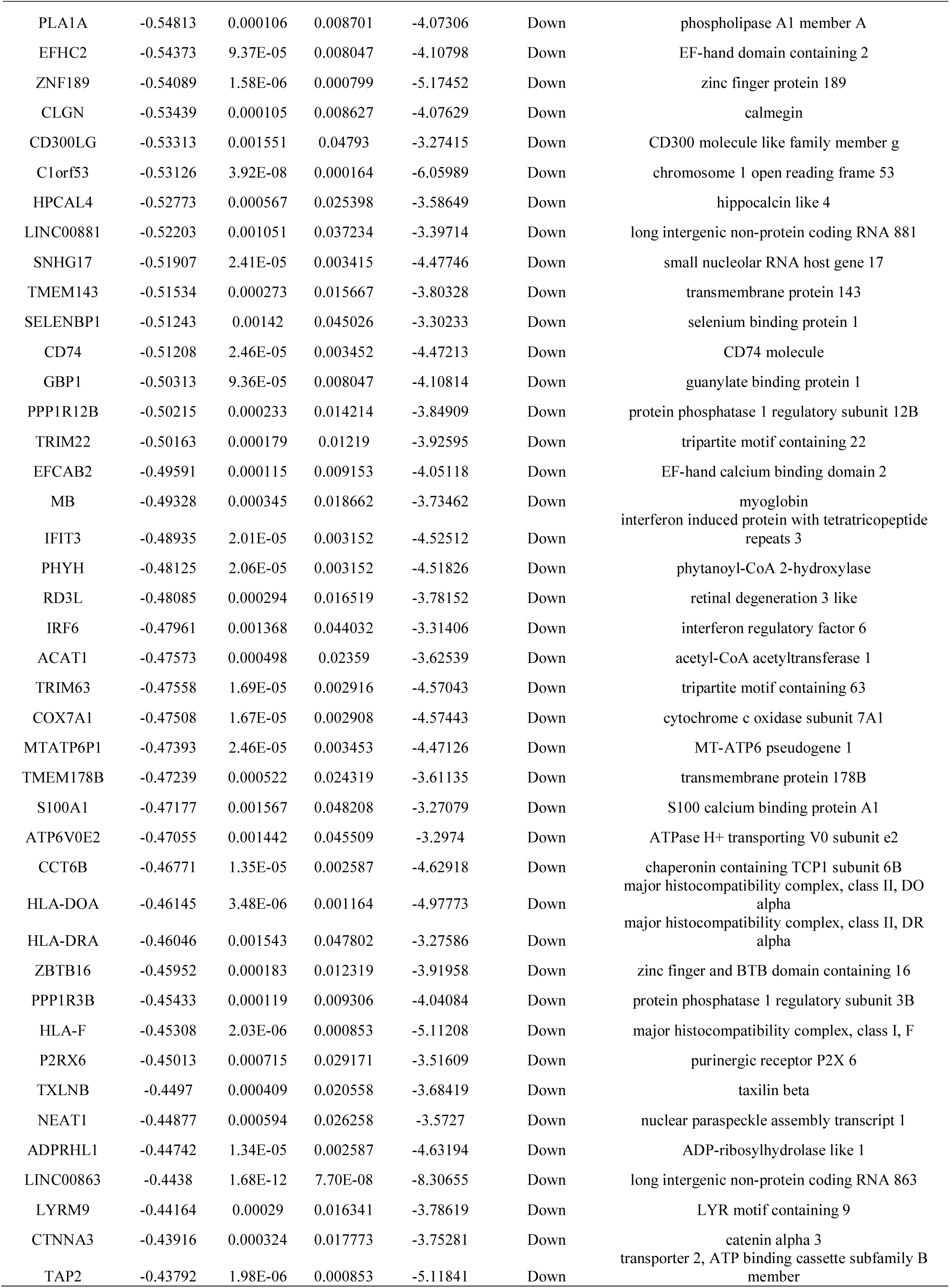
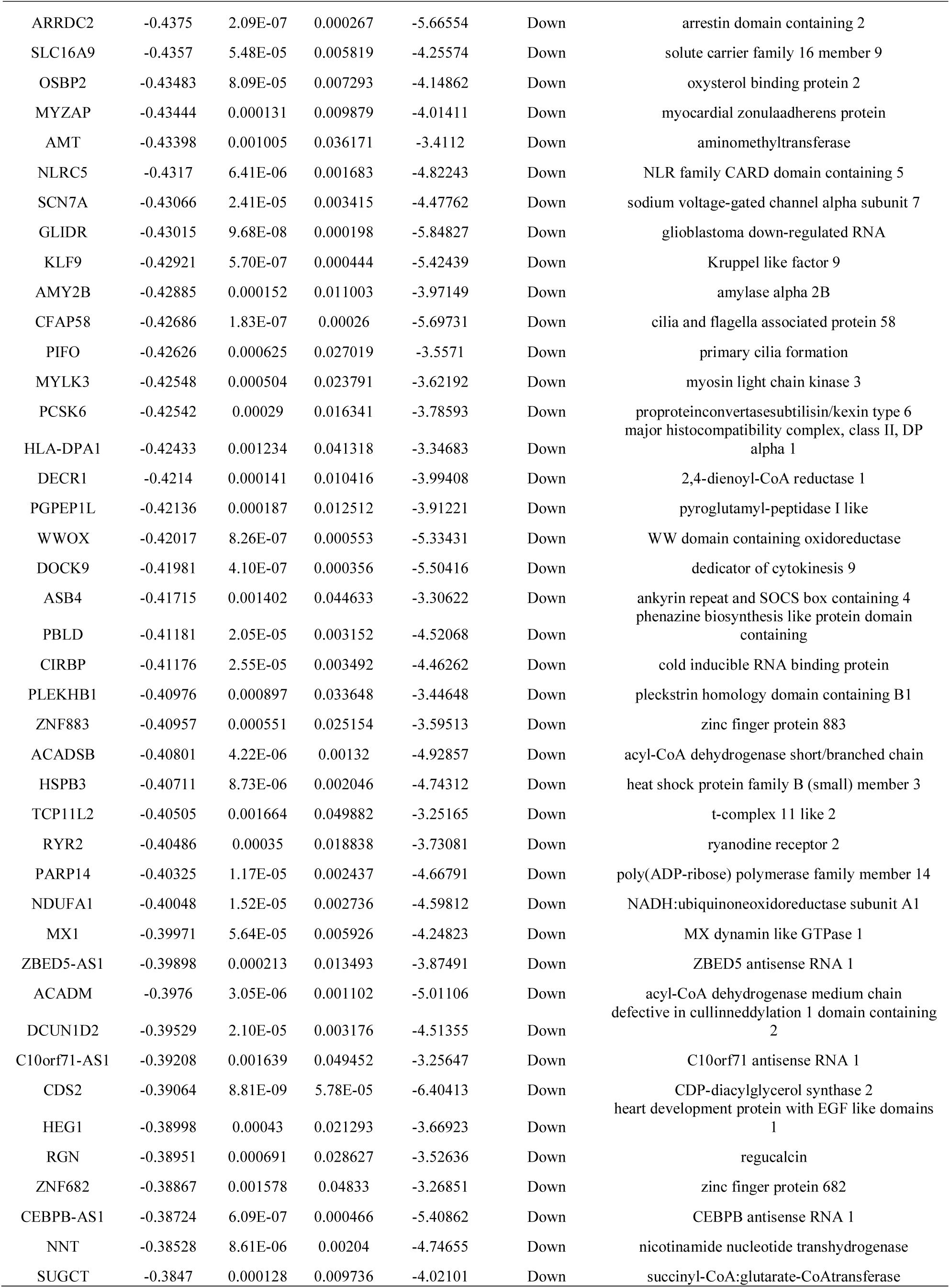
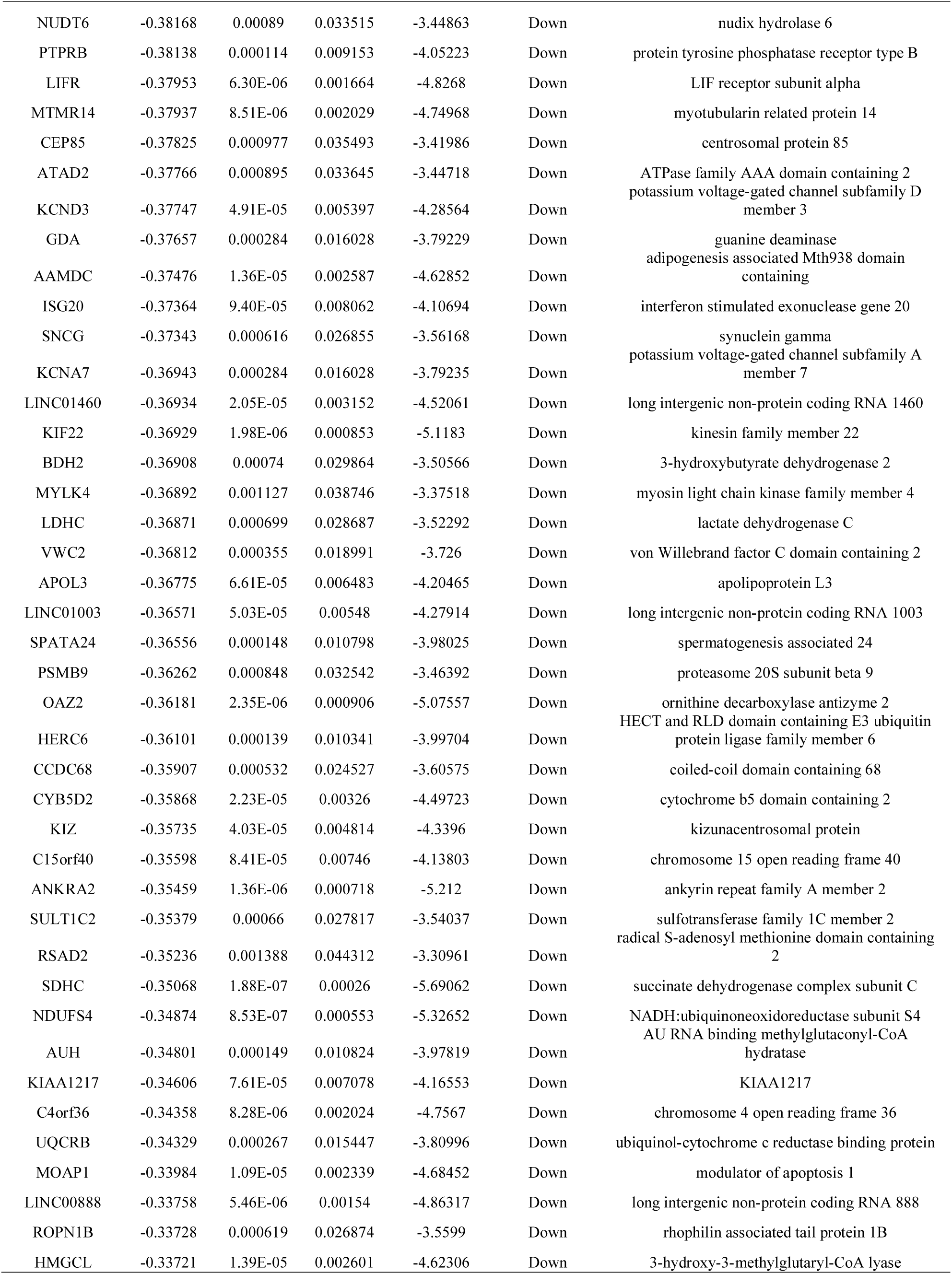
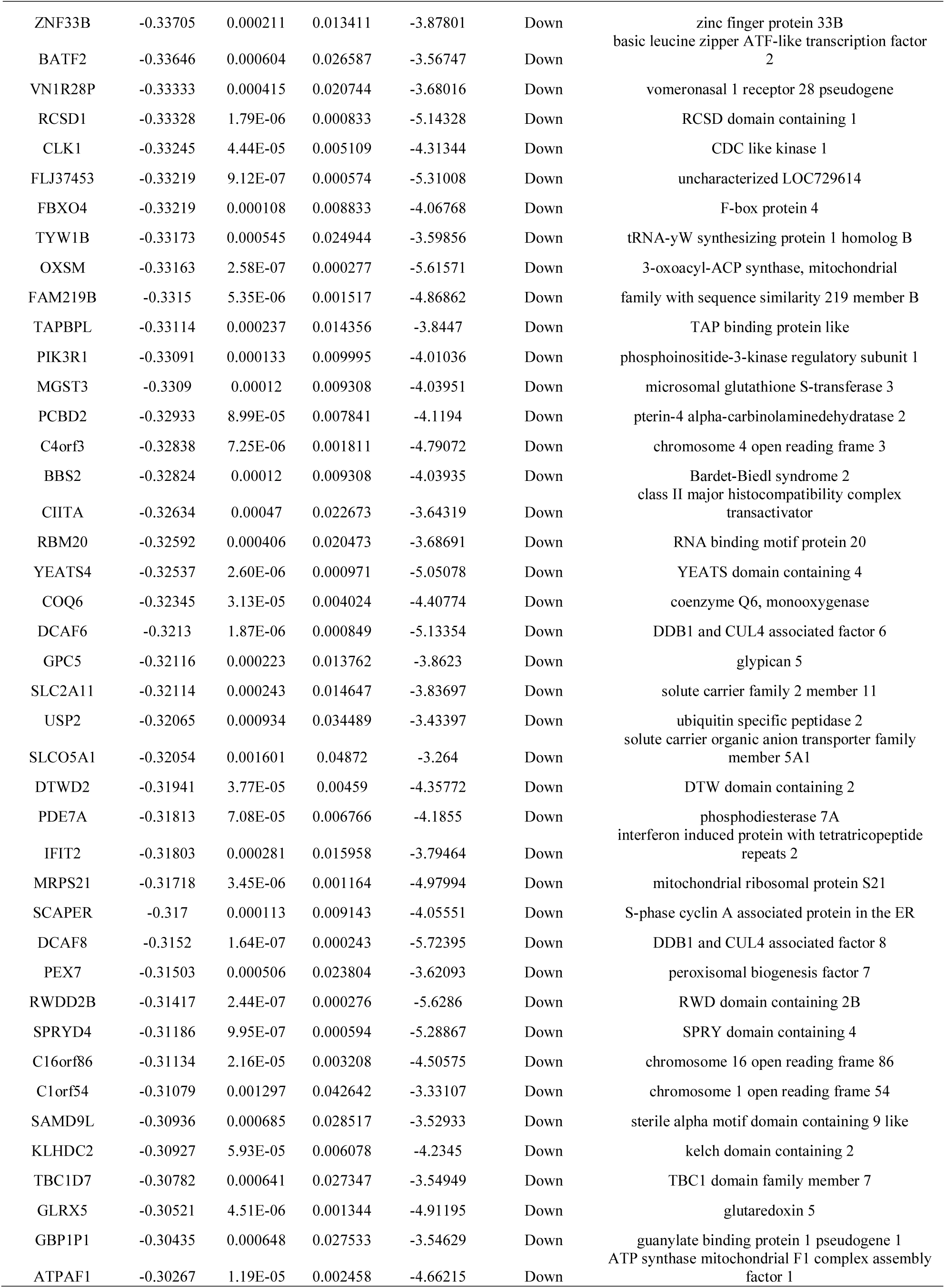
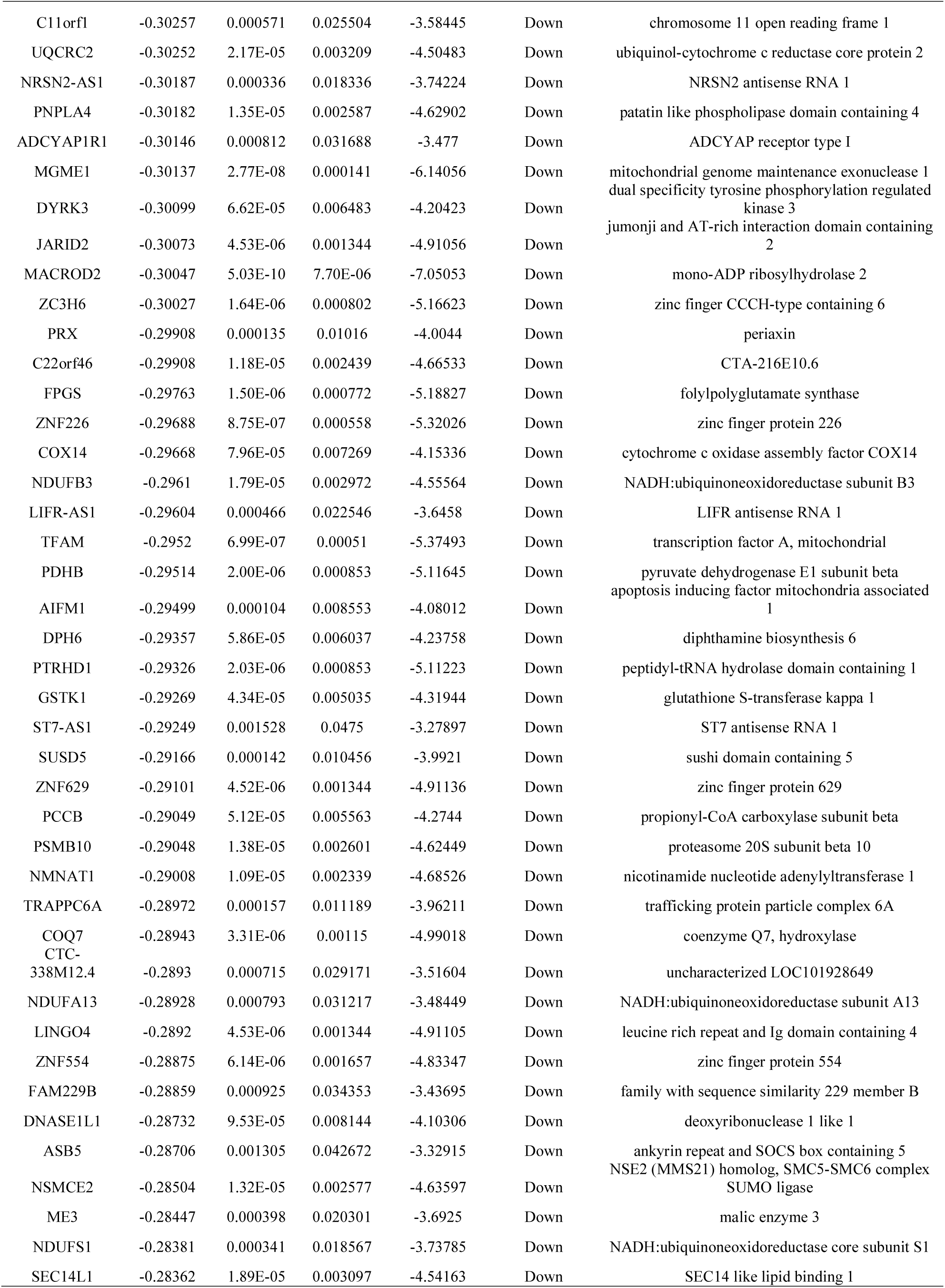
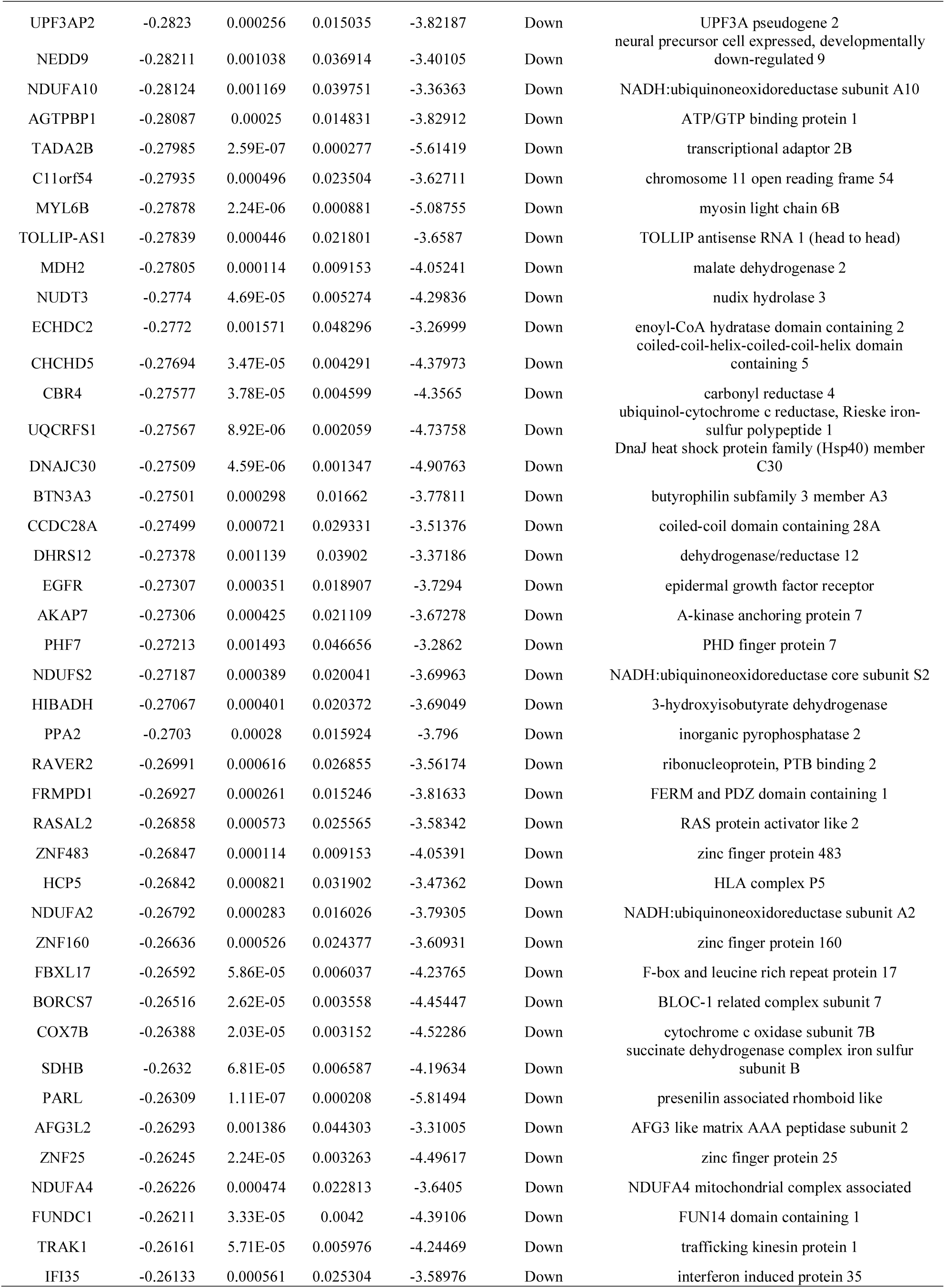
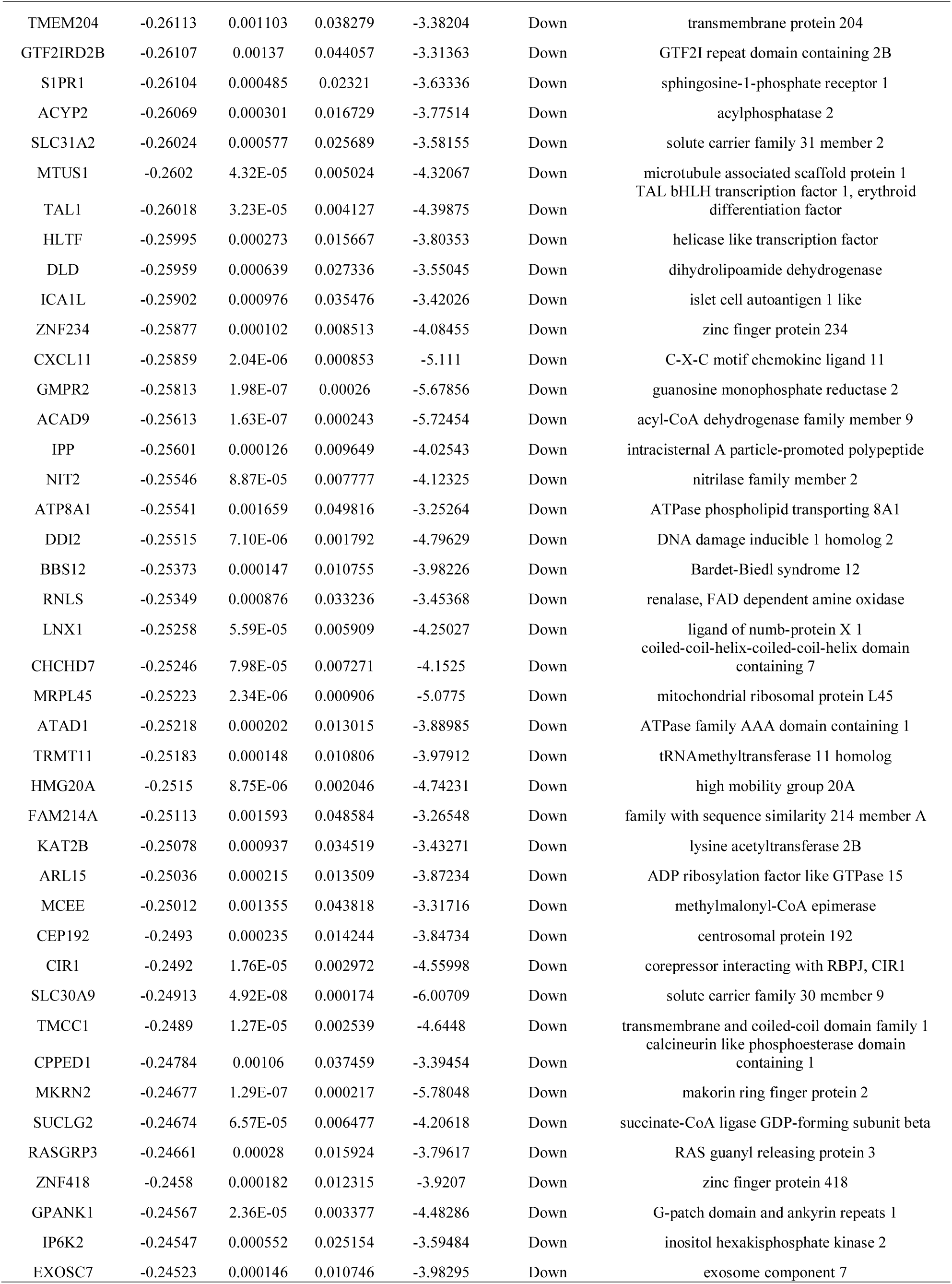
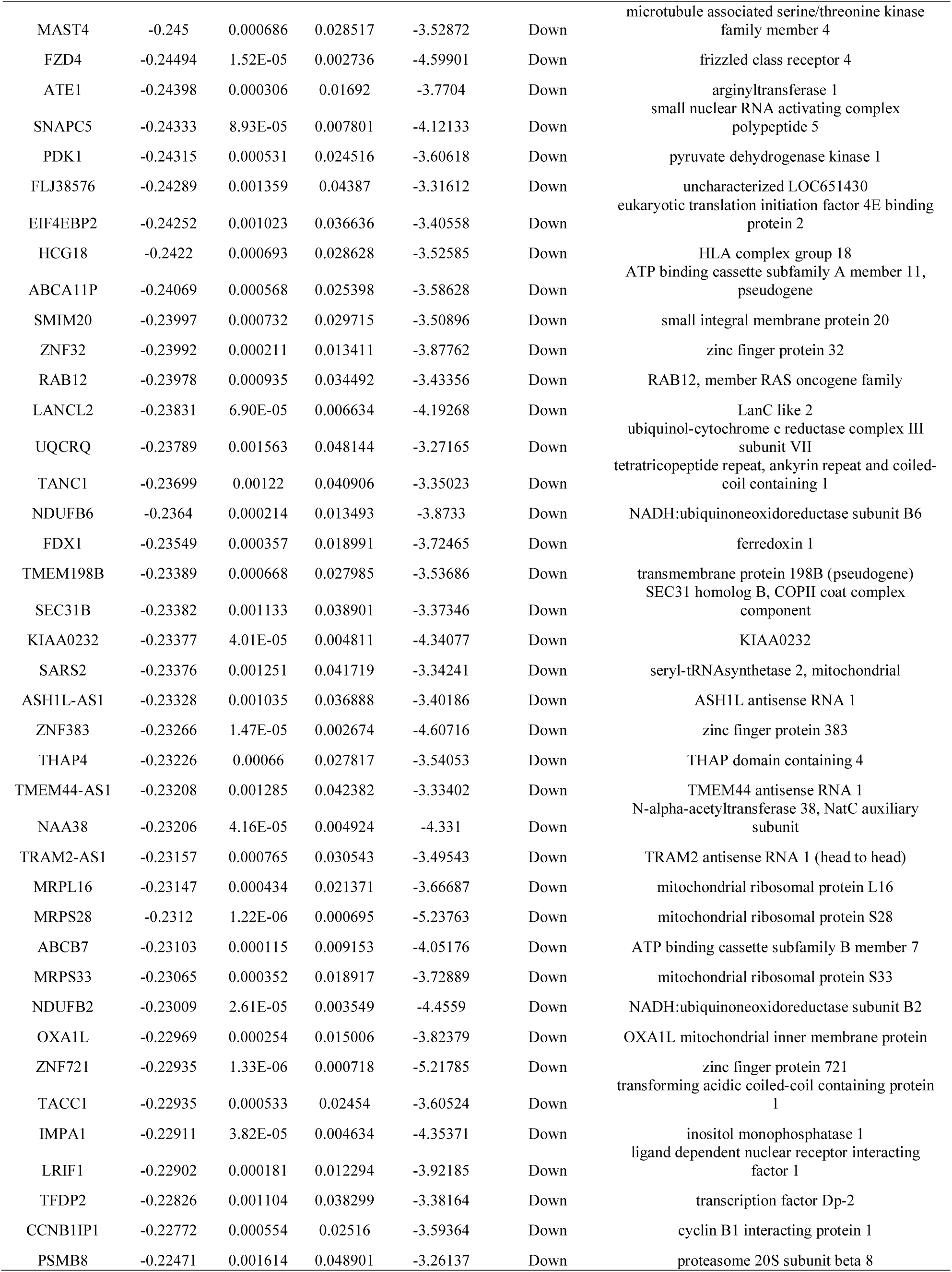
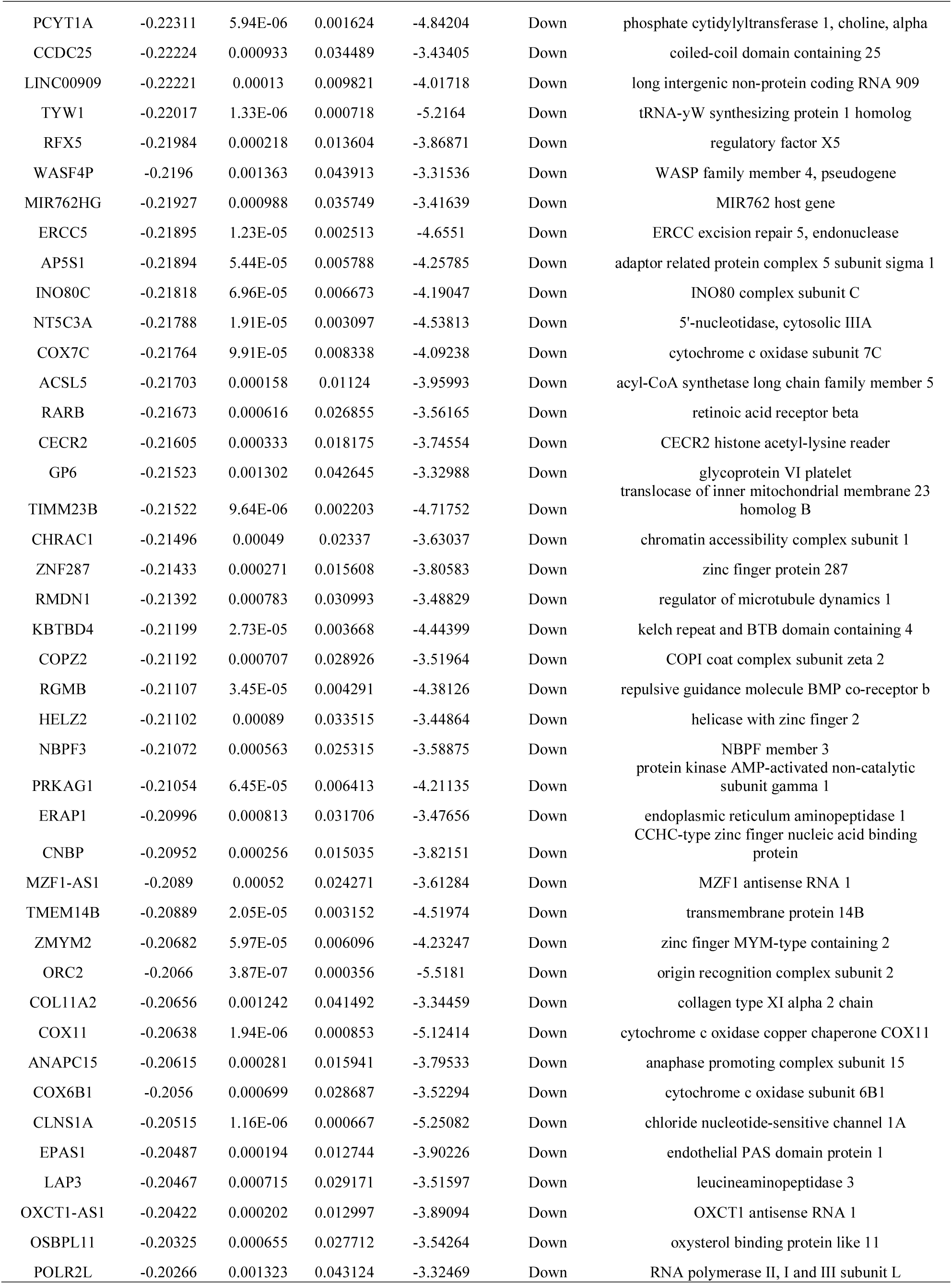
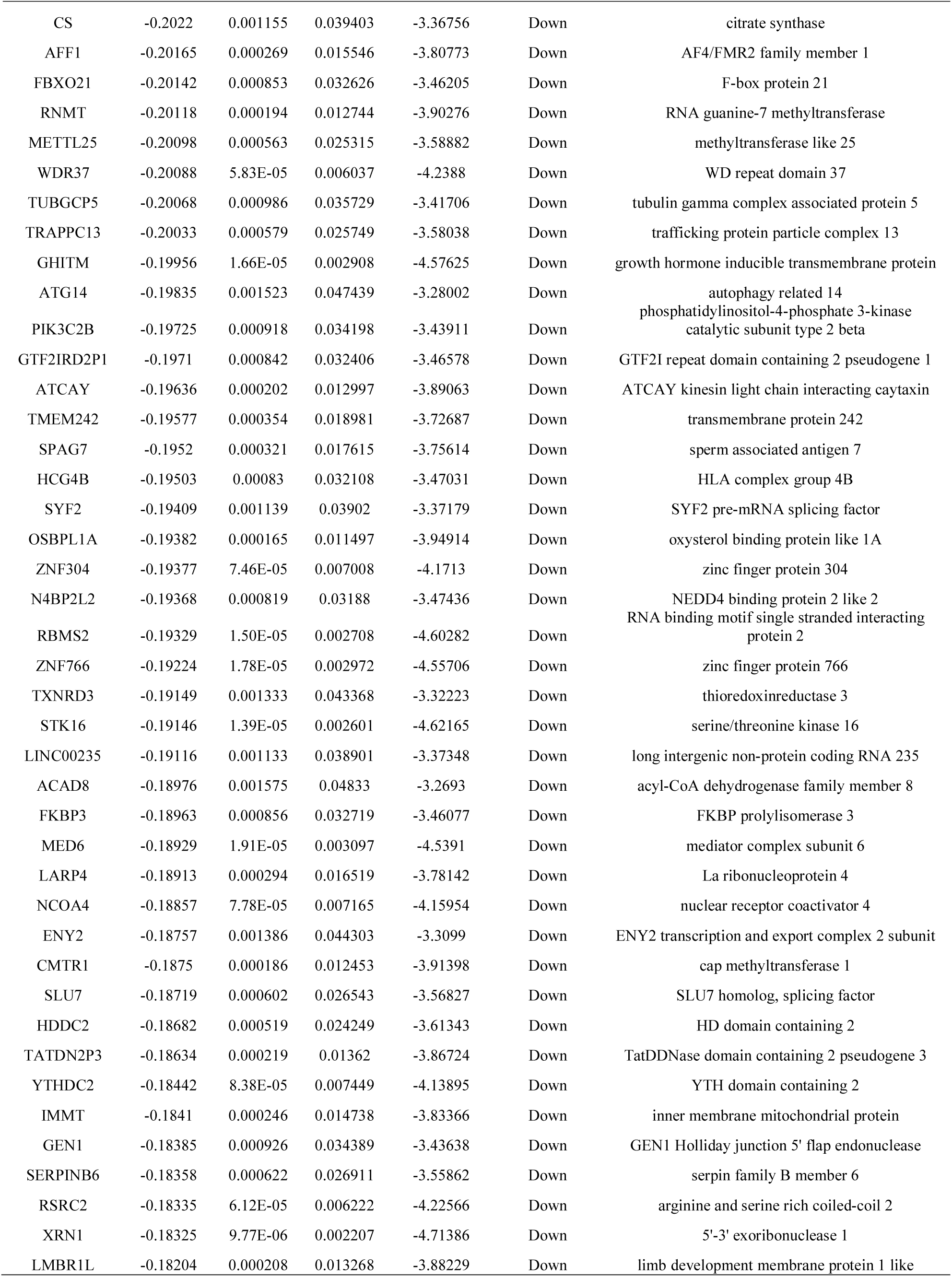
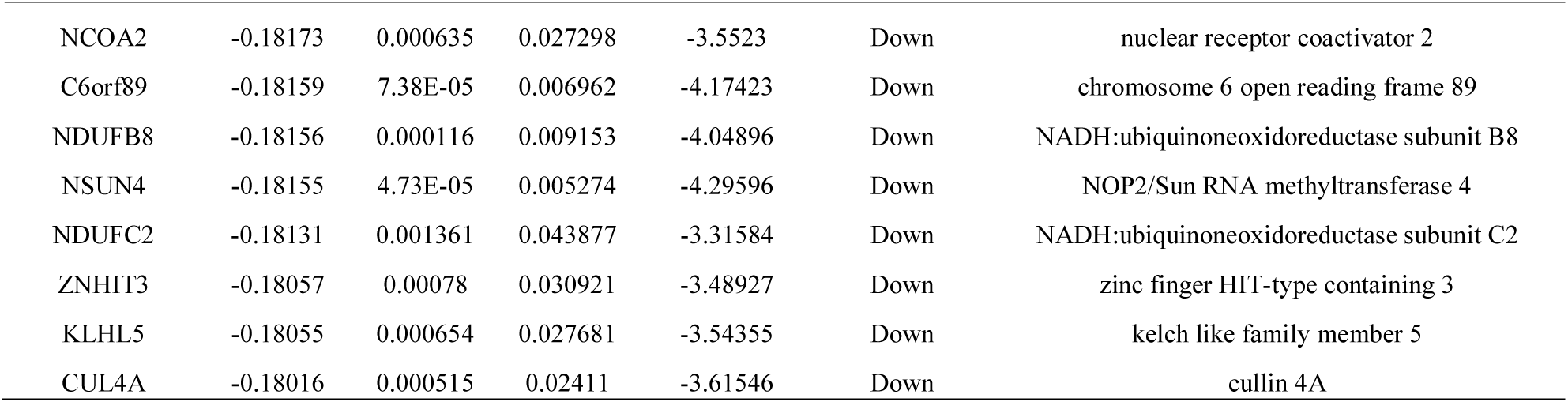
The statistical metrics for key differentially expressed genes (DEGs)

### GO and REACTOME pathway enrichment analysis of DEGs

The top 930 DEGs were chosen to perform GO term and REACTOME pathway analyses. We detected enrichment in several BP GO terms such as localization, organic substance transport, small molecule metabolic process and cellular metabolic process and are listed in Table 2. In terms of CC, cytoplasm, membrane, intracellular anatomical structure and organelle lumen and are listed in Table 2. What’s more, some MF GO terms, such as protein binding, enzyme binding, catalytic activity and nucleoside phosphate binding, and are listed in Table 2. As to REACTOME pathway enrichment analysis, SARS-CoV infections, asparagine N-linked glycosylation, the citric acid (TCA) cycle and respiratory electron transport, and respiratory electron transport were mostly associated with these genes and are listed in Table 3.

**Table 2.**
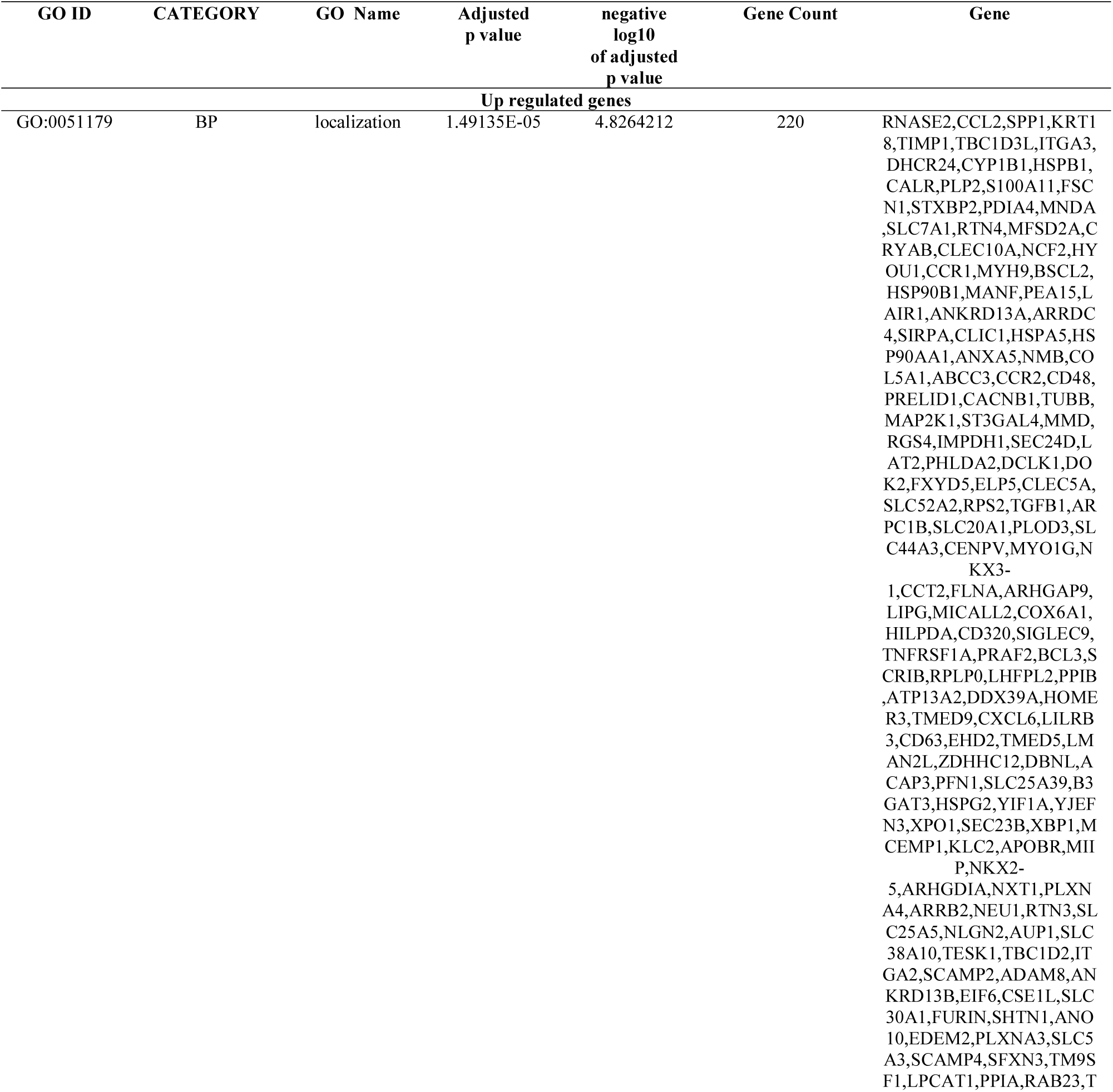
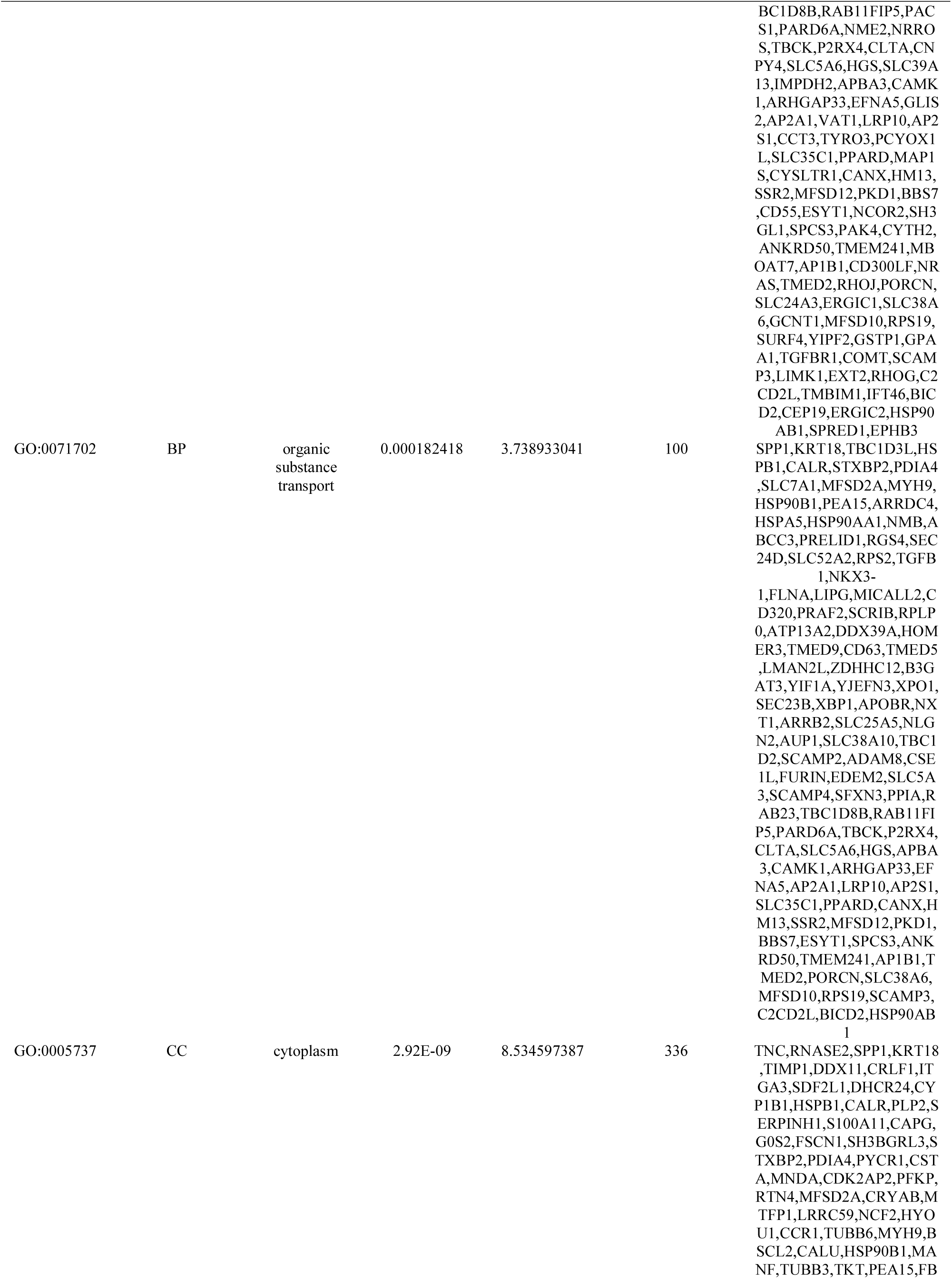
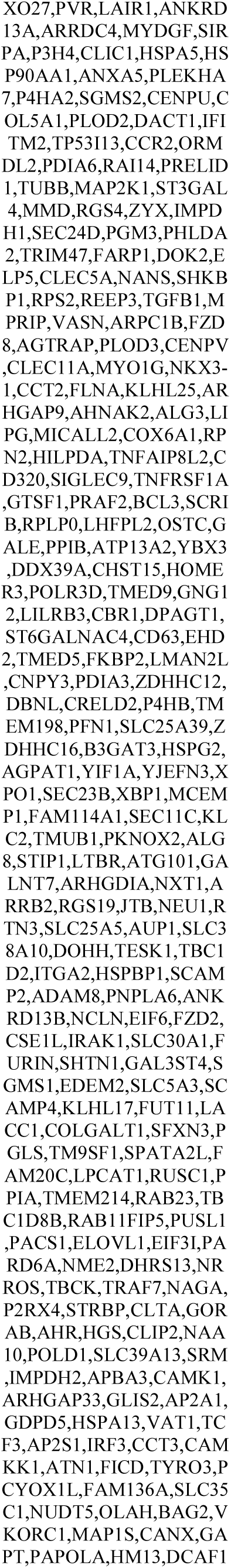
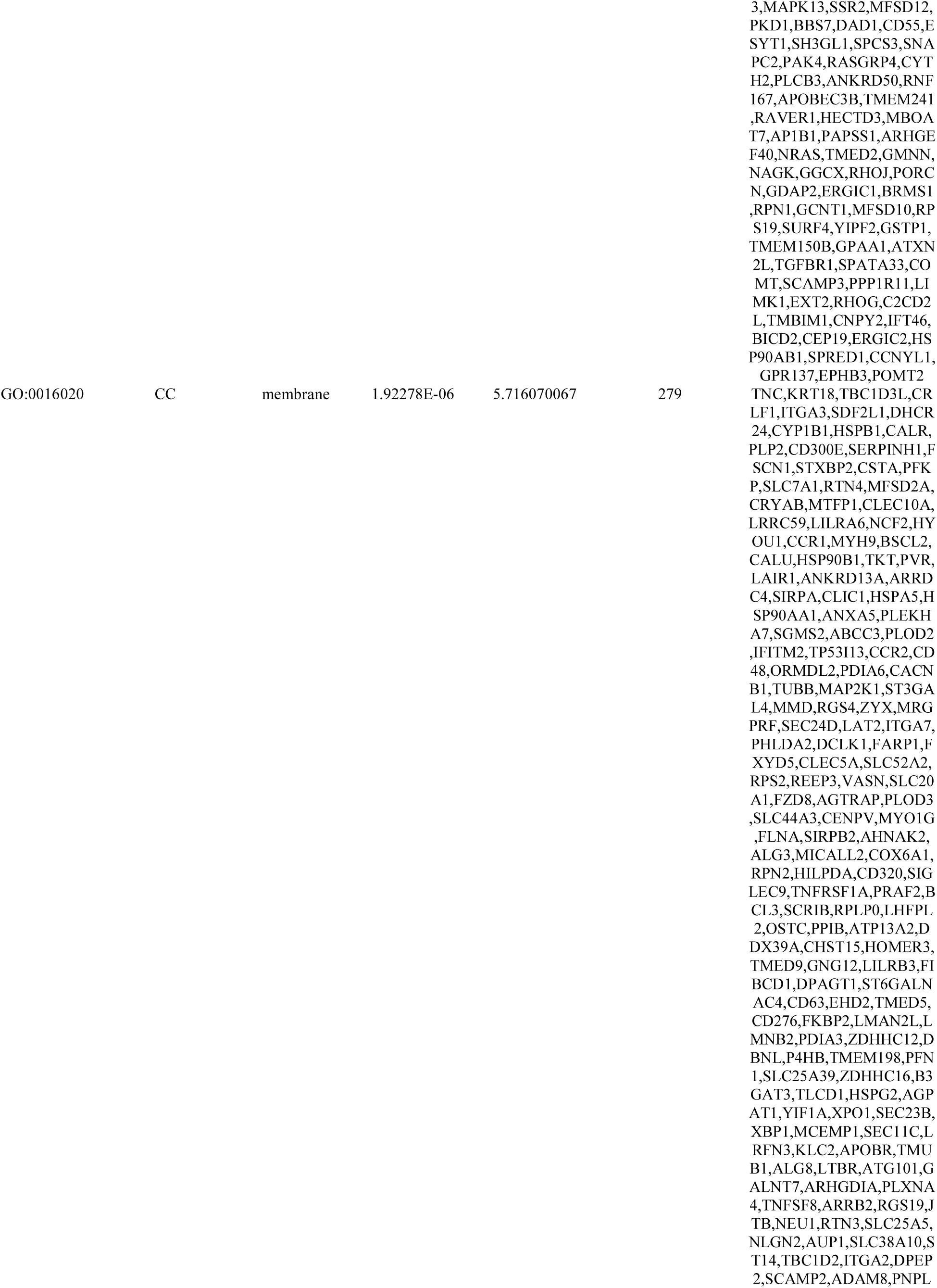
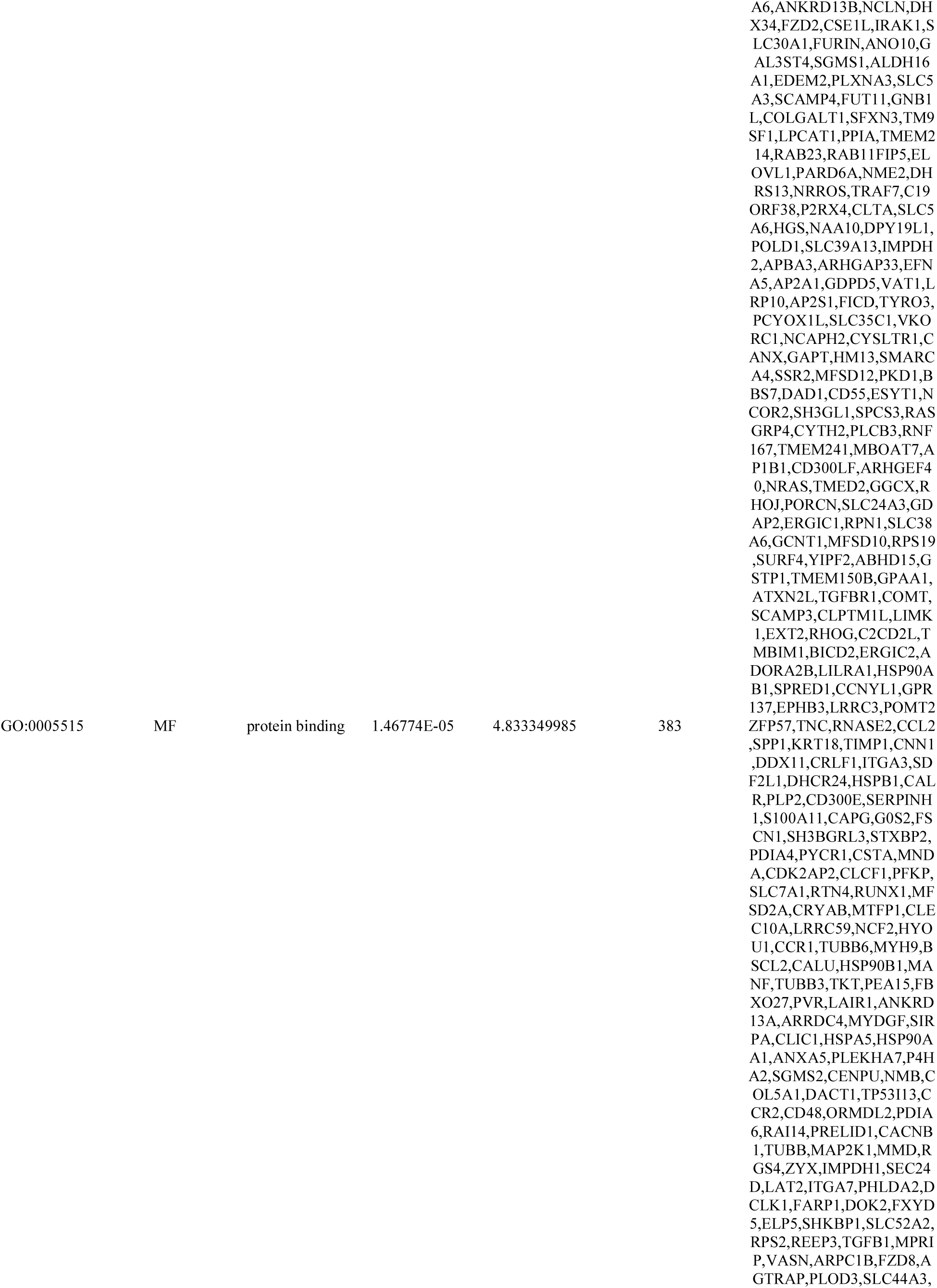
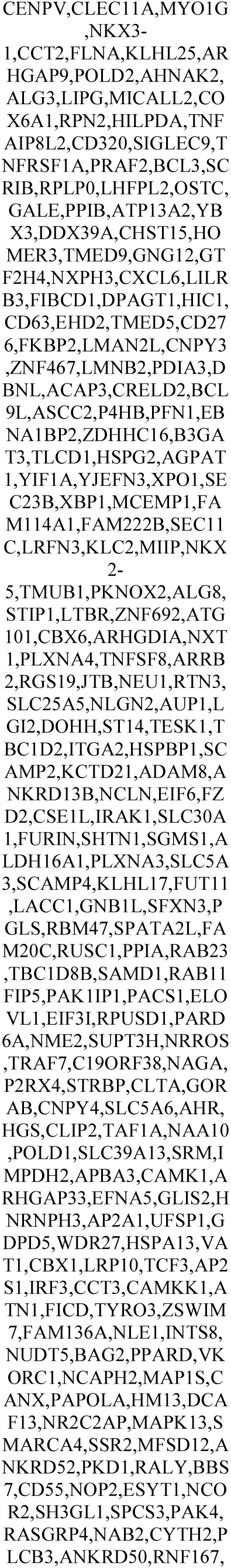
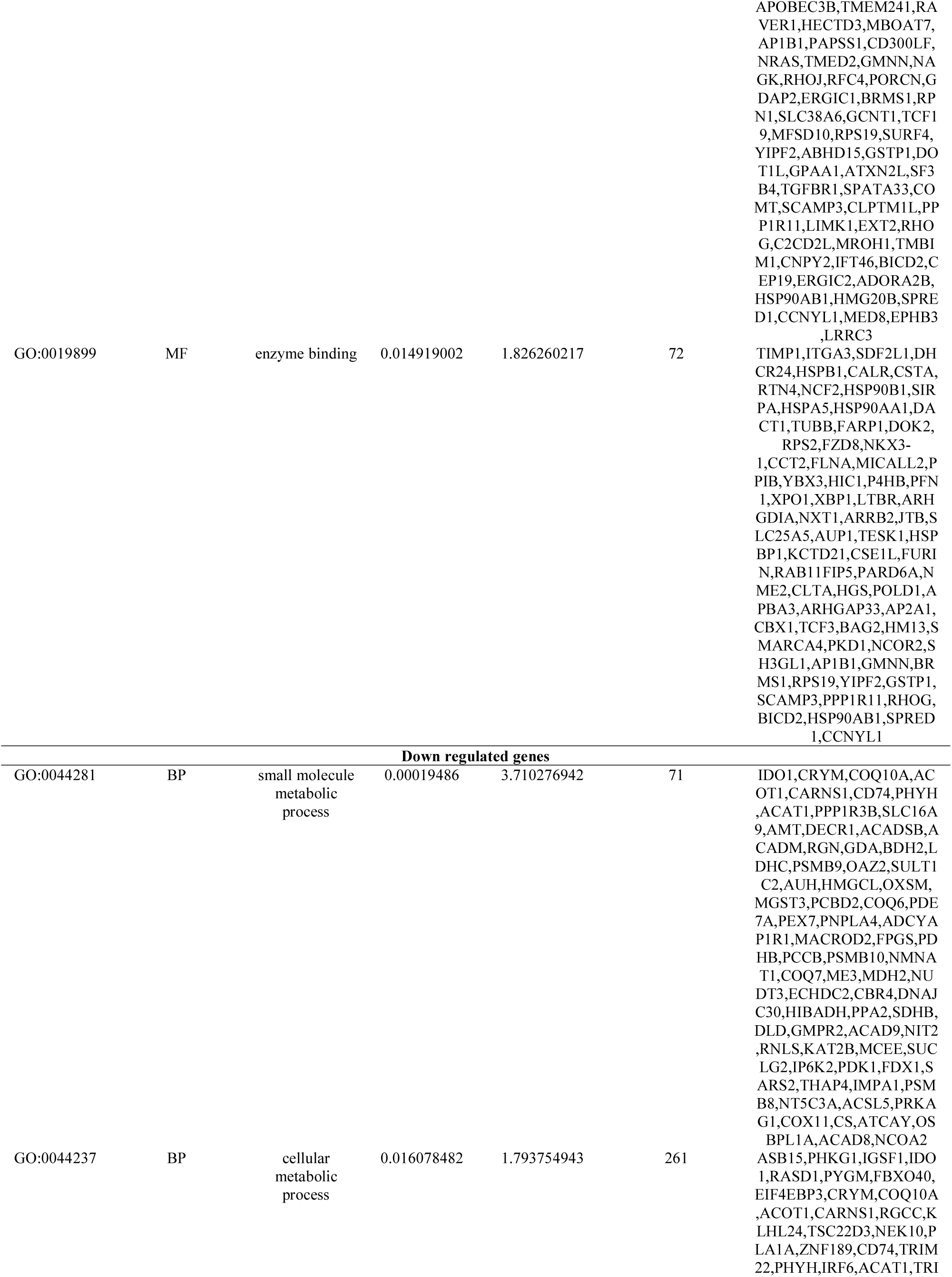
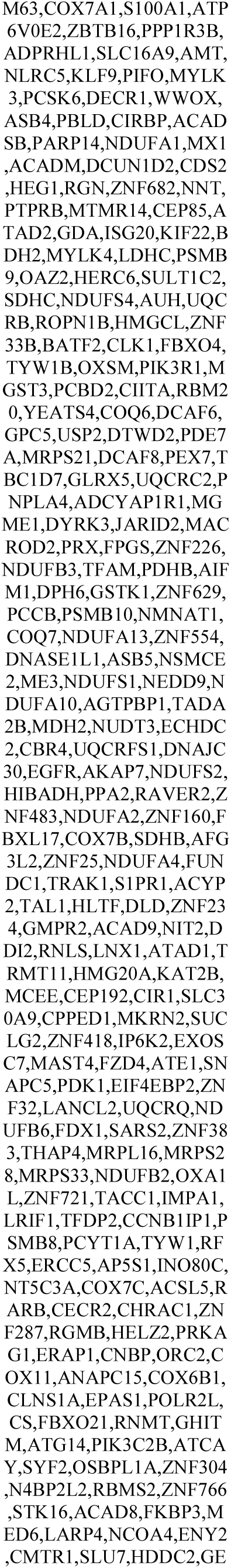
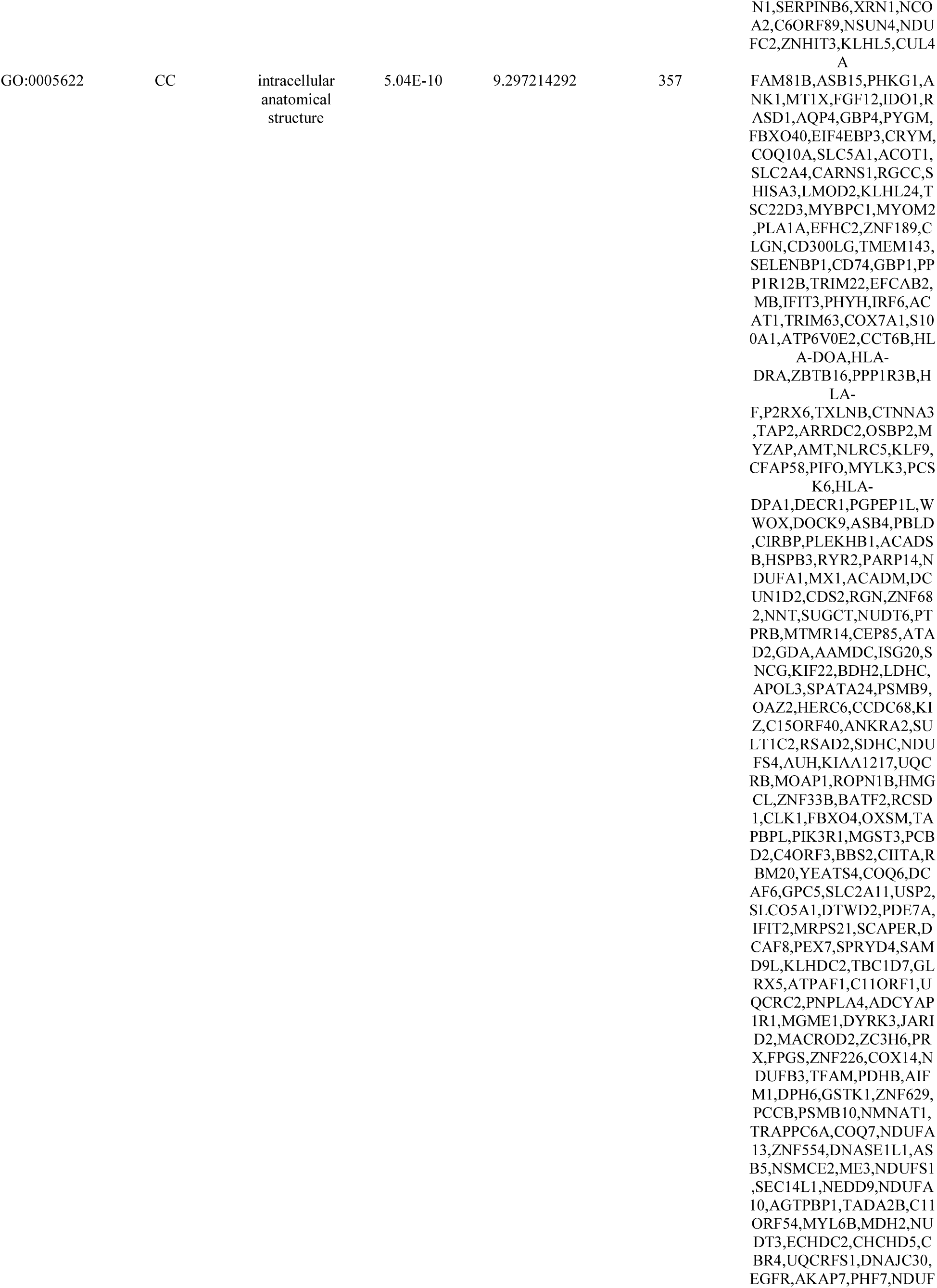
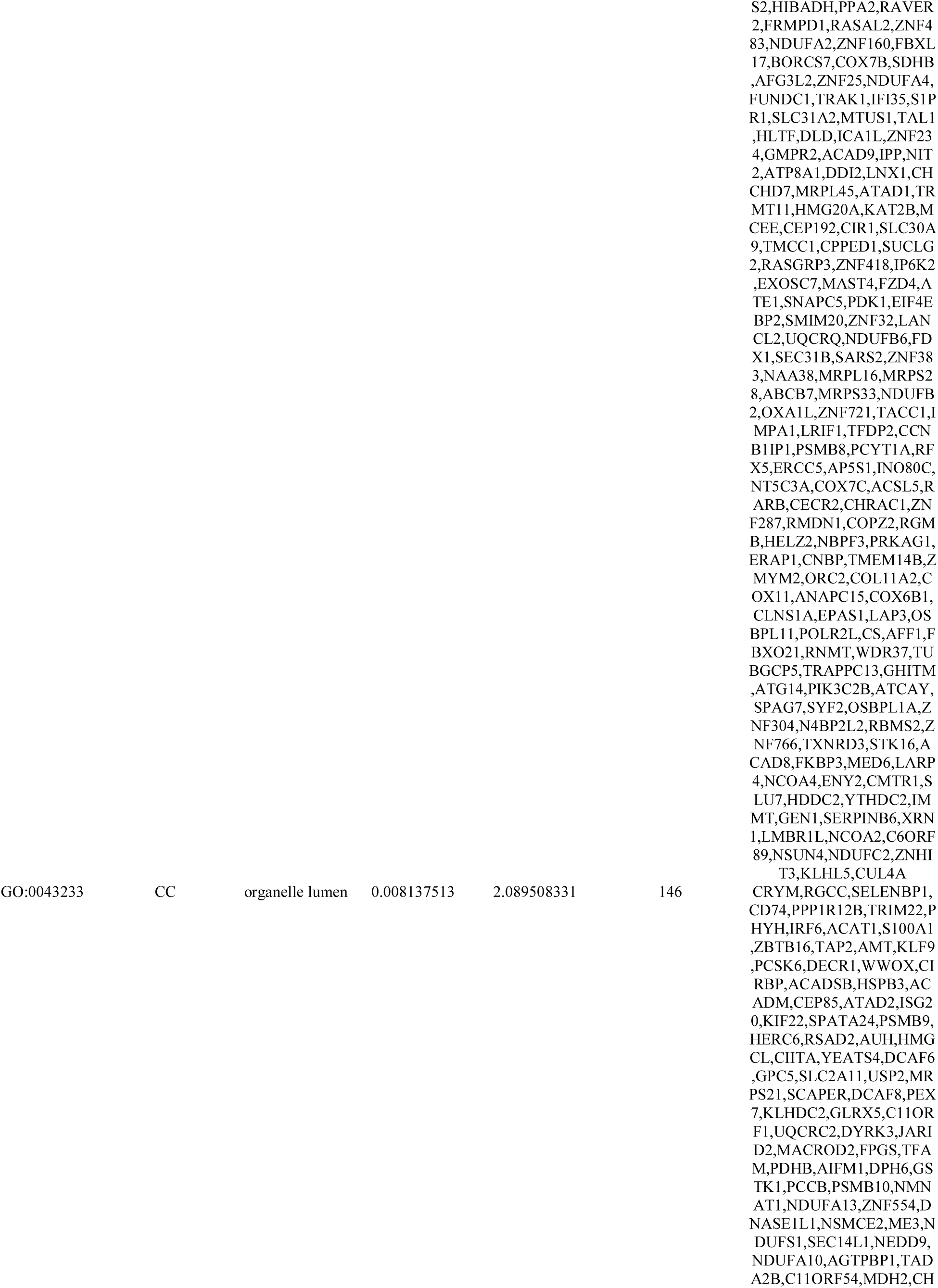
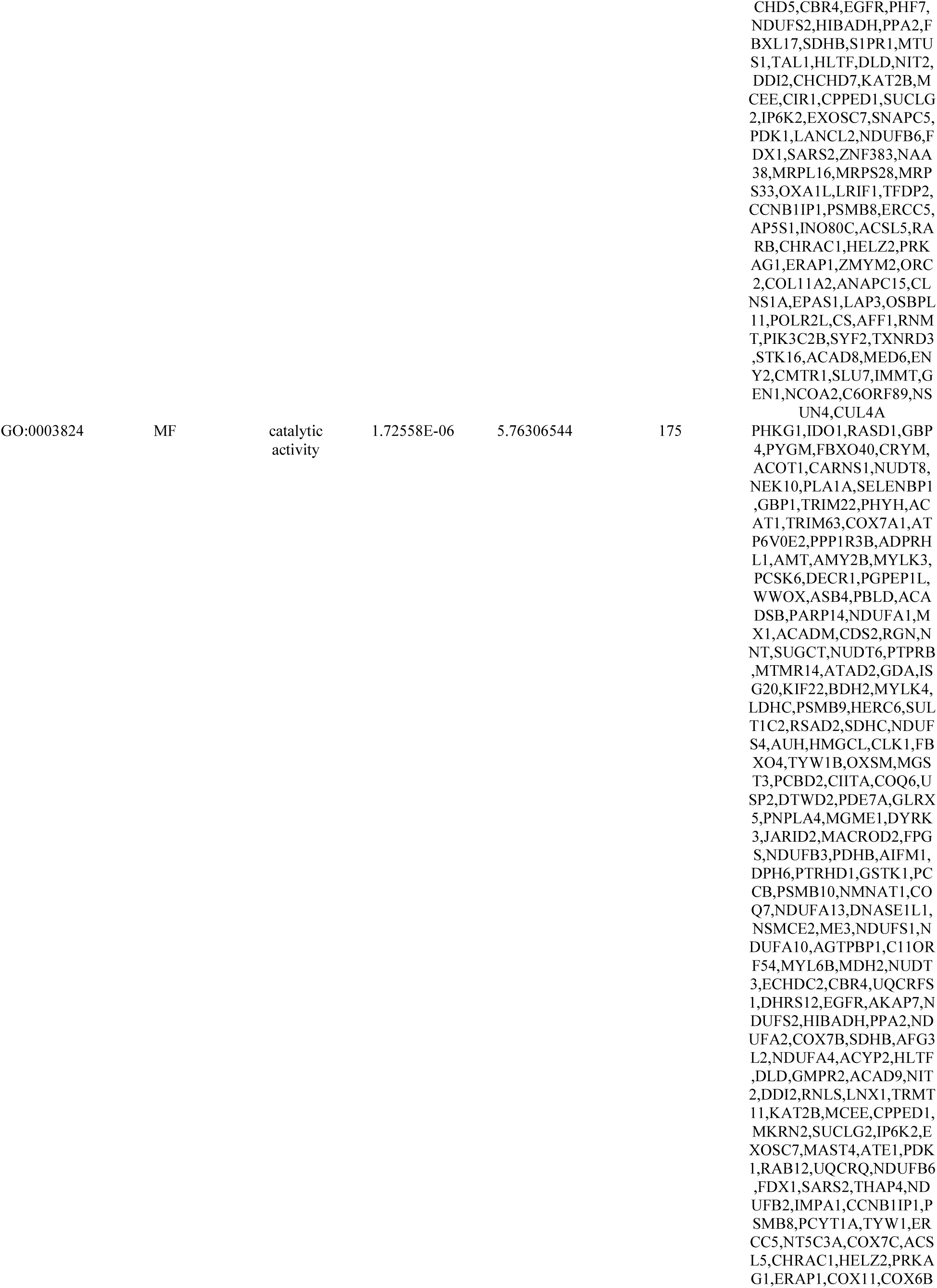
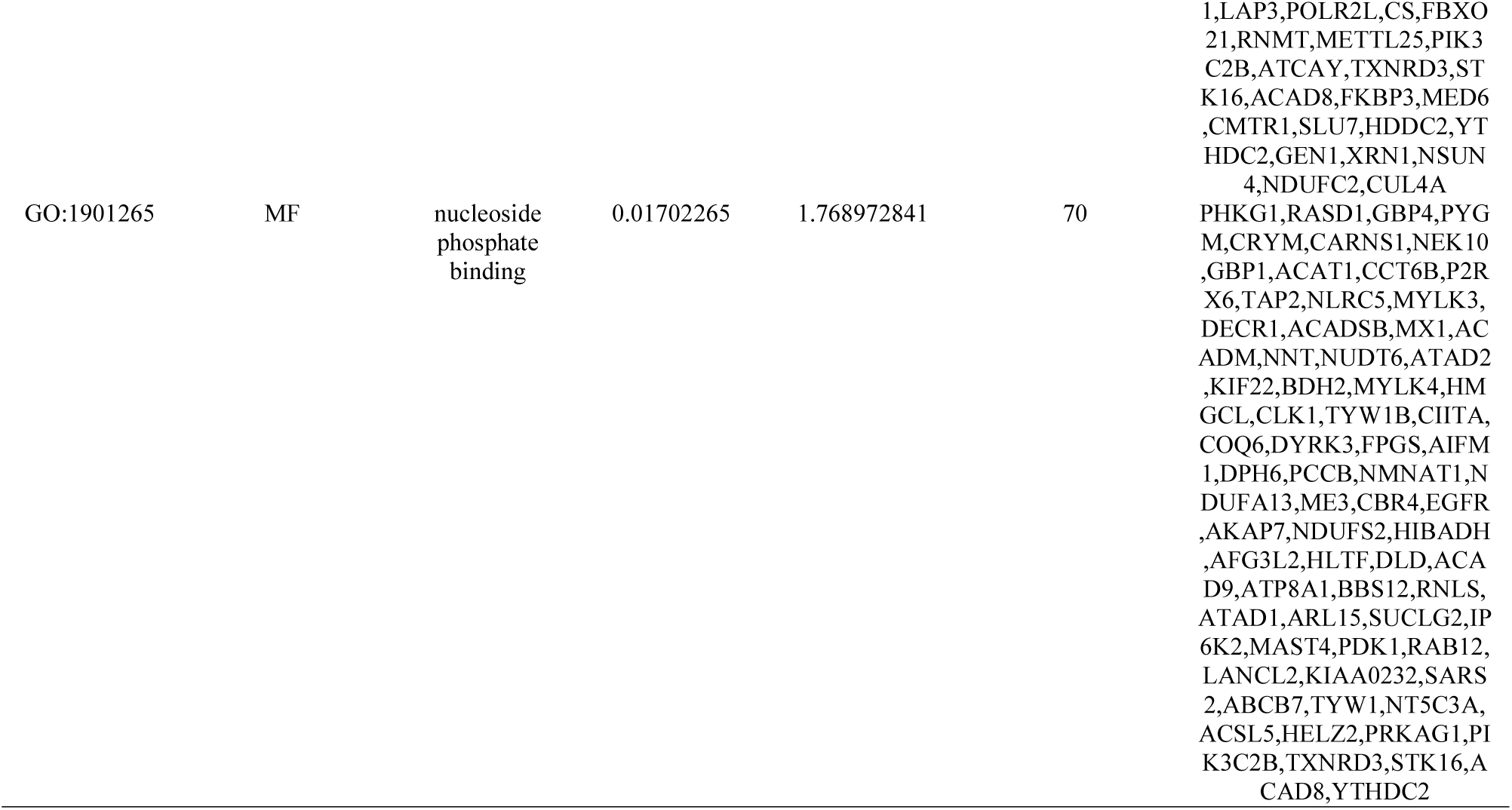
The enriched GO terms of the up and down regulated differentially expressed genes

**Table 3.**
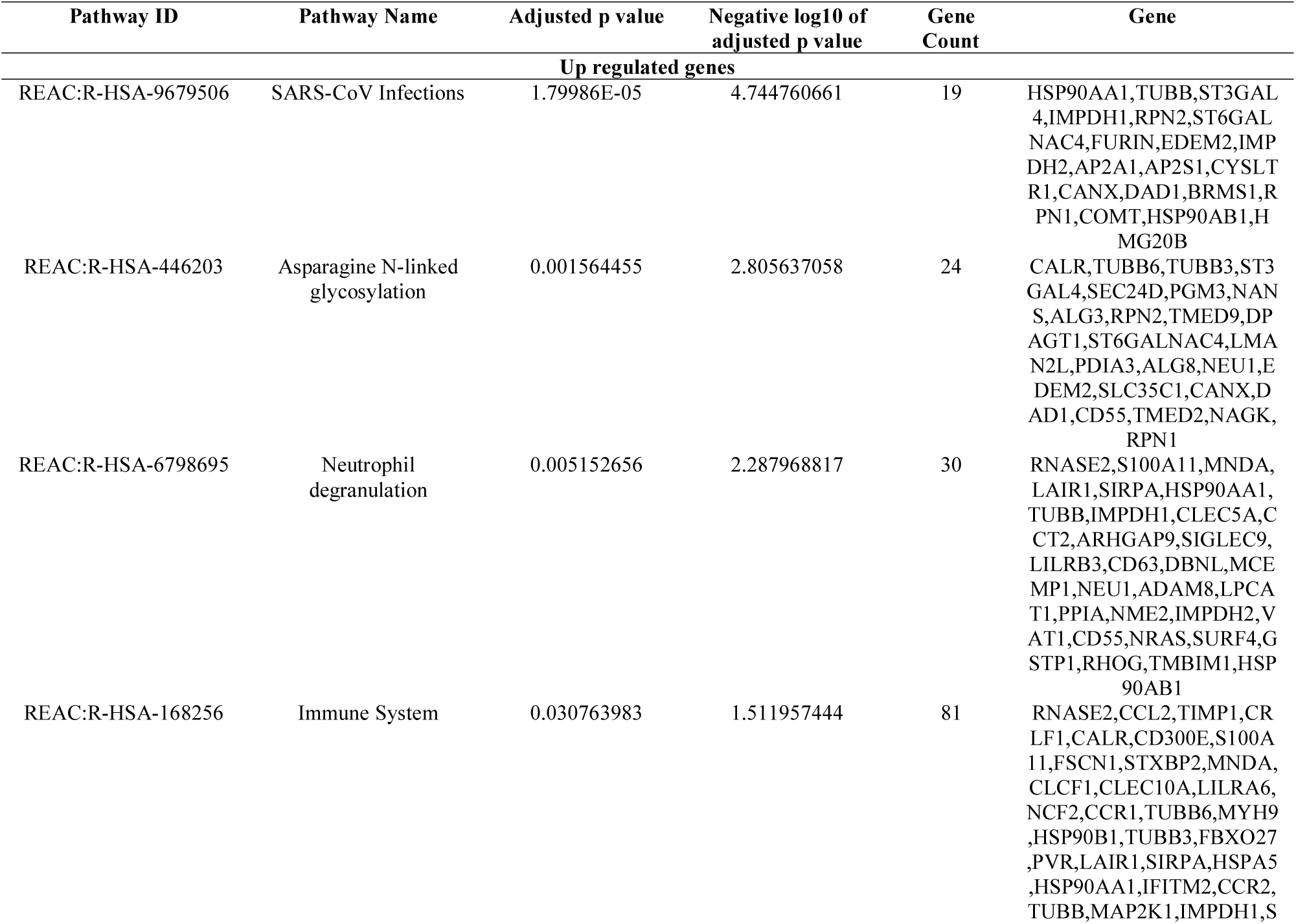
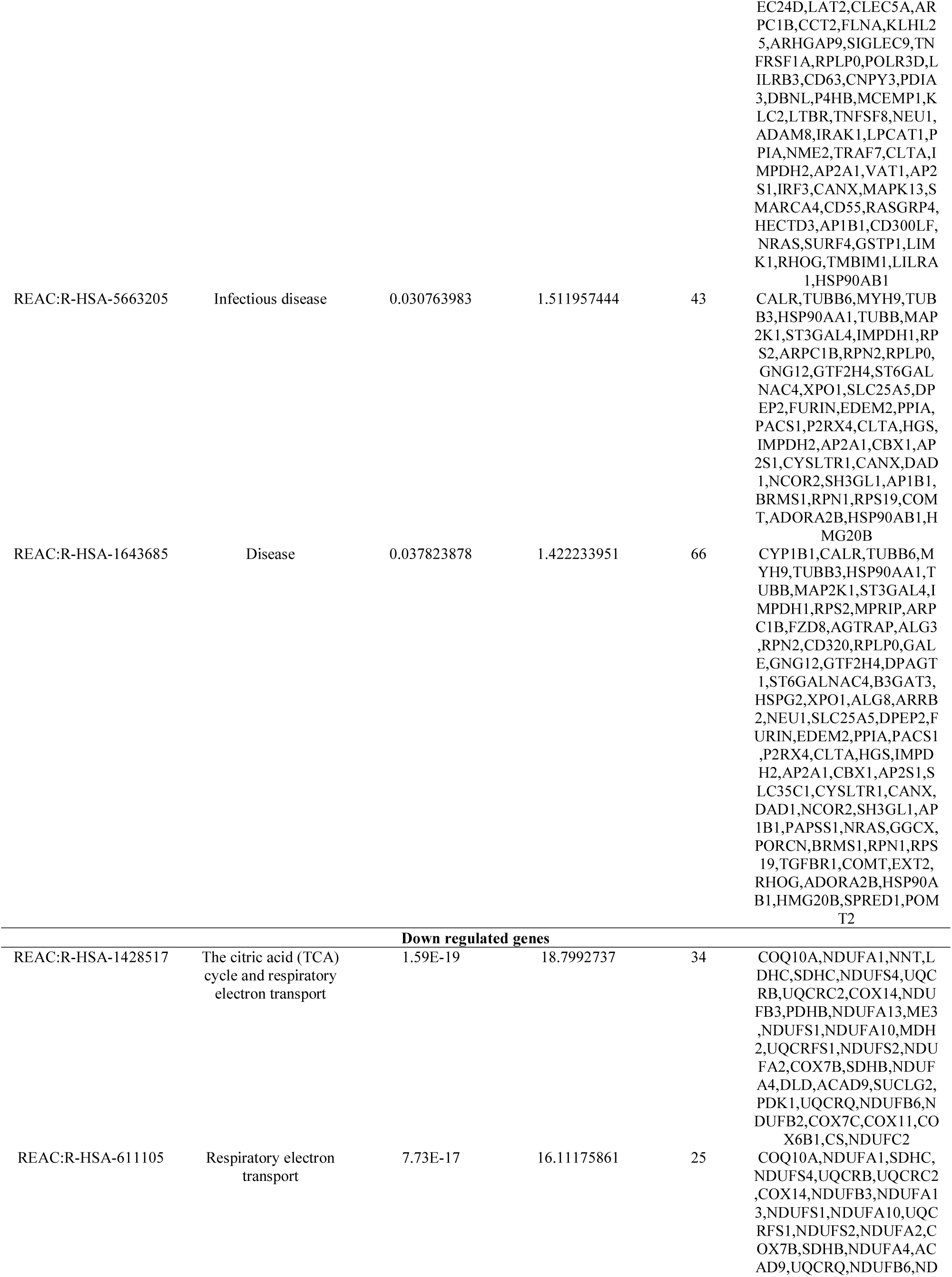
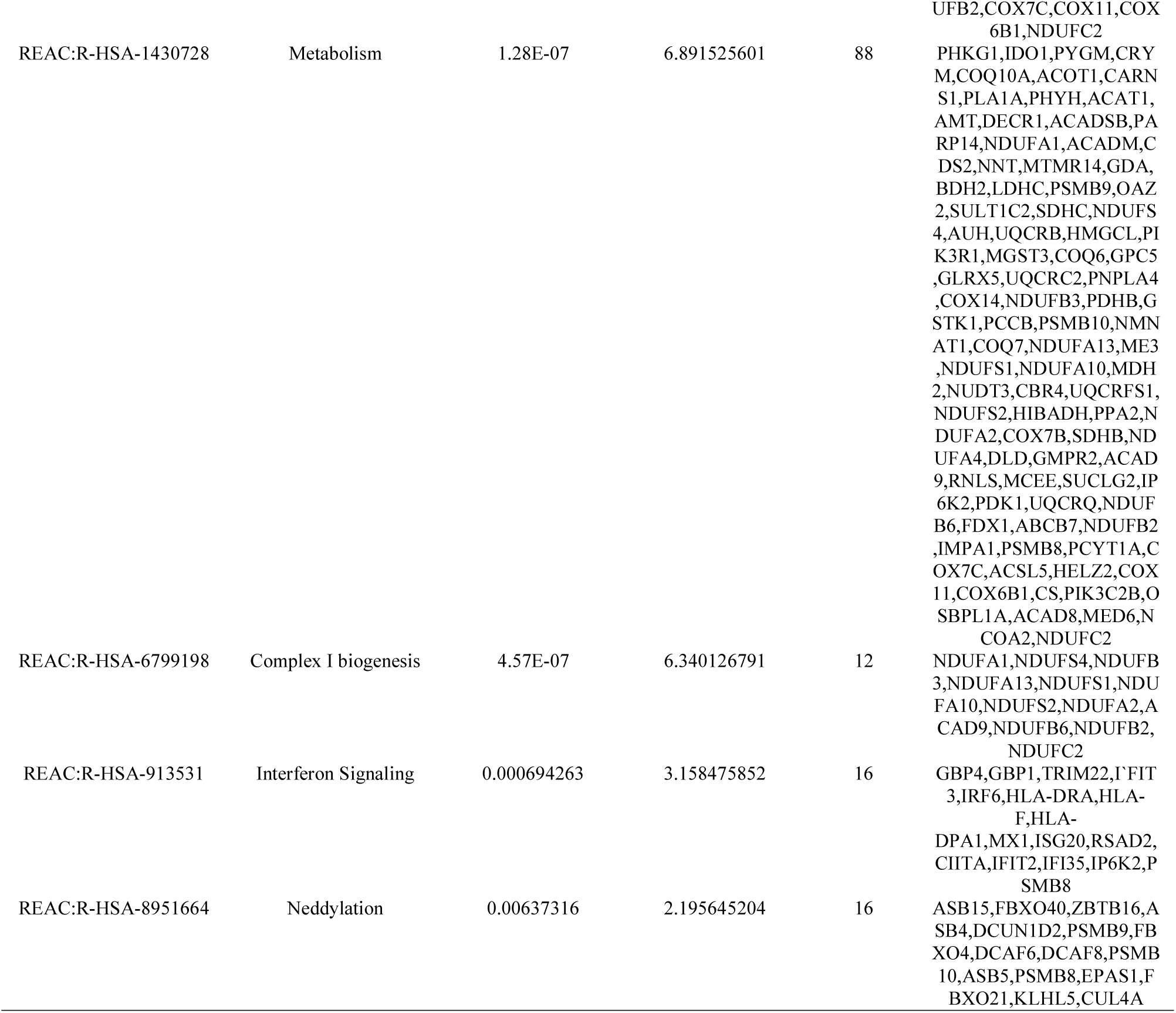
The enriched pathway terms of the up and down regulated differentially expressed genes

**Table 5.**
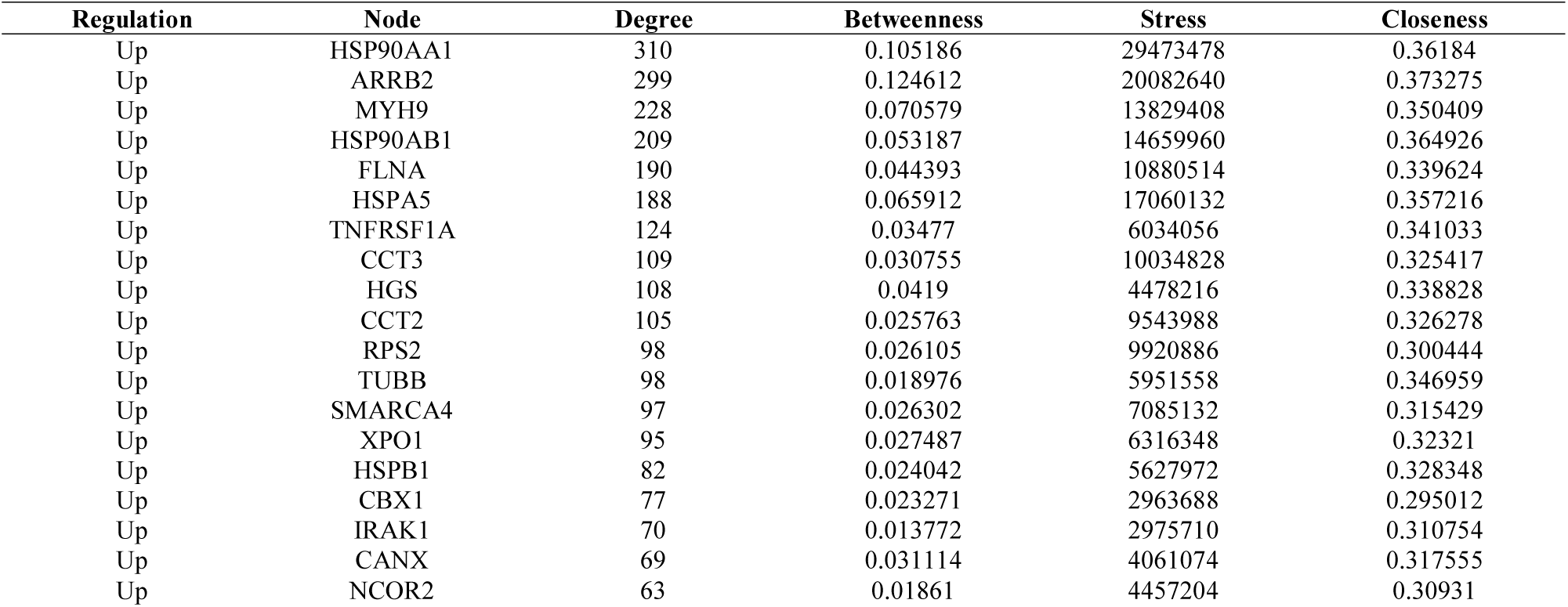
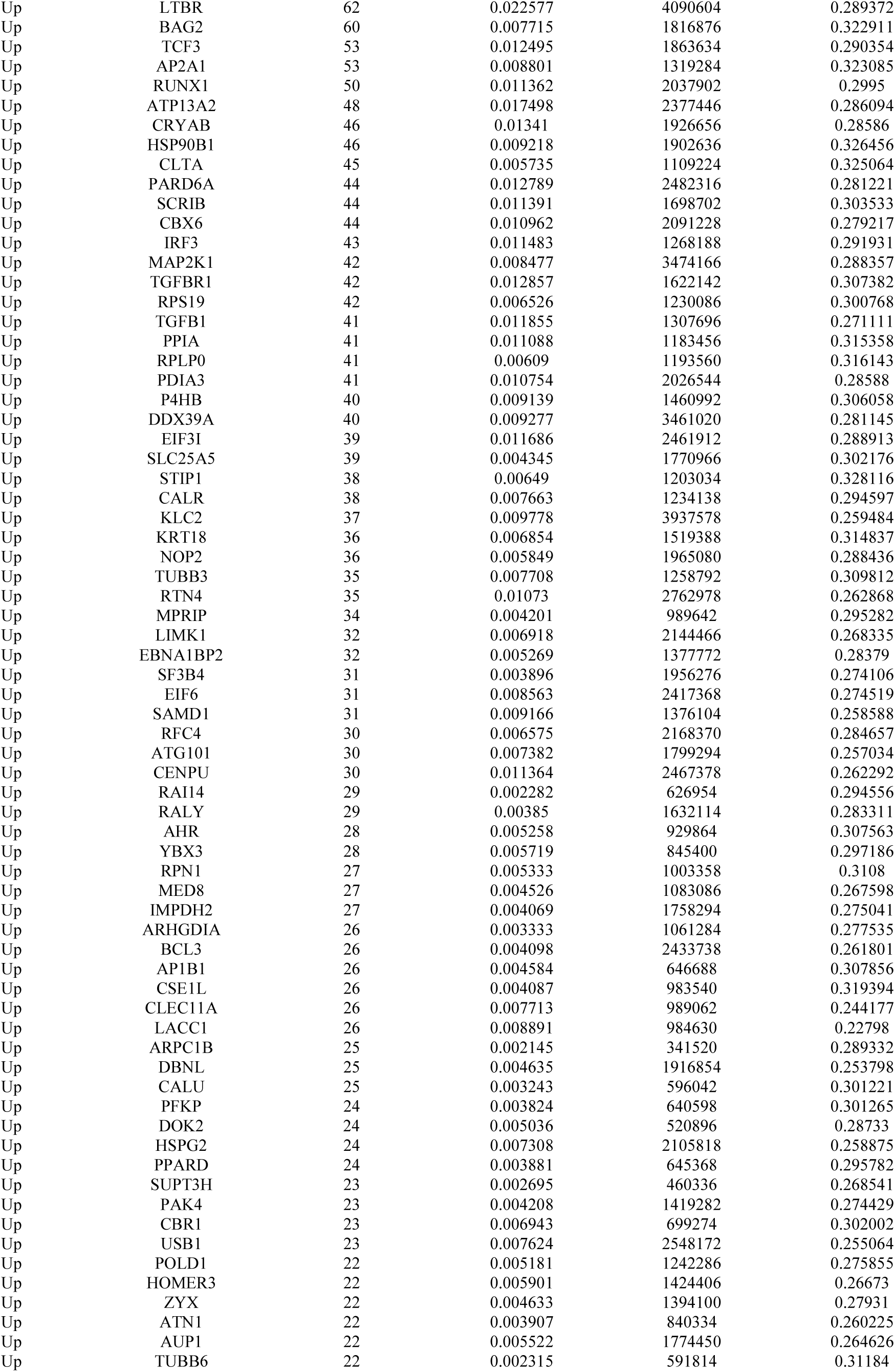
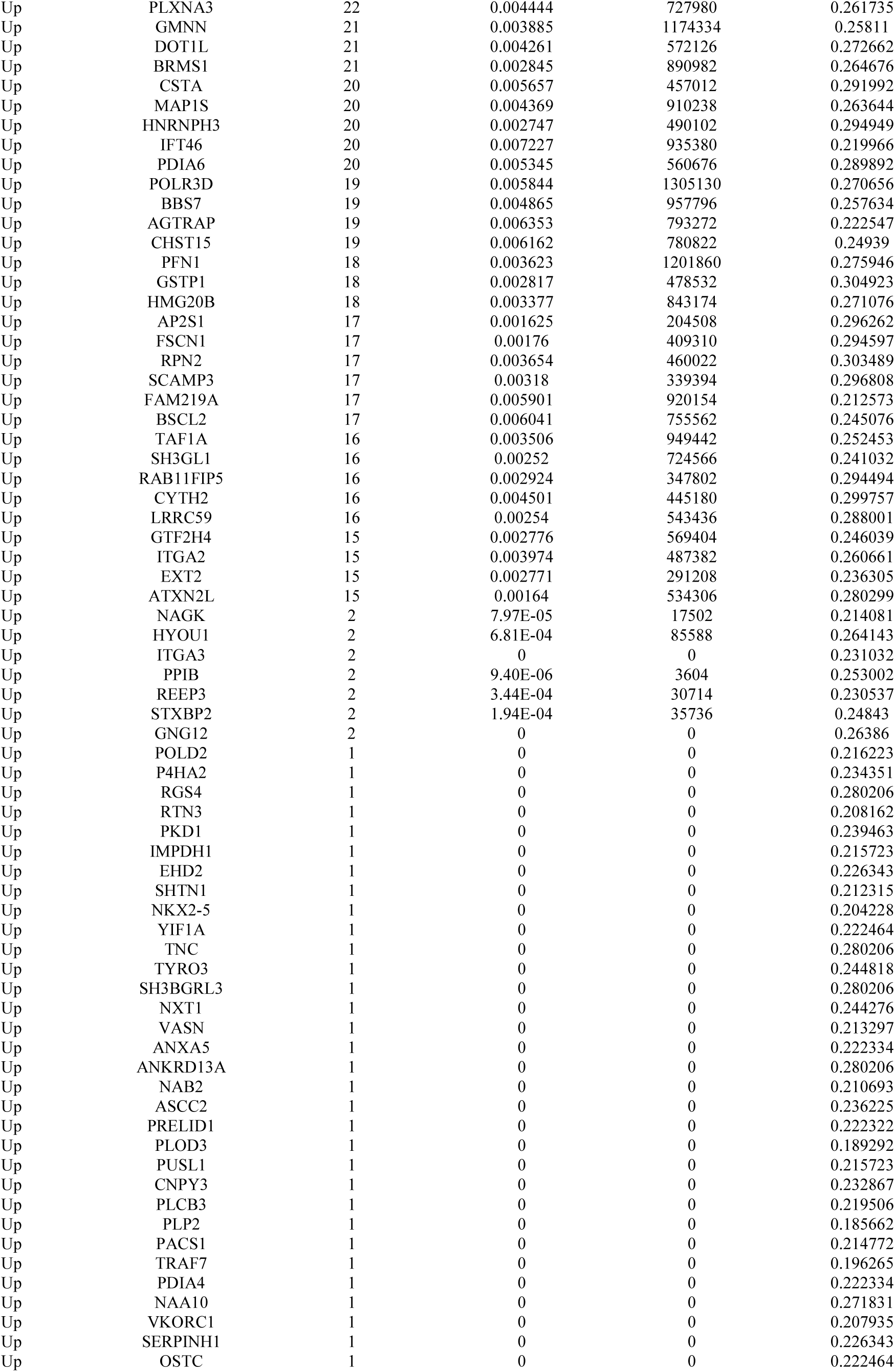
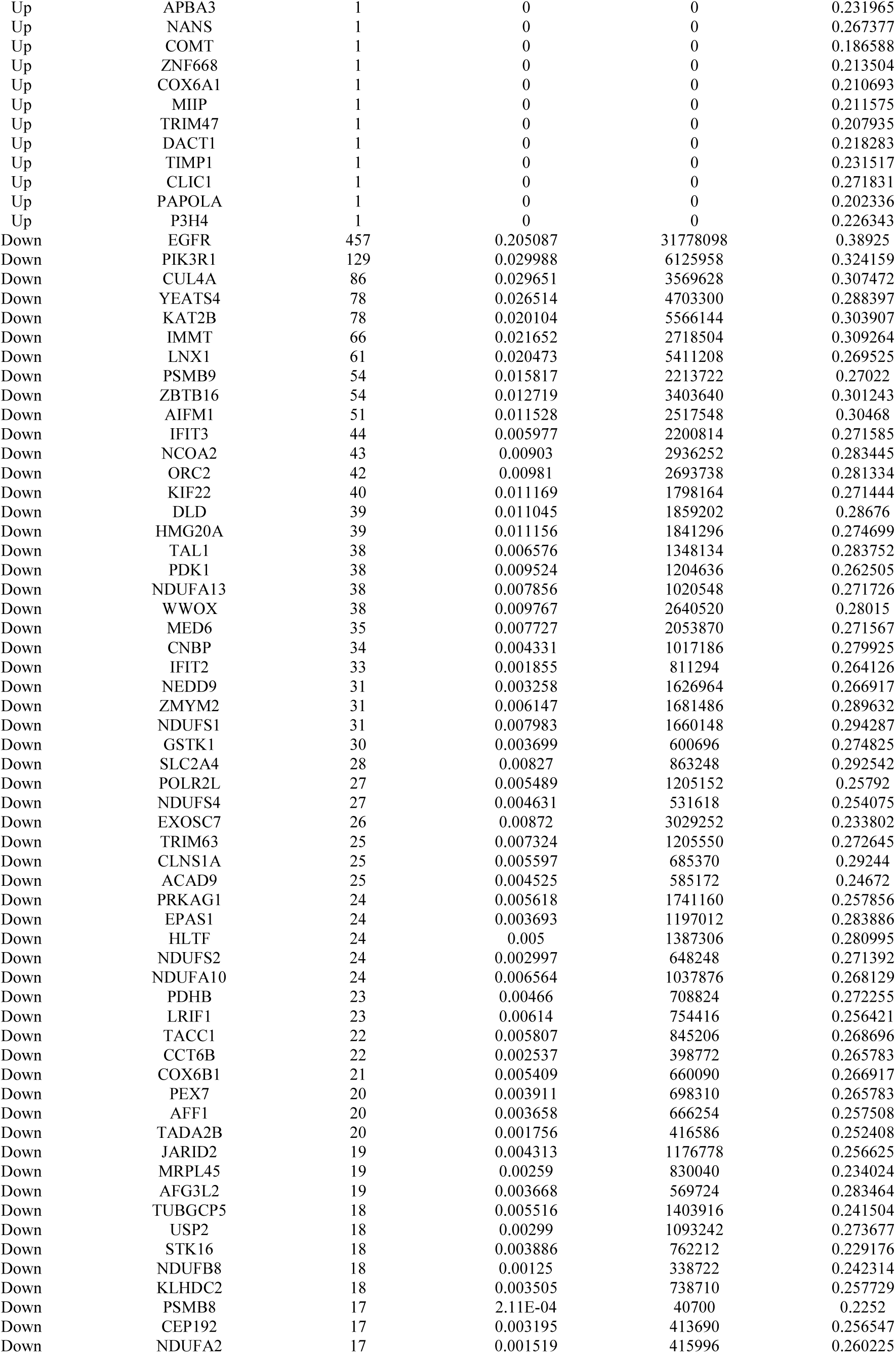
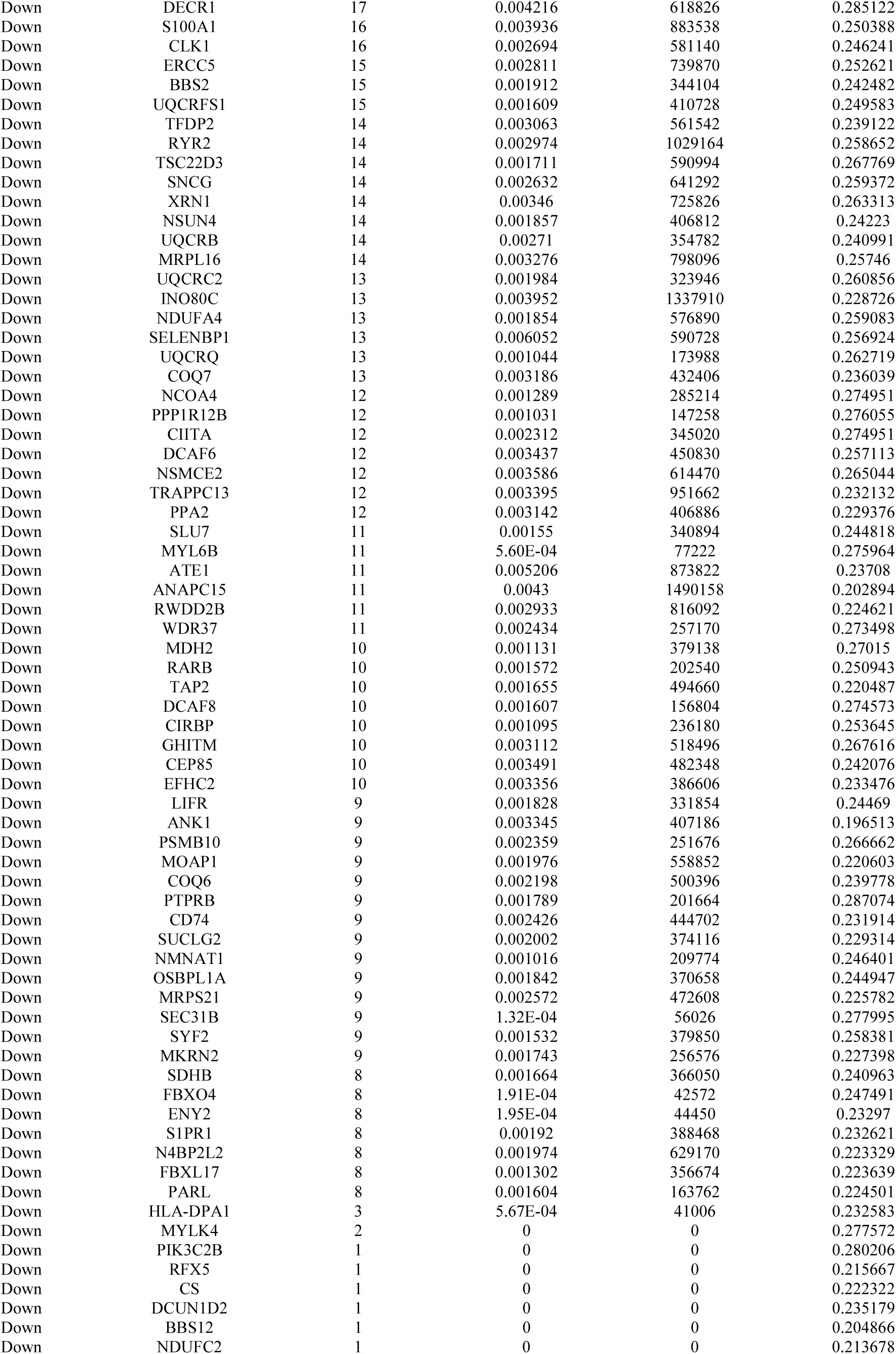
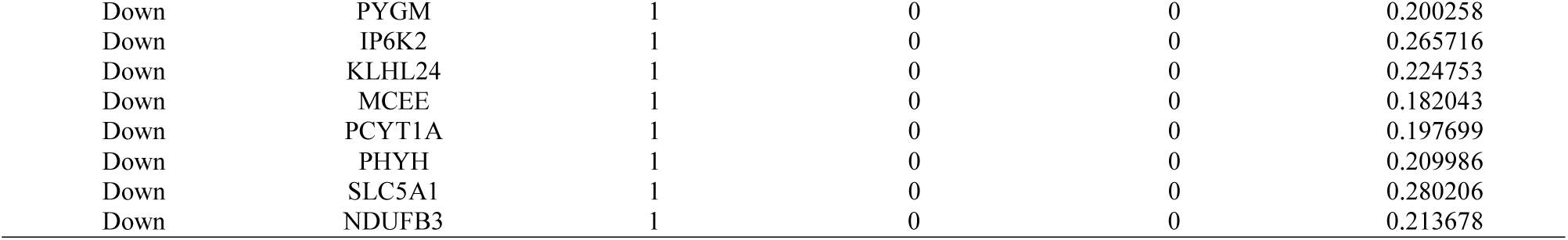
Topology table for up and down regulated genes.

**Table 6.** miRNA - target gene and TF - target gene interaction

| Regulation | Target Genes | Degree | MicroRNA | Regulation | Target Genes | Degree | TF |
| --- | --- | --- | --- | --- | --- | --- | --- |
| Up | MYH9 | 226 | hsa-mir-520e | UP | HSP90AA1 | 35 | RUNX1T1 |
| Up | TUBB | 202 | hsa-mir-8084 | UP | XPO1 | 33 | STAT1 |
| Up | XPO1 | 198 | hsa-mir-125a-5p | UP | SMARCA4 | 33 | EGR1 |
| Up | HSP90AA1 | 188 | hsa-mir-133a-3p | UP | HSPA5 | 22 | FOSB |
| Up | HSP90AB1 | 162 | hsa-mir-4801 | UP | ARRB2 | 20 | ARNT |
| Up | HSPA5 | 123 | hsa-mir-320d | UP | HSP90AB1 | 20 | JUNB |
| Up | FLNA | 106 | hsa-mir-212-3p | UP | HGS | 18 | TSG101 |
| Up | CCT3 | 54 | hsa-mir-342-3p | UP | TUBB | 16 | STAT3 |
| Up | ARRB2 | 54 | hsa-mir-3127-3p | UP | MYH9 | 16 | MAX |
| Up | CCT2 | 48 | hsa-mir-16-1-3p | UP | CCT3 | 15 | XRCC6 |
| Up | HGS | 47 | hsa-mir-4739 | UP | HSPB1 | 15 | FOS |
| Up | SMARCA4 | 44 | hsa-mir-1296-5p | UP | RPS2 | 13 | CREB1 |
| Up | RPS2 | 40 | hsa-mir-3943 | UP | FLNA | 9 | VHL |
| Up | TNFRSF1A | 22 | hsa-mir-29a-3p | UP | TNFRSF1A | 6 | DAXX |
| Up | HSPB1 | 22 | hsa-mir-148a-3p | UP | CCT2 | 3 | YY1 |
| Down | PIK3R1 | 131 | hsa-mir-138-5p | Down | KAT2B | 47 | TWIST1 |
| Down | NCOA2 | 114 | hsa-mir-539-5p | Down | ZBTB16 | 39 | GATA2 |
| Down | EGFR | 83 | hsa-mir-132-3p | Down | NCOA2 | 34 | AHR |
| Down | IFIT3 | 78 | hsa-mir-449a | Down | PIK3R1 | 34 | GTF2H1 |
| Down | PSMB9 | 68 | hsa-mir-200c-5p | Down | EGFR | 27 | STAT5B |
| Down | KAT2B | 62 | hsa-mir-320c | Down | ORC2 | 22 | CDC5L |
| Down | ZBTB16 | 53 | hsa-mir-3908 | Down | LNX1 | 21 | FOXO1 |
| Down | CUL4A | 53 | hsa-mir-16-5p | Down | IMMT | 13 | POU3F2 |
| Down | DLD | 41 | hsa-mir-30d-5p | Down | PSMB9 | 10 | TOPORS |
| Down | ORC2 | 40 | hsa-mir-7-5p | Down | CUL4A | 10 | USF1 |
| Down | YEATS4 | 35 | hsa-mir-376a-5p | Down | AIFM1 | 8 | MYC |
| Down | IMMT | 33 | hsa-mir-182-5p | Down | YEATS4 | 6 | MLLT10 |
| Down | KIF22 | 33 | hsa-mir-146a | Down | IFIT3 | 5 | STAT2 |
| Down | AIFM1 | 13 | hsa-mir-484 | Down | KIF22 | 4 | ELK1 |
| Down | LNX1 | 5 | hsa-let-7b-5p | Down | DLD | 3 | NFYA |

### Construction of the PPI network and module analysis

Using Cytoscape, a HIPPIE interactome database was used to establish a PPI network of these DEGs, with 4194 nodes and 8352 edges (Fig. 3). Based on the HIPPIE database, the DEGs with the highest PPI scores identified by the 4 centrality methods are shown in Table 4. The hub genes were obtained using the 4 centrality methods, including HSP90AA1, ARRB2, MYH9, HSP90AB1, FLNA, EGFR, PIK3R1, CUL4A, YEATS4 and KAT2B. A significant module was constructed from the PPI network of the DEGs using PEWCC1, including module 1 had 35 nodes and 124 edges (Fig.4A) and module 2 had 14 nodes and 34 edges (Fig.4B). GO and REACTOME pathway enrichment analysis showed that genes in these modules were markedly enriched in disease, immune system, cytoplasm, neutrophil degranulation, protein binding, infectious disease, SARS-CoV infections, localization, organic substance transport, the citric acid (TCA) cycle and respiratory electron transport, respiratory electron transport and metabolism.

**Fig. 3.**
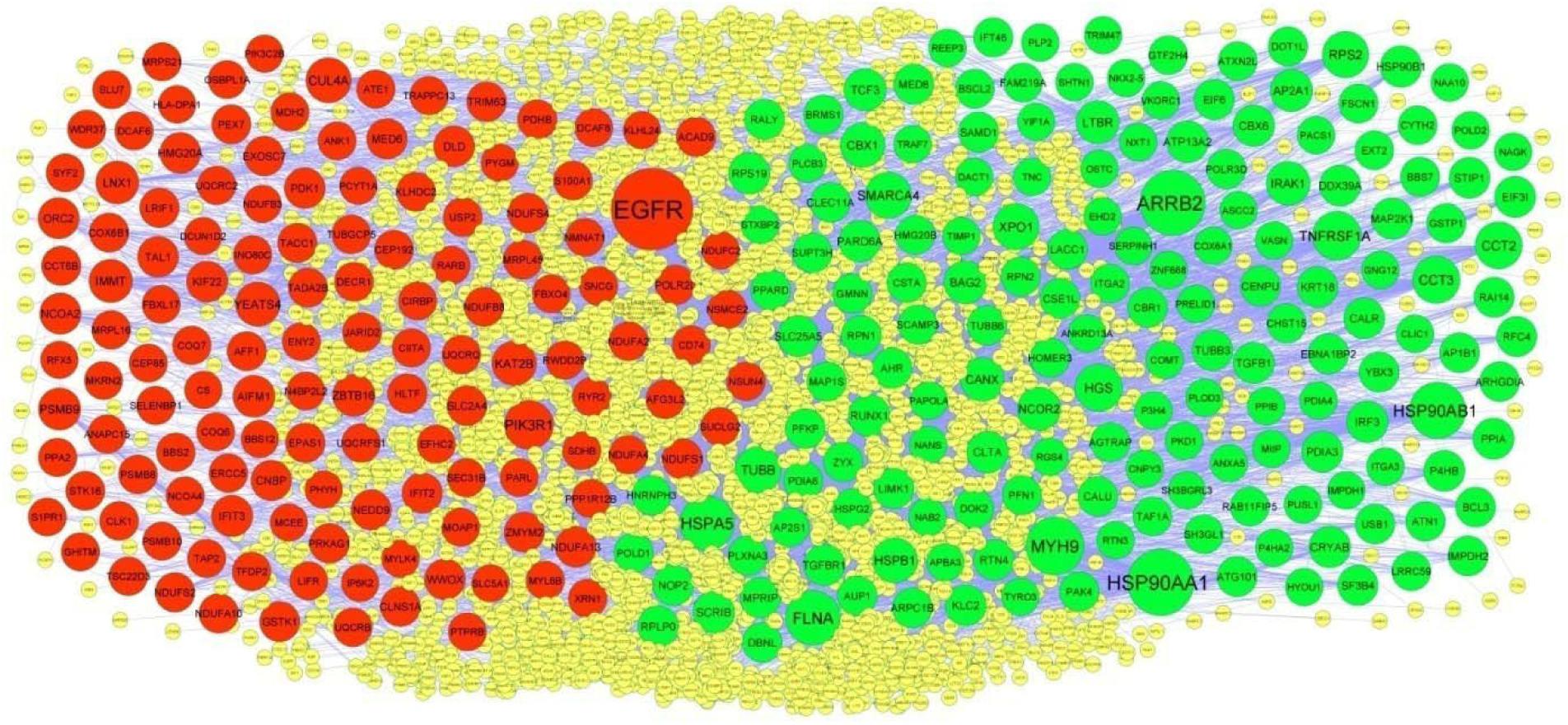
PPI network of DEGs. Up regulated genes are marked in green; down regulated genes are marked in red

**Fig. 4.**
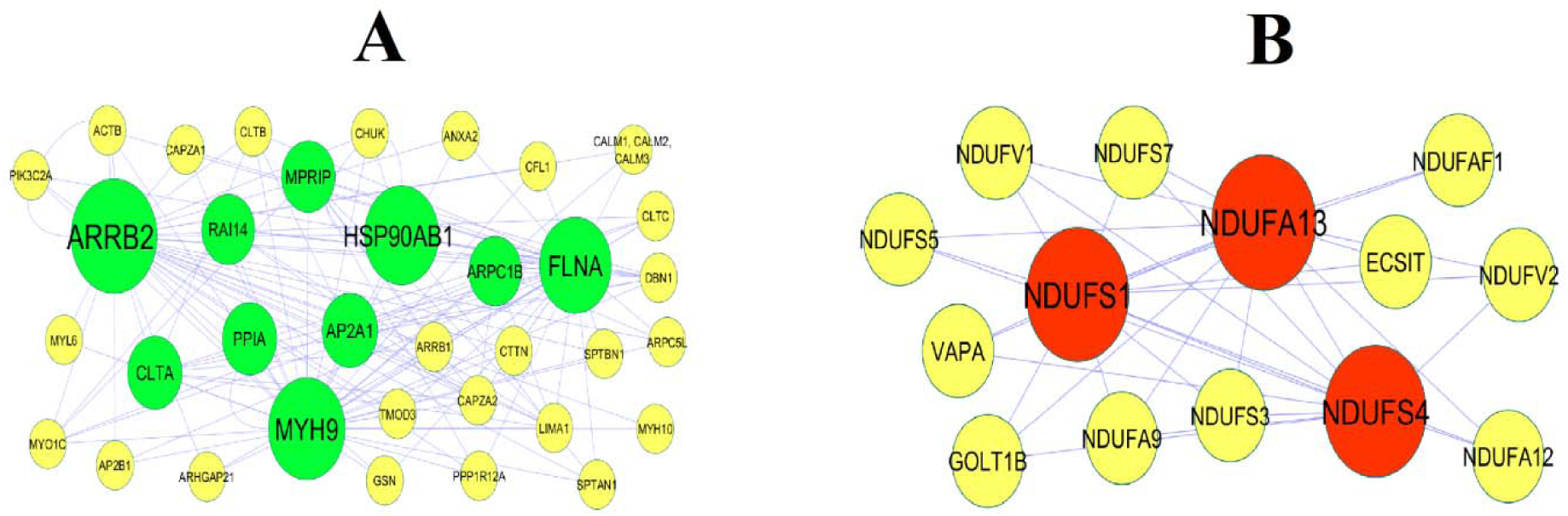
Modules of isolated form PPI of DEGs. (A) The most significant module was obtained from PPI network with 35 nodes and 124 edges for up regulated genes (B) The most significant module was obtained from PPI network with 14 nodes and 34 edges for down regulated genes. Up regulated genes are marked in green; down regulated genes are marked in red

### MiRNA-hub gene regulatory network construction

According to the information in miRNet database and Cytoscape databases, the miRNA-hub gene regulatory network relationships of miRNA and hub genes were obtained (Fig. 5). After comparing the targets with hub genes, we found that MYH9 was the potential target of 226 miRNAs (ex; hsa-mir-520e); TUBB was the potential target of 202 miRNAs (ex; hsa-mir-8084); XPO1 was the potential target of 198 miRNAs (ex; hsa-mir-125a-5p); HSP90AA1 was the potential target of 188 miRNAs (ex; hsa-mir-133a-3p); HSP90AB1 was the potential target of 162 miRNAs (ex; hsa-mir-4801); PIK3R1 was the potential target of 131 miRNAs (ex; hsa-mir-138-5p); NCOA2 was the potential target of 114 miRNAs (ex; hsa-mir-539-5p); EGFR was the potential target of 83 miRNAs (ex; hsa-mir-132-3p); IFIT3 was the potential target of 78 miRNAs (ex; hsa-mir-449a); PSMB9 was the potential target of 68 miRNAs (ex; hsa-mir-200c-5p).

**Fig. 5.**
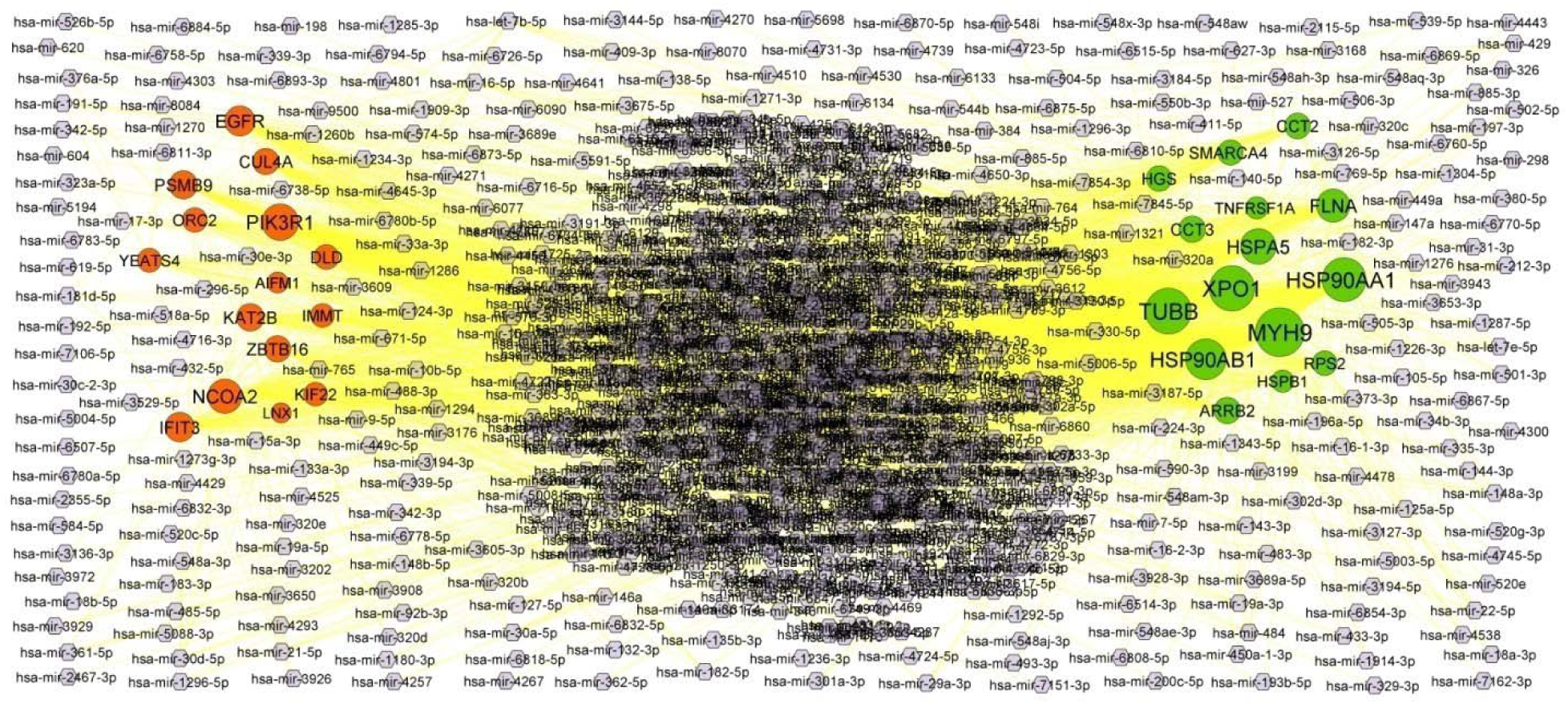
Target gene - miRNA regulatory network between target genes. The purple color diamond nodes represent the key miRNAs; up regulated genes are marked in green; down regulated genes are marked in red.

### TF-hub gene regulatory network construction

According to the information in NetworkAnalyst database and Cytoscape databases, the TF-hub gene regulatory network relationships of TF and hub genes were obtained (Fig. 6). After comparing the targets with hub genes, we found that HSP90AA1 was the potential target of 35 TFs (ex; RUNX1T1); XPO1 was the potential target of 33 TFs (ex; STAT1); SMARCA4 was the potential target of 33 TFs (ex; EGR1); HSPA5 was the potential target of 22 TFs (ex; FOSB); ARRB2 was the potential target of 20 TFs (ex; ARNT); KAT2B was the potential target of 47 TFs (ex; TWIST1); ZBTB16 was the potential target of 47 TFs (ex; GATA2); ZBTB16 was the potential target of 39 TFs (ex; GATA2); NCOA2 was the potential target of 34 TFs (ex; AHR); PIK3R1 was the potential target of 34 TFs (ex; GTF2H1); EGFR was the potential target of 27 TFs (ex; STAT5B).

**Fig. 6.**
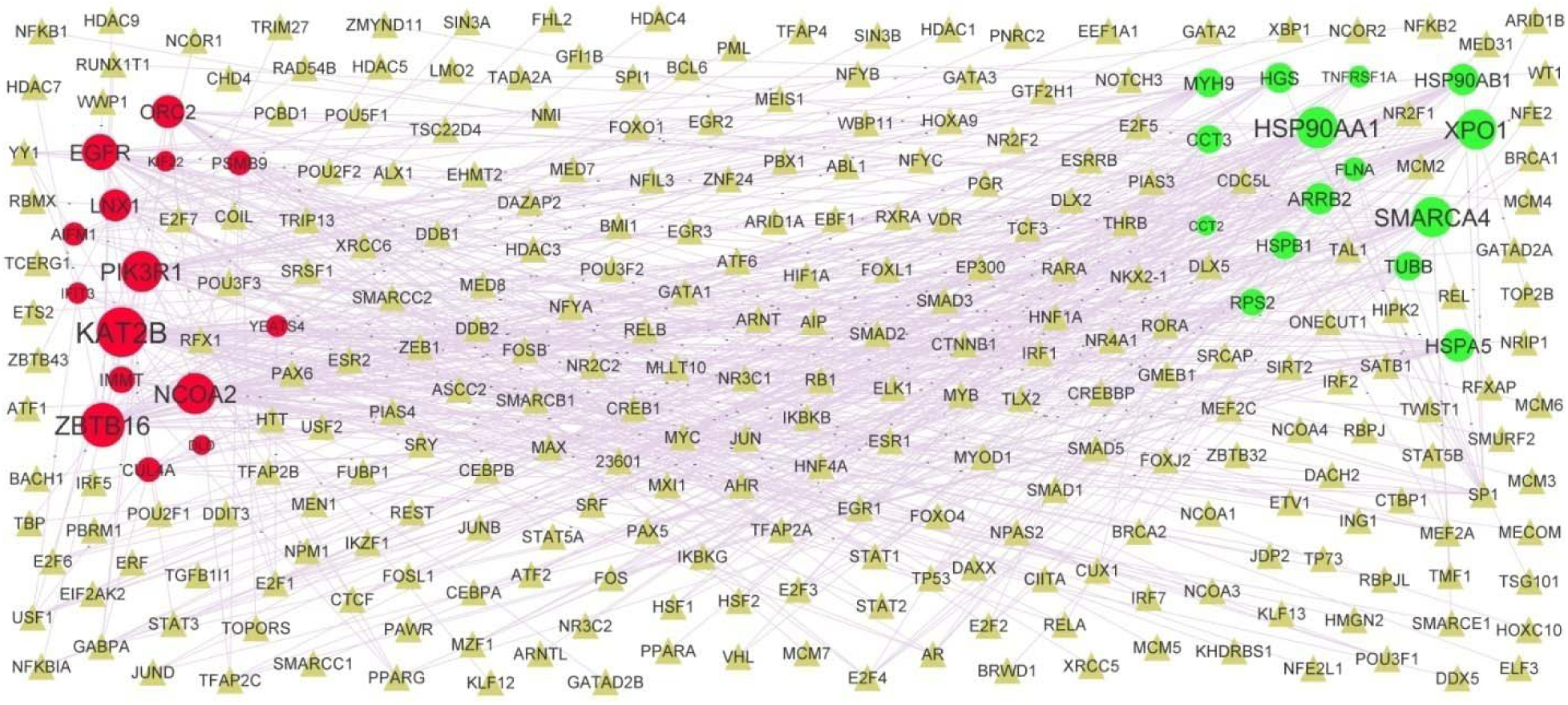
Target gene - TF regulatory network between target genes. The gray color triangle nodes represent the key TFs; up regulated genes are marked in green; down regulated genes are marked in red.

### Validation of hub genes by receiver operating characteristic curve (ROC) analysis

A ROC curve was plotted to evaluate the diagnostic value of HSP90AA1, ARRB2, MYH9, HSP90AB1, FLNA, EGFR, PIK3R1, CUL4A, YEATS4 and KAT2B (Fig. 7). The AUCs for the 10 hub genes were 0.953, 0.941, 0.976, 0.948, 0.931, 0.969, 0,958, 0.906, 0.912 and 0.950, respectively. These hub genes show good diagnostics values.

**Fig. 7.**
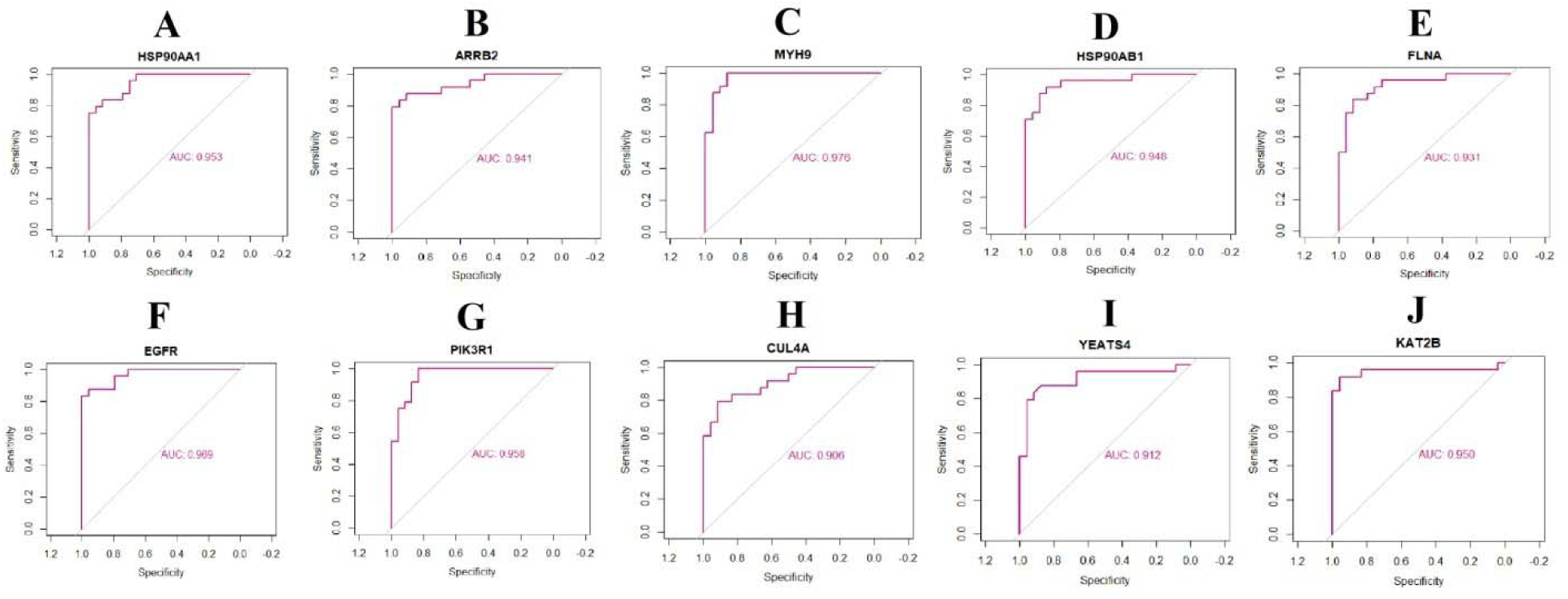
ROC curve analyses of hub genes. A) HSP90AA1 B) ARRB2 C) MYH9 D) HSP90AB1 E) FLNA F) EGFR G) PIK3R1 H) CUL4A I) YEATS4 J) KAT2B

## Discussion

Although many relevant investigation of HF have been operated, early diagnoses, adequacy of treatment and prognosis for HF remain poorly concluded. For diagnosis and treatment, it is vital to more interpret the molecular mechanisms resulting in occurrence and advancement. Bioinformatics analysis is progressively adopted to screen out biomarkers have a guiding role in the diagnosis and treatment of HF [37].

In this investigation, we performed a series of bioinformatics analysis to screen key genes and pathways. The expression profiling by high throughput sequencing data found that 464 up regulated genes and 466 down regulated genes were identified in HF samples compared to normal control samples. ZFP57 [38] and ANK1 [39] contributes to the progression of diabetics, but these genes might be novel target for HF. TNC (tenascin C) has been shown to be activated in cardiac hypertrophy [40]. CCL2 is mainly involved in the progression of myocardial infarction [41]. SPP1[42] and IGSF1 [43] plays an important role in obesity, but these genes might be novel target for HF. Kiczak et al [44] found that TIMP1 was highly expressed in the HF.

GO and REACTOME pathway enrichment analyses were used to investigate the interactions of these DEGs. SARS-CoV infections [45], asparagine N-linked glycosylation [46], neutrophil degranulation [47], immune system [48], respiratory electron transport [49], metabolism [50], complex I biogenesis [51], neddylation [52], localization [53], membrane [54], protein binding [55], small molecule metabolic process [56] and were responsible for progression of HF. Han et al [57], Yamada et al [58], Wang et al [59], García-Manzanares et al [60], Raitoharju et al [61], King et al [62], Hirokawa et al [63], Kahali et al [64], Sun et al [65] and Kuhn et al [66] found expression of DHCR24, STXBP2, CLEC5A, XPO1, ADAM8, IRF3, DOT1L, PPP1R3B, IFIT3 and PCSK6 in myocardial infarction and indicated it as a potential gene markers. Expression of CYP1B1 [67], SLC7A1 [68], MYH9 [69], LAT2 [70], FXYD5 [71], CAMK1 [72], TGFBR1 [73], HSP90AB1 [74], PLEKHA7 [75], AGTRAP (angiotensin II receptor associated protein) [76], SLC16A9 [77], ACADSB (acyl-CoA dehydrogenase short/branched chain) [78], IMPA1 [79], CD300LG [80], CIRBP (cold inducible RNA binding protein) [81], PIK3R1 [82], YEATS4 [83], USP2 [84], NEDD9 [85], CHCHD5 [86] and ERAP1 [87] promotes hypertension. HSPB1 [88], CRYAB (crystallin alpha B) [89], ANXA5 [90], CCR2 [91], RGS4 [92], TNFRSF1A [93], XBP1 [94], NKX2-5 [95], NEU1 [96], GSTP1 [97], COMT (catechol-O-methyltransferase) [98], LIMK1 [99], CAMKK1 [100], CD276 [101], SMARCA4 [102], ADORA2B [103], ACOT1 [104], RGN (regucalcin) [105], PPA2 [106], KAT2B [107], PDK1 [108], CS (citrate synthase) [109], FGF12 [110], AQP4 [111], LMOD2 [112], SELENBP1 [113], MB (myoglobin) [114], S100A1 [115], RYR2 [116], GPC5 [117], JARID2 [118], EGFR (epidermal growth factor receptor) [119], FUNDC1 [120], S1PR1 [121], EPAS1 [122] and OSBPL11 [123] genes are a potential biomarkers for the detection and prognosis of HF at an early age. A previous study reported that CALR (calreticulin) [124], BSCL2 [125], PKD1 [126], TMBIM1 [127], CHST15 [128], NAA10 [129], TCF3 [130], CNN1 [131], TAF1A [132], ACAD9 [133], KLHL24 [134], MYOM2 [135], TRIM63 [136], CTNNA3 [137], NLRC5 [138]. KLF9 [139], MYLK3 [140], RBM20 [141], GSTK1 [142], UQCRFS1 [143], NDUFS2 [144] and COX6B1 [145] are expressed in cardiomyopathy. Other research have revealed that RTN4 [146], NCF2 [147], ARHGAP9 [148], LIPG (lipase G, endothelial type) [149], BCL3 [150], HSPG2 [151], APOBR (apolipoprotein B receptor) [152], ITGA2 [153], PPIA (peptidylprolylisomerase A) [154], IRAK1 [155], VKORC1 [156], RNLS (renalase, FAD dependent amine oxidase) [157], HLA-F [158], FBXL17 [159], COL11A2 [160] and NDUFC2 [161] are expressed in coronary heart disease, suggesting that it might also function in coronary heart disease transformation and development. CCR1 [162], MANF (mesencephalic astrocyte derived neurotrophic factor) [163], HSP90AA1 [164], ARRB2 [165], SLC39A13 [166], P4HA2 [167], HECTD3 [168], CNPY2 [169], ECHDC2 [170], NDUFS4 [171] and IMMT (inner membrane mitochondrial protein) [172] are a key initiators of ischemic cardiac diseases. Mounting evidence indicates that expression of PEA15 [173], CD48 [174], EDEM2 [175], CD55 [176], NCOR2 [177], EXT2 [178], SPRED1 [179], PDIA6 [180], CD300E [181], TCF19 [182], ABHD15 [183], OXSM (3-oxoacyl-ACP synthase, mitochondrial) [184], MGST3 [185], COQ7 [186], ACSL5 [187], ANK1 [188], PYGM (glycogen phosphorylase, muscle associated) [189], FBXO40 [190], SLC2A4 [191], HLA-DOA [192], TAP2 [193], HLA-DPA1 [194], NSMCE2 [195], NDUFA4 [196], HMG20A [197], AMY2B [198] and ACYP2 [199] might be involved in the pathogenesis of diabetics, but these genes might be novel target for HF. Recent evidence indicates that the SIRPA (signal regulatory protein alpha) [200], PFN1 [201], EIF6 [202], AHR (aryl hydrocarbon receptor) [203], RUNX1 [204], IDO1 [205], PDHB (pyruvate dehydrogenase E1 subunit beta) [206], NDUFS1 [207], MTUS1 [208], ZNF418 [209] and MTMR14 [210] are the key biomarkers in cardiac hypertrophy. Previous studies had shown that the expression of CLIC1 [211], ARPC1B [212], FLNA (filamin A) [213], HILPDA (hypoxia inducible lipid droplet associated) [214], RTN3 [215], G0S2 [216], CALU (calumenin) [217], MYDGF (myeloid derived growth factor) [218], LTBR (lymphotoxin beta receptor) [219], GGCX (gamma-glutamyl carboxylase) [220], SAMD1 [221], ACAT1 [222], NNT (nicotinamide nucleotide transhydrogenase) [223] and ATG14 [224] were closely related to the occurrence of atherosclerosis. HSPA5 [225], NMB (neuromedin B) [226], ELP5 [227], NLGN2 [228], RAB23 [229], MBOAT7 [230], CEP19 [231], PFKP (phosphofructokinase, platelet) [232], TKT (transketolase) [233], P4HB [234], CRYM (crystallin mu) [235], AMT (aminomethyltransferase) [236], MACROD2 [237], NUDT3 [238], DLD (dihydrolipoamide dehydrogenase) [239], NCOA2 [240], ZBTB16 [241], TFAM (transcription factor A, mitochondrial) [242], RASAL2 [243], NDUFB6 [244], HELZ2 [245] and CIITA (class II major histocompatibility complex transactivator) [246] were reported to be expressed in obesity, but these genes might be novel target for HF. TGFB1 [247], CD63 [248], DACT1 [249] and PSMB10 [250] have been reported to be crucial for the progression of atrial fibrillation. FAM20C [251] and CD74 [252] are important in the development of cardiac arrhythmia.

To explore the pathogenesis of HF, we constructed PPI network and isolated modules from PPI network for systematic analysis. The genes in the PPI network and modules with higher score were the hub genes that affected the progression of disease. CLTA (clathrin light chain A), RAI14, MPRIP (myosin phosphatase Rho interacting protein), AP2A1, CUL4A and NDUFA13 were the novel biomarkers for the progression of HF.

We further constructed a miRNA-hub gene regulatory network and TF-hub gene regulatory network for better understanding of the interaction between miRNA and hub genes, and TF and hub genes. Hsa-mir-125a-5p [253], hsa-mir-539-5p [254] and RUNX1T1 [255] were linked with progression of obesity, but these genes might be novel target for HF. Hsa-mir-138-5p [256] and STAT1 [257] were involved in the progression of HF. Hsa-mir-200c-5p [258] and STAT5B [259] were associated with development of diabetics, but these genes might be novel target for HF. EGR1 [260] and GATA2 [261] were liable for advancement of coronary heart disease. FOSB [262] and AHR [263] were associated with development of ischemic cardiac diseases. TWIST1 was responsible for progression of atherosclerosis [264]. TUBB (tubulin beta class I), hsa-mir-520e, hsa-mir-8084, hsa-mir-133a-3p, hsa-mir-4801, hsa-mir-132-3p, hsa-mir-449a, ARNT, and GTF2H1 were the novel biomarkers for the progression of HF.

In summary, the present data provide a comprehensive bioinformatics analysis of DEGs that might be related to the progression of HF. We have identified 930 candidate DEGs with NGS data and integrated bioinformatics analyses. A variety of novel genes and signaling pathways might be associated in the pathogenesis of HF. We also conclude that HSP90AA1, ARRB2, MYH9, HSP90AB1, FLNA, EGFR, PIK3R1, CUL4A, YEATS4 and KAT2B might be associated with progression of HF. These findings could lead to an increase in our understanding of the etiology and underlying molecular events of HF.

## Acknowledgement

I thank Marina Stolina, Amgen Inc, Cardiometabolic Disorders, One Amgen Center Drive, Thousand Oaks, California, USA, very much, the author who deposited their profiling by high throughput sequencing dataset GSE161472, into the public GEO database.

## Ethical approval

This article does not contain any studies with human participants or animals performed by any of the authors.

## Informed consent

No informed consent because this study does not contain human or animals participants.

## Availability of data and materials

The datasets supporting the conclusions of this article are available in the GEO (Gene Expression Omnibus) (https://www.ncbi.nlm.nih.gov/geo/) repository. [(GSE161472) https://www.ncbi.nlm.nih.gov/geo/query/acc.cgi?acc=GSE161472)]

## Consent for publication

Not applicable.

## Author Contributions

B. V. - Writing original draft, and review and editing

C. V. - Software and investigation

## References

1. Tanai E, Frantz S. Pathophysiology of Heart Failure. Compr Physiol. 2015;6(1):187–214. doi:10.1002/cphy.c140055

2. Yap J. Risk stratification in heart failure: Existing challenges and potential promise. Int J Cardiol. 2020;313:97–98. doi:10.1016/j.ijcard.2020.04.042

3. Ohkuma T, Komorita Y, Peters SAE, Woodward M. Diabetes as a risk factor for heart failure in women and men: a systematic review and meta-analysis of 47 cohorts including 12 million individuals. Diabetologia. 2019;62(9):1550–1560. doi:10.1007/s00125-019-4926-x

4. Messerli FH, Rimoldi SF, Bangalore S. The Transition From Hypertension to Heart Failure: Contemporary Update. JACC Heart Fail. 2017;5(8):543–551. doi:10.1016/j.jchf.2017.04.012

5. Ayinapudi K, Samson R, Le Jemtel TH, Marrouche NF, Oparil S. Weight Reduction for Obesity-Induced Heart Failure with Preserved Ejection Fraction. Curr Hypertens Rep. 2020;22(8):47. doi:10.1007/s11906-020-01074-w

6. Skrzynia C, Berg JS, Willis MS, Jensen BC. Genetics and heart failure: a concise guide for the clinician. Curr Cardiol Rev. 2015;11(1):10–17. doi:10.2174/1573403x09666131117170446

7. Escolar V, Lozano A, Larburu N, Kerexeta J, Álvarez R, Juez B, Echebarria A, Azcona A, Artola G. Impact of environmental factors on heart failure decompensations. ESC Heart Fail. 2019;6(6):1226–1232. doi:10.1002/ehf2.12506

8. Ayoub KF, Pothineni NVK, Rutland J, Ding Z, Mehta JL. Immunity, Inflammation, and Oxidative Stress in Heart Failure: Emerging Molecular Targets. Cardiovasc Drugs Ther. 2017;31(5-6):593–608. doi:10.1007/s10557-017-6752-z

9. Ziaeian B, Fonarow GC. Epidemiology and aetiology of heart failure. Nat Rev Cardiol. 2016;13(6):368–378. doi:10.1038/nrcardio.2016.25

10. Latronico MV, Elia L, Condorelli G, Catalucci D. Heart failure: targeting transcriptional and post-transcriptional control mechanisms of hypertrophy for treatment. Int J Biochem Cell Biol. 2008;40(9):1643–1648. doi:10.1016/j.biocel.2008.03.002

11. Hua X, Wang YY, Jia P, Xiong Q, Hu Y, Chang Y, Lai S, Xu Y, Zhao Z, Song J. Multi-level transcriptome sequencing identifies COL1A1 as a candidate marker in human heart failure progression. BMC Med. 2020;18(1):2. doi:10.1186/s12916-019-1469-4

12. Zeng L, Gu N, Chen J, Jin G, Zheng Y. IRX1 hypermethylation promotes heart failure by inhibiting CXCL14 expression. Cell Cycle. 2019;18(23):3251–3262. doi:10.1080/15384101.2019.1673099

13. Wang C, Wang F, Cao Q, Li Z, Huang L, Chen S. The Effect of Mecp2 on Heart Failure. Cell Physiol Biochem. 2018;47(6):2380–2387. doi:10.1159/000491610

14. Ma J, Lu L, Guo W, Ren J, Yang J. Emerging Role for RBM20 and its Splicing Substrates in Cardiac Function and Heart Failure. Curr Pharm Des. 2016;22(31):4744–4751. doi:10.2174/1381612822666160701145322

15. Schilling J, Kelly DP. The PGC-1 cascade as a therapeutic target for heart failure. J Mol Cell Cardiol. 2011;51(4):578–583. doi:10.1016/j.yjmcc.2010.09.021

16. Stylianidis V, Hermans KCM, Blankesteijn WM. Wnt Signaling in Cardiac Remodeling and Heart Failure. Handb Exp Pharmacol. 2017;243:371–393. doi:10.1007/164_2016_56

17. Chen K, Chen W, Liu SL, Wu TS, Yu KF, Qi J, Wang Y, Yao H, Huang XY, Han Y, et al. Epigallocatechingallate attenuates myocardial injury in a mouse model of heart failure through TGFβ1/Smad3 signaling pathway. Mol Med Rep. 2018;17(6):7652–7660. doi:10.3892/mmr.2018.8825

18. Wang X, Meng H, Wang Q, Shao M, Lu W, Chen X, Jiang Y, Li C, Wang Y, Tu P. et al. Baoyuan decoction ameliorates apoptosis via AT1-CARP signaling pathway in H9C2 cells and heart failure post-acute myocardial infarction rats. J Ethnopharmacol. 2020;252:112536. doi:10.1016/j.jep.2019.112536

19. Ananthakrishnan R, Moe GW, Goldenthal MJ, Marín-García J. Akt signaling pathway in pacing-induced heart failure. Mol Cell Biochem. 2005;268(1-2):103–110. doi:10.1007/s11010-005-3699-3

20. Xu Y, Li X, Liu X, Zhou M. Neuregulin-1/ErbB signaling and chronic heart failure. Adv Pharmacol. 2010;59:31–51. doi:10.1016/S1054-3589(10)59002-1

21. Abplanalp WT, John D, Cremer S, Assmus B, Dorsheimer L, Hoffmann J, Becker-Pergola G, Rieger MA, Zeiher AM, Vasa-Nicotera M, et al. Single-cell RNA-sequencing reveals profound changes in circulating immune cells in patients with heart failure. Cardiovasc Res. 2021;117(2):484–494. doi:10.1093/cvr/cvaa101

22. Clough E, Barrett T. The Gene Expression Omnibus Database. Methods Mol Biol. 2016;1418:93–110. doi:10.1007/978-1-4939-3578-9_5

23. Ritchie ME, Phipson B, Wu D, Hu Y, Law CW, Shi W, Smyth GK. limma powers differential expression analyses for RNA-sequencing and microarray studies. Nucleic Acids Res. 2015;43(7):e47. doi:10.1093/nar/gkv007

24. Thomas PD. The Gene Ontology and the Meaning of Biological Function. Methods Mol Biol. 2017;1446:15 doi:10.1007/978-1-4939-3743-1_2

25. Fabregat A, Jupe S, Matthews L, Sidiropoulos K, Gillespie M, Garapati P, Haw R, Jassal B, Korninger F, May B et al The Reactome Pathway Knowledgebase. Nucleic Acids Res. 2018;46(D1):D649–D655. doi:10.1093/nar/gkx1132

26. Reimand J, Kull M, Peterson H, Hansen J, Vilo J. g:Profiler--a web-based toolset for functional profiling of gene lists from large-scale experiments. Nucleic Acids Res. 2007;35(Web Server issue):W193–W200. doi:10.1093/nar/gkm226

27. Alanis-Lobato G, Andrade-Navarro MA, Schaefer MH. HIPPIE v2.0: enhancing meaningfulness and reliability of protein-protein interaction networks. Nucleic Acids Res. 2017;45(D1):D408–D414. doi:10.1093/nar/gkw985

28. Shannon P, Markiel A, Ozier O, Baliga NS, Wang JT, Ramage D, Amin N, Schwikowski B, Ideker T Cytoscape: a software environment for integrated models of biomolecular interaction networks. Genome Res 2003;13(11):2498–2504. doi:10.1101/gr.1239303

29. Przulj N, Wigle DA, Jurisica I. Functional topology in a network of protein interactions. Bioinformatics. 2004;20(3):340–348. doi:10.1093/bioinformatics/btg415

30. Nguyen TP, Liu WC, Jordán F. Inferring pleiotropy by network analysis: linked diseases in the human PPI network. BMC Syst Biol. 2011;5:179. doi:10.1186/1752-0509-5-179

31. Shi Z, Zhang B. Fast network centrality analysis using GPUs. BMC Bioinformatics. 2011;12:149. doi:10.1186/1471-2105-12-149

32. Fadhal E, Gamieldien J, Mwambene EC. Protein interaction networks as metric spaces: a novel perspective on distribution of hubs. BMC Syst Biol. 2014;8:6. doi:10.1186/1752-0509-8-6

33. Zaki N, Efimov D, Berengueres J. Protein complex detection using interaction reliability assessment and weighted clustering coefficient. BMC Bioinformatics. 2013;14:163. doi:10.1186/1471-2105-14

34. Fan Y, Xia J (2018) miRNet-Functional Analysis and Visual Exploration of miRNA-Target Interactions in a Network Context. Methods Mol Biol 1819:215–233. doi:10.1007/978-1-4939-8618-7_10

35. Zhou G, Soufan O, Ewald J, Hancock REW, Basu N, Xia J (2019) NetworkAnalyst 3.0: a visual analytics platform for comprehensive gene expression profiling and meta-analysis. Nucleic Acids Res 47:W234–W241. doi:10.1093/nar/gkz240

36. Robin X, Turck N, Hainard A, Tiberti N, Lisacek F, Sanchez JC, Müller M. pROC: an open-source package for R and S+ to analyze and compare ROC curves. BMC Bioinformatics 2011;12:77. doi:10.1186/1471-2105-12-77

37. Li X, Li B, Jiang H. Identification of time□series differentially expressed genes and pathways associated with heart failure post□myocardial infarction using integrated bioinformatics analysis. Mol Med Rep. 2019;19(6):5281–5290. doi:10.3892/mmr.2019.10190

38. Touati A, Errea-Dorronsoro J, Nouri S, Halleb Y, Pereda A, Mahdhaoui N, Ghith A, Saad A, Perez de Nanclares G, H’mida Ben Brahim D. Transient neonatal diabetes mellitus and hypomethylation at additional imprinted loci: novel ZFP57 mutation and review on the literature. Acta Diabetol. 2019;56(3):301–307. doi:10.1007/s00592-018-1239-3

39. Sun L, Zhang X, Wang T, Chen M, Qiao H. Association of ANK1 variants with new-onset type 2 diabetes in a Han Chinese population from northeast China. Exp Ther Med. 2017;14(4):3184–3190. doi:10.3892/etm.2017.4866

40. Song L, Wang L, Li F, Yukht A, Qin M, Ruther H, Yang M, Chaux A, Shah PK, Sharifi BG. Bone Marrow-Derived Tenascin-C Attenuates Cardiac Hypertrophy by Controlling Inflammation. J Am Coll Cardiol. 2017;70(13):1601–1615. doi:10.1016/j.jacc.2017.07.789

41. Dewald O, Zymek P, Winkelmann K, Koerting A, Ren G, Abou-Khamis T, Michael LH, Rollins BJ, Entman ML, Frangogiannis NG. CCL2/Monocyte Chemoattractant Protein-1 regulates inflammatory responses critical to healing myocardial infarcts. Circ Res. 2005;96(8):881–889. doi:10.1161/01.RES.0000163017.13772.3a

42. Yang H, Graham LC, Reagan AM, Grabowska WA, Schott WH, Howell GR. Transcriptome profiling of brain myeloid cells revealed activation of Itgal, Trem1, and Spp1 in western diet-induced obesity. J Neuroinflammation. 2019;16(1):169. doi:10.1186/s12974-019-1527-z

43. Ghanny S, Zidell A, Pedro H, Joustra SD, Losekoot M, Wit JM, Aisenberg J. The IGSF1 Deficiency Syndrome May Present with Normal Free T4 Levels, Severe Obesity, or Premature Testicular Growth. J Clin Res Pediatr Endocrinol. 2020;10.4274/jcrpe.galenos.2020.2020.0125. doi:10.4274/jcrpe.galenos.2020.2020.0125

44. Kiczak L, Tomaszek A, Bania J, Paslawska U, Zacharski M, Noszczyk-Nowak A, Janiszewski A, Skrzypczak P, Ardehali H, Jankowska EA, et al. Expression and complex formation of MMP9, MMP2, NGAL, and TIMP1 in porcine myocardium but not in skeletal muscles in male pigs with tachycardia-induced systolic heart failure. Biomed Res Int. 2013;2013:283856. doi:10.1155/2013/283856

45. Belhadjer Z, Méot M, Bajolle F, Khraiche D, Legendre A, Abakka S, Auriau J, Grimaud M, Oualha M, Beghetti M, et al. Acute Heart Failure in Multisystem Inflammatory Syndrome in Children in the Context of Global SARS-CoV-2 Pandemic. Circulation. 2020;142(5):429–436. doi:10.1161/CIRCULATIONAHA.120.048360

46. Yang S, Chatterjee S, Cipollo J. The Glycoproteomics-MS for Studying Glycosylation in Cardiac Hypertrophy and Heart Failure. Proteomics Clin Appl. 2018;12(5):e1700075. doi:10.1002/prca.201700075

47. Bai B, Cheng M, Jiang L, Xu J, Chen H, Xu Y. High Neutrophil to Lymphocyte Ratio and Its Gene Signatures Correlate With Diastolic Dysfunction in Heart Failure With Preserved Ejection Fraction. Front Cardiovasc Med. 2021;8:614757. doi:10.3389/fcvm.2021.614757

48. De Angelis E, Pecoraro M, Rusciano MR, Ciccarelli M, Popolo A. Cross-Talk between Neurohormonal Pathways and the Immune System in Heart Failure: A Review of the Literature. Int J Mol Sci. 2019;20(7):1698. doi:10.3390/ijms20071698

49. Casademont J, Miró O. Electron transport chain defects in heart failure. Heart Fail Rev. 2002;7(2):131–139. doi:10.1023/a:1015372407647

50. Fukushima A, Milner K, Gupta A, Lopaschuk GD. Myocardial Energy Substrate Metabolism in Heart Failure : from Pathways to Therapeutic Targets. Curr Pharm Des. 2015;21(25):3654–3664. doi:10.2174/1381612821666150710150445

51. Pisano A, Cerbelli B, Perli E, Pelullo M, Bargelli V, Preziuso C, Mancini M, He L, Bates MG, Lucena JR, et al. Impaired mitochondrial biogenesis is a common feature to myocardial hypertrophy and end-stage ischemic heart failure. Cardiovasc Pathol. 2016;25(2):103–112. doi:10.1016/j.carpath.2015.09.009

52. Zou J, Ma W, Littlejohn R, Li J, Stansfield BK, Kim IM, Liu J, Zhou J, Weintraub NL, Su H. Transient inhibition of neddylation at neonatal stage evokes reversible cardiomyopathy and predisposes the heart to isoproterenol-induced heart failure. Am J Physiol Heart Circ Physiol. 2019;316(6):H1406–H1416. doi:10.1152/ajpheart.00806.2018

53. Treskatsch S, Feldheiser A, Shaqura M, Dehe L, Habazettl H, Röpke TK, Shakibaei M, Schäfer M, Spies CD, Mousa SA. Cellular localization and adaptive changes of the cardiac delta opioid receptor system in an experimental model of heart failure in rats. Heart Vessels. 2016;31(2):241–250. doi:10.1007/s00380-014-0620-6

54. Dangers L, Bréchot N, Schmidt M, Lebreton G, Hékimian G, Nieszkowska A, Besset S, Trouillet JL, Chastre J, Leprince P, et al. Extracorporeal Membrane Oxygenation for Acute Decompensated Heart Failure. Crit Care Med. 2017;45(8):1359–1366. doi:10.1097/CCM.0000000000002485

55. Rodríguez-Calvo R, Girona J, Alegret JM, Bosquet A, Ibarretxe D, Masana L. Role of the fatty acid-binding protein 4 in heart failure and cardiovascular disease. J Endocrinol. 2017;233(3):R173–R184. doi:10.1530/JOE-17-0031

56. Chen L, Song J, Hu S. Metabolic remodeling of substrate utilization during heart failure progression. Heart Fail Rev. 2019;24(1):143–154. doi:10.1007/s10741-018-9713-0

57. Han F, Chen Q, Su J, Zheng A, Chen K, Sun S, Wu H, Jiang L, Xu X, Yang M, et al. MicroRNA-124 regulates cardiomyocyte apoptosis and myocardial infarction through targeting Dhcr24. J Mol Cell Cardiol. 2019;132:178–188. doi:10.1016/j.yjmcc.2019.05.007

58. Yamada Y, Sakuma J, Takeuchi I, Yasukochi Y, Kato K, Oguri M, Fujimaki T, Horibe H, Muramatsu M, Sawabe M, et al. Identification of STXBP2 as a novel susceptibility locus for myocardial infarction in Japanese individuals by an exome-wide association study. Oncotarget. 2017;8(20):33527–33535. doi:10.18632/oncotarget.16536

59. Wang X, Hu Y, Wang Y, Shen D, Tao G. CLEC5A knockdown protects against the cardiac dysfunction after Myocardial infarction by suppressing macrophage polarization, NLRP3 inflammasome activation and pyroptosis. Biochem Cell Biol. 2021;10.1139/bcb-2020-0672. doi:10.1139/bcb-2020-0672

60. García-Manzanares M, Tarazón E, Ortega A, Gil-Cayuela C, Martínez-Dolz L, González-Juanatey JR, Lago F, Portolés M, Roselló-Lletí E, Rivera M. XPO1 Gene Therapy Attenuates Cardiac Dysfunction in Rats with Chronic Induced Myocardial Infarction. J Cardiovasc Transl Res. 2020;13(4):593–600. doi:10.1007/s12265-019-09932-y

61. Raitoharju E, Seppälä I, Levula M, Kuukasjärvi P, Laurikka J, Nikus K, Huovila AP, Oksala N, Klopp N, Illig T, et al. Common variation in the ADAM8 gene affects serum sADAM8 concentrations and the risk of myocardial infarction in two independent cohorts. Atherosclerosis. 2011;218(1):127–133. doi:10.1016/j.atherosclerosis.2011.05.005

62. King KR, Aguirre AD, Ye YX, Sun Y, Roh JD, Ng RP Jr, Kohler RH, Arlauckas SP, Iwamoto Y, Savol A, et al. IRF3 and type I interferons fuel a fatal response to myocardial infarction. Nat Med. 2017;23(12):1481–1487. doi:10.1038/nm.4428

63. Hirokawa M, Morita H, Tajima T, Takahashi A, Ashikawa K, Miya F, Shigemizu D, Ozaki K, Sakata Y, Nakatani D, et al. A genome-wide association study identifies PLCL2 and AP3D1-DOT1L-SF3A2 as new susceptibility loci for myocardial infarction in Japanese. Eur J Hum Genet. 2015;23(3):374–380. doi:10.1038/ejhg.2014.110

64. Kahali B, Chen Y, Feitosa MF, Bielak LF, O’Connell JR, Musani SK, Hegde Y, Chen Y, Stetson LC, Guo X, et al. A Noncoding Variant Near PPP1R3B Promotes Liver Glycogen Storage and MetS, but Protects Against Myocardial Infarction. J Clin Endocrinol Metab. 2021;106(2):372–387. doi:10.1210/clinem/dgaa855

65. Kuhn TC, Knobel J, Burkert-Rettenmaier S, Li X, Meyer IS, Jungmann A, Sicklinger F, Backs J, Lasitschka F, Müller OJ, et al. Secretome Analysis of Cardiomyocytes Identifies PCSK6 (Proprotein Convertase Subtilisin/Kexin Type 6) as a Novel Player in Cardiac Remodeling After Myocardial Infarction. Circulation. 2020;141(20):1628–1644. doi:10.1161/CIRCULATIONAHA.119.044914

66. Sun J, Zhang Q, Liu X, Shang X. Downregulation of IFIT3 relieves the inflammatory response and myocardial fibrosis of mice with myocardial infarction and improves their cardiac function. Exp Anim. 2021;10.1538/expanim.21-0060. doi:10.1538/expanim.21-0060

67. Pingili AK, Jennings BL, Mukherjee K, Akroush W, Gonzalez FJ, Malik KU. 6β0Hydroxytestosterone, a metabolite of testosterone generated by CYP1B1, contributes to vascular changes in angiotensin II-induced hypertension in male mice. Biol Sex Differ. 2020;11(1):4. doi:10.1186/s13293-019-0280-4

68. Wang Y, Jin L. miRNA-145 is associated with spontaneous hypertension by targeting SLC7A1. Exp Ther Med. 2018;15(1):548–552. doi:10.3892/etm.2017.5371

69. Liu L, Wang C, Mi Y, Liu D, Li L, Fan J, Nan L, Jia N, Du Y. Association of MYH9 Polymorphisms with Hypertension in Patients with Chronic Kidney Disease in China. Kidney Blood Press Res. 2016;41(6):956–965. doi:10.1159/000452597

70. Pinho MJ, Gomes P, Serrão MP, Bonifácio MJ, Soares-da-Silva P. Organ-specific overexpression of renal LAT2 and enhanced tubular L-DOPA uptake precede the onset of hypertension. Hypertension. 2003;42(4):613–618. doi:10.1161/01.HYP.0000091822.00166.B1

71. Huang X, Wang B, Yang D, Shi X, Hong J, Wang S, Dai X, Zhou X, Geng YJ. Reduced expression of FXYD domain containing ion transport regulator 5 in association with hypertension. Int J Mol Med. 2012;29(2):231–238. doi:10.3892/ijmm.2011.837

72. Zhang S, Liu J, Zheng K, Chen L, Sun Y, Yao Z, Sun Y, Lin Y, Lin K, Yuan L. Exosomal miR-211 contributes to pulmonary hypertension via attenuating CaMK1/PPAR-γ axis. Vascul Pharmacol. 2021;136:106820. a doi:10.1016/j.vph.2020.106820

73. Zhao H, Wang Y, Zhang X, Guo Y, Wang X. miR-181b-5p inhibits endothelial-mesenchymal transition in monocrotaline-induced pulmonary arterial hypertension by targeting endocan and TGFBR1. Toxicol Appl Pharmacol. 2020;386:114827. doi:10.1016/j.taap.2019.114827

74. Li M, Mulkey F, Jiang C, O’Neil BH, Schneider BP, Shen F, Friedman PN, Momozawa Y, Kubo M, Niedzwiecki D, et al. Identification of a Genomic Region between SLC29A1 and HSP90AB1 Associated with Risk of Bevacizumab-Induced Hypertension: CALGB 80405 (Alliance). Clin Cancer Res. 2018;24(19):4734–4744. doi:10.1158/1078-0432.CCR-17-1523

75. Endres BT, Priestley JR, Palygin O, Flister MJ, Hoffman MJ, Weinberg BD, Grzybowski M, Lombard JH, Staruschenko A, Moreno C, et al. Mutation of Plekha7 attenuates salt-sensitive hypertension in the rat. Proc Natl Acad Sci U S A. 2014;111(35):12817–12822. doi:10.1073/pnas.1410745111

76. Wakui H, Uneda K, Tamura K, Ohsawa M, Azushima K, Kobayashi R, Ohki K, Dejima T, Kanaoka T, Tsurumi-Ikeya Y, et al. Renal tubule angiotensin II type 1 receptor-associated protein promotes natriuresis and inhibits salt-sensitive blood pressure elevation. J Am Heart Assoc. 2015;4(3):e001594. doi:10.1161/JAHA.114.001594

77. Simino J, Sung YJ, Kume R, Schwander K, Rao DC. Gene-alcohol interactions identify several novel blood pressure loci including a promising locus near SLC16A9. Front Genet. 2013;4:277. doi:10.3389/fgene.2013.00277

78. Kamide K, Kokubo Y, Yang J, Matayoshi T, Inamoto N, Takiuchi S, Horio T, Miwa Y, Yoshii M, Tomoike H, et al. Association of genetic polymorphisms of ACADSB and COMT with human hypertension. J Hypertens. 2007;25(1):103–110. doi:10.1097/HJH.0b013e3280103a40

79. Rafikov R, McBride ML, Zemskova M, Kurdyukov S, McClain N, Niihori M, Langlais PR, Rafikova O. Inositol monophosphatase 1 as a novel interacting partner of RAGE in pulmonary hypertension. Am J Physiol Lung Cell Mol Physiol. 2019;316(3):L428–L444. doi:10.1152/ajplung.00393.2018

80. Støy J, Grarup N, Hørlyck A, Ibsen L, Rungby J, Poulsen PL, Brandslund I, Christensen C, Hansen T, Pedersen O, et al. Blood pressure levels in male carriers of Arg82Cys in CD300LG. PLoS One. 2014;9(10):e109646. doi:10.1371/journal.pone.0109646

81. Liu J, Ke X, Wang L, Zhang Y, Yang J. Deficiency of cold-inducible RNA-binding protein exacerbated monocrotaline-induced pulmonary artery hypertension through Caveolin1 and CAVIN1. J Cell Mol Med. 2021;25(10):4732–4743. doi:10.1111/jcmm.16437

82. Zhang W, Zhang S, Li ZF, Huang C, Ren S, Zhou R, Jiang A, Yang AN. Knockdown of PIK3R1 by shRNA inhibits the activity of the splenic macrophages associated with hypersplenism due to portal hypertension. Pathol Res Pract. 2010;206(11):760–767. doi:10.1016/j.prp.2010.07.009

83. Schneider TM, Eadon MT, Cooper-DeHoff RM, Cavanaugh KL, Nguyen KA, Arwood MJ, Tillman EM, Pratt VM, Dexter PR, McCoy AB, et al. Multi-Institutional Implementation of Clinical Decision Support for APOL1, NAT2, and YEATS4 Genotyping in Antihypertensive Management. J Pers Med. 2021;11(6):480. doi:10.3390/jpm11060480

84. Pouly D, Debonneville A, Ruffieux-Daidié D, Maillard M, Abriel H, Loffing J, Staub O. Mice carrying ubiquitin-specific protease 2 (Usp2) gene inactivation maintain normal sodium balance and blood pressure. Am J Physiol Renal Physiol. 2013;305(1):F21–F30. doi:10.1152/ajprenal.00012.2013

85. Wang J, Yang K, Yuan JX. NEDD9, a Hypoxia-upregulated Mediator for Pathogenic Platelet-Endothelial Cell Interaction in Pulmonary Hypertension. Am J Respir Crit Care Med. 2021;203(12):1455–1458. doi:10.1164/rccm.202101-0007ED

86. Wu L, Gao L, Zhao X, Zhang M, Wu J, Mi J. A new risk locus in CHCHD5 for hypertension and obesity in a Chinese child population: a cohort study. BMJ Open. 2017;7(9):e016241. doi:10.1136/bmjopen-2017-016241

87. Zee RYL, Rivera A, Inostroza Y, Ridker PM, Chasman DI, Romero JR. Gene Variation of Endoplasmic Reticulum Aminopeptidases 1 and 2, and Risk of Blood Pressure Progression and Incident Hypertension among 17,255 Initially Healthy Women. Int J Genomics. 2018;2018:2308585. doi:10.1155/2018/2308585

88. Marunouchi T, Inomata S, Sanbe A, Takagi N, Tanonaka K. Protective effect of geranylgeranylacetone via enhanced induction of HSPB1 and HSPB8 in mitochondria of the failing heart following myocardial infarction in rats. Eur J Pharmacol. 2014;730:140–147. doi:10.1016/j.ejphar.2014.02.037

89. Zhang Y, Li C, Meng H, Guo D, Zhang Q, Lu W, Wang Q, Wang Y, Tu P. BYD Ameliorates Oxidative Stress-Induced Myocardial Apoptosis in Heart Failure Post-Acute Myocardial Infarction via the P38 MAPK-CRYAB Signaling Pathway. Front Physiol. 2018;9:505. doi:10.3389/fphys.2018.00505

90. Schurgers LJ, Burgmaier M, Ueland T, Schutters K, Aakhus S, Hofstra L, Gullestad L, Aukrust P, Hellmich M, Narula J, et al. Circulating annexin A5 predicts mortality in patients with heart failure. J Intern Med. 2016;279(1):89–97. doi:10.1111/joim.12396

91. Ortlepp JR, Vesper K, Mevissen V, Schmitz F, Janssens U, Franke A, Hanrath P, Weber C, Zerres K, Hoffmann R. Chemokine receptor (CCR2) genotype is associated with myocardial infarction and heart failure in patients under 65 years of age. J Mol Med (Berl). 2003;81(6):363–367. doi:10.1007/s00109-003-0435-x

92. Mittmann C, Chung CH, Höppner G, Michalek C, Nose M, Schüler C, Schuh A, Eschenhagen T, Weil J, Pieske B, et al. Expression of ten RGS proteins in human myocardium: functional characterization of an upregulation of RGS4 in heart failure. Cardiovasc Res. 2002;55(4):778–786. doi:10.1016/s0008-6363(02)00459-5

93. Stojanov S, Dejaco C, Lohse P, Huss K, Duftner C, Belohradsky BH, Herold M, Schirmer M. Clinical and functional characterisation of a novel TNFRSF1A c.605T>A/V173D cleavage site mutation associated with tumour necrosis factor receptor-associated periodic fever syndrome (TRAPS), cardiovascular complications and excellent response to etanercept treatment. Ann Rheum Dis. 2008;67(9):1292–1298. doi:10.1136/ard.2007.079376

94. Duan Q, Chen C, Yang L, Li N, Gong W, Li S, Wang DW. MicroRNA regulation of unfolded protein response transcription factor XBP1 in the progression of cardiac hypertrophy and heart failure in vivo. J Transl Med. 2015;13:363. doi:10.1186/s12967-015-0725-4

95. Deng B, Wang JX, Hu XX, Duan P, Wang L, Li Y, Zhu QL. Nkx2.5 enhances the efficacy of mesenchymal stem cells transplantation in treatment heart failure in rats. Life Sci. 2017;182:65–72. doi:10.1016/j.lfs.2017.06.014

96. Heimerl M, Sieve I, Ricke-Hoch M, Erschow S, Battmer K, Scherr M, Hilfiker-Kleiner D. Neuraminidase-1 promotes heart failure after ischemia/reperfusion injury by affecting cardiomyocytes and invading monocytes/macrophages. Basic Res Cardiol. 2020;115(6):62. doi:10.1007/s00395-020-00821-z

97. Simeunovic D, Odanovic N, Pljesa-Ercegovac M, Radic T, Radovanovic S, Coric V, Milinkovic I, Matic M, Djukic T, Ristic A, et al. Glutathione Transferase P1 Polymorphism Might Be a Risk Determinant in Heart Failure. Dis Markers. 2019;2019:6984845. doi:10.1155/2019/6984845

98. Krakoff LR, Buccino RA, Spann JF Jr, De Champlain J. Cardiac catechol O-methyltransferase and monoamine oxidase activity in congestive heart failure. Am J Physiol. 1968;215(3):549–552. doi:10.1152/ajplegacy.1968.215.3.549

99. Su Q, Zhang P, Yu D, Wu Z, Li D, Shen F, Liao P, Yin G. Upregulation of miR-93 and inhibition of LIMK1 improve ventricular remodeling and alleviate cardiac dysfunction in rats with chronic heart failure by inhibiting RhoA/ROCK signaling pathway activation. Aging (Albany NY). 2019;11(18):7570–7586. doi:10.18632/aging.102272

100. Beghi S, Cavaliere F, Manfredini M, Ferrarese S, Corazzari C, Beghi C, Buschini A. Polymorphism rs7214723 in CAMKK1: a new genetic variant associated with cardiovascular diseases. Biosci Rep. 2021;41(7):BSR20210326. doi:10.1042/BSR20210326

101. Anzalone R, Corrao S, Lo Iacono M, Loria T, Corsello T, Cappello F, Di Stefano A, Giannuzzi P, Zummo G, Farina F, et al. Isolation and characterization of CD276+/HLA-E+ human subendocardial mesenchymal stem cells from chronic heart failure patients: analysis of differentiative potential and immunomodulatory markers expression. Stem Cells Dev. 2013;22(1):1–17. doi:10.1089/scd.2012.0402

102. Ito S, Asahina H, Yamaguchi N, Tomaru U, Hasegawa T, Hatanaka Y, Hatanaka KC, Taguchi H, Harada T, Ohira H, et al. A case of radio-insensitive SMARCA4-deficient thoracic undifferentiated carcinoma with severe right heart failure. Respir Med Case Rep. 2021;32:101364. doi:10.1016/j.rmcr.2021.101364

103. Zhai YJ, Liu P, He HR, Zheng XW, Wang Y, Yang QT, Dong YL, Lu J. The association of ADORA2A and ADORA2B polymorphisms with the risk and severity of chronic heart failure: a case-control study of a northern Chinese population. Int J Mol Sci. 2015;16(2):2732–2746. doi:10.3390/ijms16022732

104. Xia C, Dong R, Chen C, Wang H, Wang DW. Cardiomyocyte specific expression of Acyl-coA thioesterase 1 attenuates sepsis induced cardiac dysfunction and mortality. Biochem Biophys Res Commun. 2015;468(4):533–540. doi:10.1016/j.bbrc.2015.10.078

105. Yamaguchi M. Regulatory role of regucalcin in heart calcium signaling: Insight into cardiac failure (Review). Biomed Rep. 2014;2(3):303–308. doi:10.3892/br.2014.245

106. Zhao A, Shen J, Ding Y, Sheng M, Zuo M, Lv H, Wang J, Shen Y, Wang H, Sun L. Long-read sequencing identified a novel nonsense and a de novo missense of PPA2 in trans in a Chinese patient with autosomal recessive infantile sudden cardiac failure. Clin Chim Acta. 2021;519:163–171. doi:10.1016/j.cca.2021.03.029

107. Hou YS, Wang JZ, Shi S, Han Y, Zhang Y, Zhi JX, Xu C, Li FF, Wang GY, Liu SL. Identification of epigenetic factor KAT2B gene variants for possible roles in congenital heart diseases. Biosci Rep. 2020;40(4):BSR20191779. doi:10.1042/BSR20191779

108. Mora A, Davies AM, Bertrand L, Sharif I, Budas GR, Jovanović S, Mouton V, Kahn CR, Lucocq JM, Gray GA, et al. Deficiency of PDK1 in cardiac muscle results in heart failure and increased sensitivity to hypoxia. EMBO J. 2003;22(18):4666–4676. doi:10.1093/emboj/cdg469

109. Jaenisch RB, Bertagnolli M, Borghi-Silva A, Arena R, Lago PD. Respiratory Muscle Training Improves Diaphragm Citrate Synthase Activity and Hemodynamic Function in Rats with Heart Failure. Braz J Cardiovasc Surg. 2017;32(2):104–110. doi:10.21470/1678-9741-2017-0002

110. Li Q, Zhao Y, Wu G, Chen S, Zhou Y, Li S, Zhou M, Fan Q, Pu J, Hong K, et al. De Novo FGF12 (Fibroblast Growth Factor 12) Functional Variation Is Potentially Associated With Idiopathic Ventricular Tachycardia. J Am Heart Assoc. 2017;6(8):e006130. doi:10.1161/JAHA.117.006130

111. Cheng Y, Chao J, Dai D, Dai Y, Zhu D, Liu B. AQP4-knockout aggravation of isoprenaline-induced myocardial injury is mediated by p66Shc and endoplasmic reticulum stress. Clin Exp Pharmacol Physiol. 2017;44(11):1106–1115. doi:10.1111/1440-1681.12812

112. Pappas CT, Farman GP, Mayfield RM, Konhilas JP, Gregorio CC. Cardiac-specific knockout of Lmod2 results in a severe reduction in myofilament force production and rapid cardiac failure. J Mol Cell Cardiol. 2018;122:88–97. doi:10.1016/j.yjmcc.2018.08.009

113. Kühn EC, Slagman A, Kühn-Heid ECD, Seelig J, Schwiebert C, Minich WB, Stoppe C, Möckel M, Schomburg L. Circulating levels of selenium-binding protein 1 (SELENBP1) are associated with risk for major adverse cardiac events and death. J Trace Elem Med Biol. 2019;52:247–253. doi:10.1016/j.jtemb.2019.01.005

114. O’Brien PJ, O’Grady M, McCutcheon LJ, Shen H, Nowack L, Horne RD, Mirsalimi SM, Julian RJ, Grima EA, Moe GW, et al. Myocardial myoglobin deficiency in various animal models of congestive heart failure. J Mol Cell Cardiol. 1992;24(7):721–730. doi:10.1016/0022-2828(92)93386-x

115. Weber C, Neacsu I, Krautz B, Schlegel P, Sauer S, Raake P, Ritterhoff J, Jungmann A, Remppis AB, Stangassinger M, et al. Therapeutic safety of high myocardial expression levels of the molecular inotrope S100A1 in a preclinical heart failure model. Gene Ther. 2014;21(2):131–138. doi:10.1038/gt.2013.63

116. Dobrev D, Wehrens XH. Role of RyR2 phosphorylation in heart failure and arrhythmias: Controversies around ryanodine receptor phosphorylation in cardiac disease. Circ Res. 2014;114(8):1311–1319. doi:10.1161/CIRCRESAHA.114.300568

117. Arking DE, Reinier K, Post W, Jui J, Hilton G, O’Connor A, Prineas RJ, Boerwinkle E, Psaty BM, Tomaselli GF, et al. Genome-wide association study identifies GPC5 as a novel genetic locus protective against sudden cardiac arrest. PLoS One. 2010;5(3):e9879. doi:10.1371/journal.pone.0009879

118. Bovill E, Westaby S, Reji S, Sayeed R, Crisp A, Shaw T. Induction by left ventricular overload and left ventricular failure of the human Jumonji gene (JARID2) encoding a protein that regulates transcription and reexpression of a protective fetal program. J Thorac Cardiovasc Surg. 2008;136(3):709–716. doi:10.1016/j.jtcvs.2008.02.020

119. Watanabe H, Ichihara E, Kano H, Ninomiya K, Tanimoto M, Kiura K. Congestive Heart Failure During Osimertinib Treatment for Epidermal Growth Factor Receptor (EGFR)-mutant Non-small Cell Lung Cancer (NSCLC). Intern Med. 2017;56(16):2195–2197. doi:10.2169/internalmedicine.8344-16

120. Li W, Yin L, Sun X, Wu J, Dong Z, Hu K, Sun A, Ge J.Alpha-lipoic acid protects against pressure overload-induced heart failure via ALDH2-dependent Nrf1-FUNDC1 signaling. Cell Death Dis. 2020;11(7):599. doi:10.1038/s41419-020-02805-2

121. Cannavo A, Rengo G, Liccardo D, Pagano G, Zincarelli C, De Angelis MC, Puglia R, Di Pietro E, Rabinowitz JE, Barone MV, et al. β adrenergic receptor and sphingosine-1-phosphate receptor 1 (S1PR1) reciprocal downregulation influences cardiac hypertrophic response and progression to heart failure: protective role of S1PR1 cardiac gene therapy. Circulation. 2013;128(15):1612–1622. doi:10.1161/CIRCULATIONAHA.113.002659

122. Heaton MP, Bassett AS, Whitman KJ, Krafsur GM, Lee SI, Carlson JM, Clark HJ, Smith HR, Pelster MC, Basnayake V, et al. Evaluation of EPAS1 variants for association with bovine congestive heart failure. F1000Res. 2019;8:1189. doi:10.12688/f1000research.19951.1

123. Bouchard L, Faucher G, Tchernof A, Deshaies Y, Marceau S, Lescelleur O, Biron S, Bouchard C, Pérusse L, Vohl MC. Association of OSBPL11 gene polymorphisms with cardiovascular disease risk factors in obesity. Obesity (Silver Spring). 2009;17(7):1466–1472. doi:10.1038/oby.2009.71

124. Frustaci A, De Luca A, Galea N, Verardo R, Guida V, Carrozzo R, Chimenti C, Frustaci E, Sansone L, Russo MA. Novel dilated cardiomyopathy associated to Calreticulin and Myo7A gene mutation in Usher syndrome. ESC Heart Fail. 2021;8(3):2310–2315. doi:10.1002/ehf2.13260

125. Zhou H, Lei X, Yan Y, Lydic T, Li J, Weintraub NL, Su H, Chen W. Targeting ATGL to rescue BSCL2 lipodystrophy and its associated cardiomyopathy. JCI Insight. 2019;5(14):e129781. doi:10.1172/jci.insight.129781

126. Miura A, Kondo H, Yamamoto T, Okumura Y, Nishio H. Sudden Unexpected Death of Infantile Dilated Cardiomyopathy with JPH2 and PKD1 Gene Variants. Int Heart J. 2020;61(5):1079–1083. doi:10.1536/ihj.20-155

127. Gong F, Gu J, Wang H. Up regulated Tmbim1 activation promotes high fat diet (HFD)-induced cardiomyopathy by enhancement of inflammation and oxidative stress. Biochem Biophys Res Commun. 2018;504(4):797–804. doi:10.1016/j.bbrc.2018.08.059

128. Watanabe K, Arumugam S, Sreedhar R, Thandavarayan RA, Nakamura T, Nakamura M, Harima M, Yoneyama H, Suzuki K. Small interfering RNA therapy against carbohydrate sulfotransferase 15 inhibits cardiac remodeling in rats with dilated cardiomyopathy. Cell Signal. 2015;27(7):1517–1524. doi:10.1016/j.cellsig.2015.03.004

129. Shishido A, Morisada N, Tominaga K, Uemura H, Haruna A, Hanafusa H, Nozu K, Iijima K. Japanese boy with NAA10-related syndrome and hypertrophic cardiomyopathy. Hum Genome Var. 2020;7:23. doi:10.1038/s41439-020-00110-0

130. Su D, Ju Y, Han W, Yang Y, Wang F, Wang T, Tang J. Tcf3-activated lncRNA Gas5 regulates newborn mouse cardiomyocyte apoptosis in diabetic cardiomyopathy. J Cell Biochem. 2020;121(11):4337–4346. doi:10.1002/jcb.29630

131. Lu D, Zhang L, Bao D, Lu Y, Zhang X, Liu N, Ge W, Gao X, Li H, Zhang L. Calponin1 inhibits dilated cardiomyopathy development in mice through the ε PKC pathway. Int J Cardiol. 2014;173(2):146–153. doi:10.1016/j.ijcard.2014.02.032

132. Long PA, Theis JL, Shih YH, Maleszewski JJ, Abell Aleff PC, Evans JM, Xu X, Olson TM. Recessive TAF1A mutations reveal ribosomopathy in siblings with end-stage pediatric dilated cardiomyopathy. Hum Mol Genet. 2017;26(15):2874–2881. doi:10.1093/hmg/ddx169

133. Jacobi-Polishook T, Yosha-Orpaz N, Sagi Y, Lev D, Lerman-Sagie T. Successful pregnancy in a patient with mitochondrial cardiomyopathy due to ACAD9 deficiency. JIMD Rep. 2020;56(1):9–13. doi:10.1002/jmd2.12157

134. Hedberg-Oldfors C, Abramsson A, Osborn DPS, Danielsson O, Fazlinezhad A, Nilipour Y, Hübbert L, Nennesmo I, Visuttijai K, Bharj J, et al. Cardiomyopathy with lethal arrhythmias associated with inactivation of KLHL24. Hum Mol Genet. 2019;28(11):1919–1929. doi:10.1093/hmg/ddz032

135. Auxerre-Plantié E, Nielsen T, Grunert M, Olejniczak O, Perrot A, Özcelik C, Harries D, Matinmehr F, Dos Remedios C, Mühlfeld C, Kraft T, et al. Identification of MYOM2 as a candidate gene in hypertrophic cardiomyopathy and Tetralogy of Fallot, and its functional evaluation in the Drosophila heart. Dis Model Mech. 2020;13(12):dmm045377. doi:10.1242/dmm.045377

136. Salazar-Mendiguchía J, Ochoa JP, Palomino-Doza J, Domínguez F, Díez-López C, Akhtar M, Ramiro-León S, Clemente MM, Pérez-Cejas A, Robledo M, et al. Mutations in TRIM63 cause an autosomal-recessive form of hypertrophic cardiomyopathy. Heart. 2020;106(17):1342–1348. doi:10.1136/heartjnl-2020-316913

137. Janssens B, Mohapatra B, Vatta M, Goossens S, Vanpoucke G, Kools P, Montoye T, van Hengel J, Bowles NE, van Roy F, et al. Assessment of the CTNNA3 gene encoding human alpha T-catenin regarding its involvement in dilated cardiomyopathy. Hum Genet. 2003;112(3):227–236. doi:10.1007/s00439-002-0857-5

138. Wang B, Wu Y, Ge Z, Zhang X, Yan Y, Xie Y. NLRC5 deficiency ameliorates cardiac fibrosis in diabetic cardiomyopathy by regulating EndMT through Smad2/3 signaling pathway. Biochem Biophys Res Commun. 2020;528(3):545–553. doi:10.1016/j.bbrc.2020.05.151

139. Zhang WY, Wang J, Li AZ. A study of the effects of SGLT-2 inhibitors on diabetic cardiomyopathy through miR-30d/KLF9/VEGFA pathway. Eur Rev Med Pharmacol Sci. 2020;24(11):6346–6359. doi:10.26355/eurrev_202006_21533

140. Williams JL, Paudyal A, Awad S, Nicholson J, Grzesik D, Botta J, Meimaridou E, Maharaj AV, Stewart M, Tinker A, et al. Mylk3 null C57BL/6N mice develop cardiomyopathy, whereas Nnt null C57BL/6J mice do not. Life Sci Alliance. 2020;3(4):e201900593. doi:10.26508/lsa.201900593

141. Lennermann D, Backs J, van den Hoogenhof MMG. New Insights in RBM20 Cardiomyopathy. Curr Heart Fail Rep. 2020;17(5):234–246. doi:10.1007/s11897-020-00475-x

142. Sasagawa S, Nishimura Y, Okabe S, Murakami S, Ashikawa Y, Yuge M, Kawaguchi K, Kawase R, Okamoto R, Ito M, et al. Downregulation of GSTK1 Is a Common Mechanism Underlying Hypertrophic Cardiomyopathy. Front Pharmacol. 2016;7:162. doi:10.3389/fphar.2016.00162

143. Gusic M, Schottmann G, Feichtinger RG, Du C, Scholz C, Wagner M, Mayr JA, Lee CY, Yépez VA, Lorenz N, et al. Bi-Allelic UQCRFS1 Variants Are Associated with Mitochondrial Complex III Deficiency, Cardiomyopathy, and Alopecia Totalis. Am J Hum Genet. 2020;106(1):102–111. doi:10.1016/j.ajhg.2019.12.005

144. Loeffen J, Elpeleg O, Smeitink J, Smeets R, Stöckler-Ipsiroglu S, Mandel H, Sengers R, Trijbels F, van den Heuvel L. Mutations in the complex I NDUFS2 gene of patients with cardiomyopathy and encephalomyopathy. Ann Neurol. 2001;49(2):195–201. doi:10.1002/1531-8249(20010201)49:2<195::aid-ana39>3.0.co;2-m

145. Abdulhag UN, Soiferman D, Schueler-Furman O, Miller C, Shaag A, Elpeleg O, Edvardson S, Saada A. Mitochondrial complex IV deficiency, caused by mutated COX6B1, is associated with encephalomyopathy, hydrocephalus and cardiomyopathy. Eur J Hum Genet. 2015;23(2):159–164. doi:10.1038/ejhg.2014.85

146. Domarkienė I, Pranculis A, Germanas S, Jakaitienė A, Vitkus D, Dženkevičiūtė V, Kučinskienė Z, Kučinskas V. RTN4 and FBXL17 Genes are Associated with Coronary Heart Disease in Genome-Wide Association Analysis of Lithuanian Families. Balkan J Med Genet. 2013;16(2):17–22. doi:10.2478/bjmg-2013-0026

147. Mo XG, Liu W, Yang Y, Imani S, Lu S, Dan G, Nie X, Yan J, Zhan R, Li X, et al. NCF2, MYO1F, S1PR4, and FCN1 as potential noninvasive diagnostic biomarkers in patients with obstructive coronary artery: A weighted gene co-expression network analysis. J Cell Biochem. 2019;120(10):18219–18235. doi:10.1002/jcb.29128

148. Takefuji M, Asano H, Mori K, Amano M, Kato K, Watanabe T, Morita Y, Katsumi A, Itoh T, Takenawa T, et al. Mutation of ARHGAP9 in patients with coronary spastic angina. J Hum Genet. 2010;55(1):42–49. doi:10.1038/jhg.2009.120

149. Vergeer M, Cohn DM, Boekholdt SM, Sandhu MS, Prins HM, Ricketts SL, Wareham NJ, Kastelein JJ, Khaw KT, Kamphuisen PW, et al. Lack of association between common genetic variation in endothelial lipase (LIPG) and the risk for CAD and DVT. Atherosclerosis. 2010;211(2):558–564. doi:10.1016/j.atherosclerosis.2010.04.004

150. Prasongsukarn K, Dechkhajorn W, Benjathummarak S, Maneerat Y. TRPM2, PDLIM5, BCL3, CD14, GBA Genes as Feasible Markers for Premature Coronary Heart Disease Risk. Front Genet. 2021;12:598296. doi:10.3389/fgene.2021.598296

151. Cai H, Wang XL, Wilcken DE. Genetic polymorphism of heparan sulfate proteoglycan (perlecan, HSPG2), lipid profiles and coronary artery disease in the Australian population. Atherosclerosis. 2000;148(1):125–129. doi:10.1016/s0021-9150(99)00213-0

152. Castrechini S, Rivasi AP. Apolipoprotein B, apolipoprotein E, and angiotensin-converting enzyme polymorphisms in 2 Italian populations at different risk for coronary artery disease and comparison of allele frequencies among European populations. Hum Biol. 1999;71(6):933–945

153. Grinshtein YI, Kosinova AA, Grinshtein IY, Subbotina TN, Savchenko AA. The Prognostic Value of Combinations of Genetic Polymorphisms in the ITGB3, ITGA2, and CYP2C19*2 Genes in Predicting Cardiovascular Outcomes After Coronary Bypass Grafting. Genet Test Mol Biomarkers. 2018;22(4):259–265. doi:10.1089/gtmb.2017.0177

154. Vinitha A, Kutty VR, Vivekanand A, Reshmi G, Divya G, Sumi S, Santosh KR, Pratapachandran NS, Ajit MS, Kartha CC, et al. PPIA rs6850: A > G single-nucleotide polymorphism is associated with raised plasma cyclophilin A levels in patients with coronary artery disease. Mol Cell Biochem. 2016;412(1-2):259–268. doi:10.1007/s11010-015-2632-7

155. Ramkaran P, Khan S, Phulukdaree A, Moodley D, Chuturgoon AA. miR-146a polymorphism influences levels of miR-146a, IRAK-1, and TRAF-6 in young patients with coronary artery disease. Cell Biochem Biophys. 2014;68(2):259–266. doi:10.1007/s12013-013-9704-7

156. Watzka M, Nebel A, El Mokhtari NE, Ivandic B, Müller J, Schreiber S, Oldenburg J. Functional promoter polymorphism in the VKORC1 gene is no major genetic determinant for coronary heart disease in Northern Germans. Thromb Haemost. 2007;97(6):998–1002.

157. Stec A, Ksiazek A, Buraczynska M. Rs10887800 renalase gene polymorphism is associated with an increased risk of coronary artery disease in hemodialyzed patients. Int Urol Nephrol. 2016;48(6):871–876. doi:10.1007/s11255-016-1270-7

158. Zidi I, Kharrat N, Abdelhedi R, Hassine AB, Laaribi AB, Yahia HB, Abdelmoula NB, Abid L, Rebai A, Rizzo R. Nonclassical human leukocyte antigen (HLA-G, HLA-E, and HLA-F) in coronary artery disease. Hum Immunol. 2016;77(4):325–329. doi:10.1016/j.humimm.2016.01.008

159. Domarkien I, Pranculis A, Germanas S, Jakaitienė A, Vitkus D, Dženkevičiūtė V, Kučinskienė Z, Kučinskas V. RTN4 and FBXL17 Genes are Associated with Coronary Heart Disease in Genome-Wide Association Analysis of Lithuanian Families. Balkan J Med Genet. 2013;16(2):17–22. doi:10.2478/bjmg-2013-0026

160. Sheu JJ, Lin YJ, Chang JS, Wan L, Chen SY, Huang YC, Chan C, Chiu IW, Tsai FJ. Association of COL11A2 polymorphism with susceptibility to Kawasaki disease and development of coronary artery lesions. Int J Immunogenet. 2010;37(6):487–492. doi:10.1111/j.1744-313X.2010.00952.x

161. Raffa S, Chin XLD, Stanzione R, Forte M, Bianchi F, Cotugno M, Marchitti S, Micaloni A, Gallo G, Schirone L, et al. The reduction of NDUFC2 expression is associated with mitochondrial impairment in circulating mononuclear cells of patients with acute coronary syndrome. Int J Cardiol. 2019;286:127–133. doi:10.1016/j.ijcard.2019.02.027

162. Chen C, Peng H, Zeng Y, Dong G. CD14, CD163, and CCR1 are involved in heart and blood communication in ischemic cardiac diseases. J Int Med Res. 2020;48(9):300060520951649. doi:10.1177/0300060520951649

163. Tadimalla A, Belmont PJ, Thuerauf DJ, Glassy MS, Martindale JJ, Gude N, Sussman MA, Glembotski CC. Mesencephalic astrocyte-derived neurotrophic factor is an ischemia-inducible secreted endoplasmic reticulum stress response protein in the heart. Circ Res. 2008;103(11):1249–1258. doi:10.1161/CIRCRESAHA.108.180679

164. Zhu WS, Guo W, Zhu JN, Tang CM, Fu YH, Lin QX, Tan N, Shan ZX. Hsp90aa1: a novel target gene of miR-1 in cardiac ischemia/reperfusion injury. Sci Rep. 2016;6:24498. doi:10.1038/srep24498

165. Wang Y, Jin L, Song Y, Zhang M, Shan D, Liu Y, Fang M, Lv F, Xiao RP, Zhang Y. β-arrestin 2 mediates cardiac ischemia-reperfusion injury via inhibiting GPCR-independent cell survival signalling. Cardiovasc Res. 2017;113(13):1615–1626. doi:10.1093/cvr/cvx147

166. Wang J, Cheng X, Zhao H, Yang Q, Xu Z. Downregulation of the zinc transporter SLC39A13 (ZIP13) is responsible for the activation of CaMKII at reperfusion and leads to myocardial ischemia/reperfusion injury in mouse hearts. J Mol Cell Cardiol. 2021;152:69–79. doi:10.1016/j.yjmcc.2020.12.002

167. Natarajan R, Salloum FN, Fisher BJ, Ownby ED, Kukreja RC, Fowler AA. Activation of hypoxia-inducible factor-1 via prolyl-4 hydoxylase-2 gene silencing attenuates acute inflammatory responses in postischemic myocardium. Am J Physiol Heart Circ Physiol. 2007;293(3):H1571–H1580. doi:10.1152/ajpheart.00291.2007

168. Zhou W, Zhong Z, Lin D, Liu Z, Zhang Q, Xia H, Peng S, Liu A, Lu Z, Wang Y, Ye S, et al. Hypothermic oxygenated perfusion inhibits HECTD3-mediated TRAF3 polyubiquitination to alleviate DCD liver ischemia-reperfusion injury. Cell Death Dis. 2021;12(2):211. doi:10.1038/s41419-021-03493-2

169. Cui Y, Wang Y, Liu G. Protective effect of Barbaloin in a rat model of myocardial ischemia reperfusion injury through the regulation of the CNPY2 PERK pathway. Int J Mol Med. 2019;43(5):2015–2023. doi:10.3892/ijmm.2019.4123

170. Du J, Li Z, Li QZ, Guan T, Yang Q, Xu H, Pritchard KA, Camara AK, Shi Y. Enoyl coenzyme a hydratase domain-containing 2, a potential novel regulator of myocardial ischemia injury. J Am Heart Assoc. 2013;2(5):e000233. doi:10.1161/JAHA.113.000233

171. Zhang H, Gong G, Wang P, Zhang Z, Kolwicz SC, Rabinovitch PS, Tian R, Wang W. Heart specific knockout of Ndufs4 ameliorates ischemia reperfusion injury. J Mol Cell Cardiol. 2018;123:38–45. doi:10.1016/j.yjmcc.2018.08.022

172. Tombo N, Imam Aliagan AD, Feng Y, Singh H, Bopassa JC. Cardiac ischemia/reperfusion stress reduces inner mitochondrial membrane protein (mitofilin) levels during early reperfusion. Free Radic Biol Med. 2020;158:181–194. doi:10.1016/j.freeradbiomed.2020.06.039

173. Savastano S, Valentino R, Di Somma C, Orio F, Pivonello C, Passaretti F, Brancato V, Formisano P, Colao A, Beguinot F, et al. Serum 25-Hydroxyvitamin D Levels, phosphoprotein enriched in diabetes gene product (PED/PEA-15) and leptin-to-adiponectin ratio in women with PCOS. Nutr Metab (Lond). 2011;8:84. doi:10.1186/1743-7075-8-84

174. Ramos-Lopez E, Ghebru S, Van Autreve J, Aminkeng F, Herwig J, Seifried E, Seidl C, Van der Auwera B, Badenhoop K. Neither an intronic CA repeat within the CD48 gene nor the HERV-K18 polymorphisms are associated with type 1 diabetes. Tissue Antigens. 2006;68(2):147–152. doi:10.1111/j.1399-0039.2006.00637.x

175. Alhaidan Y, Christesen HT, Højlund K, Al Balwi MA, Brusgaard K. A novel gene in early childhood diabetes: EDEM2 silencing decreases SLC2A2 and PXD1 expression, leading to impaired insulin secretion. Mol Genet Genomics. 2020;295(5):1253–1262. doi:10.1007/s00438-020-01695-5

176. Ma XW, Chang ZW, Qin MZ, Sun Y, Huang HL, He Y. Decreased expression of complement regulatory proteins, CD55 and CD59, on peripheral blood leucocytes in patients with type 2 diabetes and macrovascular diseases. Chin Med J (Engl). 2009;122(18):2123–2128.

177. Cividini F, Scott BT, Suarez J, Casteel DE, Heinz S, Dai A, Diemer T, Suarez JA, Benner CW, Ghassemian M, Dillmann WH. Ncor2/PPARα - Dependent Upregulation of MCUb in the Type 2 Diabetic Heart Impacts Cardiac Metabolic Flexibility and Function. Diabetes. 2021;70(3):665–679. doi:10.2337/db20-0779

178. Ren Q, Xiao J, Han X, Yang W, Ji L. Impact of variants of the EXT2 gene on Type 2 diabetes and its related traits in the Chinese han population. Endocr Res. 2015;40(2):79–82. doi:10.3109/07435800.2014.952015

179. Meng S, Cao JT, Zhang B, Zhou Q, Shen CX, Wang CQ. Downregulation of microRNA-126 in endothelial progenitor cells from diabetes patients, impairs their functional properties, via target gene Spred-1. J Mol Cell Cardiol. 2012;53(1):64–72. doi:10.1016/j.yjmcc.2012.04.003

180. Al-Fadhli FM, Afqi M, Sairafi MH, Almuntashri M, Alharby E, Alharbi G, Abdud Samad F, Hashmi JA, Zaytuni D, Bahashwan AA, et al. Biallelic loss of function variant in the unfolded protein response gene PDIA6 is associated with asphyxiating thoracic dystrophy and neonatal-onset diabetes. Clin Genet. 2021;99(5):694–703. doi:10.1111/cge.13930

181. Haseda F, Imagawa A, Nishikawa H, Mitsui S, Tsutsumi C, Fujisawa R, Sano H, Murase-Mishiba Y, Terasaki J, Sakaguchi S, et al. Antibody to CMRF35-Like Molecule 2, CD300e A Novel Biomarker Detected in Patients with Fulminant Type 1 Diabetes. PLoS One. 2016;11(8):e0160576. doi:10.1371/journal.pone.0160576

182. Cheung YH, Watkinson J, Anastassiou D. Conditional meta-analysis stratifying on detailed HLA genotypes identifies a novel type 1 diabetes locus around TCF19 in the MHC. Hum Genet. 2011;129(2):161–176. doi:10.1007/s00439-010-0908-2

183. Xia W, Pessentheiner AR, Hofer DC, Amor M, Schreiber R, Schoiswohl G, Eichmann TO, Walenta E, Itariu B, Prager G, et al. Loss of ABHD15 Impairs the Anti-lipolytic Action of Insulin by Altering PDE3B Stability and Contributes to Insulin Resistance. Cell Rep. 2018;23(7):1948–1961. doi:10.1016/j.celrep.2018.04.055

184. Gao T, Qian S, Shen S, Zhang X, Liu J, Jia W, Chen Z, Ye J. Reduction of mitochondrial 3-oxoacyl-ACP synthase (OXSM) by hyperglycemia is associated with deficiency of α-lipoic acid synthetic pathway in kidney of diabetic mice. Biochem Biophys Res Commun. 2019;512(1):106–111. doi:10.1016/j.bbrc.2019.02.155

185. Thameem F, Yang X, Permana PA, Wolford JK, Bogardus C, Prochazka M. Evaluation of the microsomal glutathione S-transferase 3 (MGST3) locus on 1q23 as a Type 2 diabetes susceptibility gene in Pima Indians. Hum Genet. 2003;113(4):353–358. doi:10.1007/s00439-003-0980-y

186. Shigeta Y, Izumi K, Shichiri M. Influence of the administration of coenzyme Q7 on abnormality of vibratory perception in diabetics. J Vitaminol (Kyoto). 1965;11(3):199–203. doi:10.5925/jnsv1954.11.199

187. Xia Q, Chesi A, Manduchi E, Johnston BT, Lu S, Leonard ME, Parlin UW, Rappaport EF, Huang P, Wells AD, et al. The type 2 diabetes presumed causal variant within TCF7L2 resides in an element that controls the expression of ACSL5. Diabetologia. 2016;59(11):2360–2368. doi:10.1007/s00125-016-4077-2

188. Sun L, Zhang X, Wang T, Chen M, Qiao H. Association of ANK1 variants with new-onset type 2 diabetes in a Han Chinese population from northeast China. Exp Ther Med. 2017;14(4):3184–3190. doi:10.3892/etm.2017.4866

189. Baker DJ, Timmons JA, Greenhaff PL. Glycogen phosphorylase inhibition in type 2 diabetes therapy: a systematic evaluation of metabolic and functional effects in rat skeletal muscle. Diabetes. 2005;54(8):2453–2459. doi:10.2337/diabetes.54.8.2453

190. Zhang L, Chen Z, Wang Y, Tweardy DJ, Mitch WE. Stat3 activation induces insulin resistance via a muscle-specific E3 ubiquitin ligase Fbxo40. Am J Physiol Endocrinol Metab. 2020;318(5):E625–E635. doi:10.1152/ajpendo.00480.2019

191. Esteves JV, Yonamine CY, Machado UF. SLC2A4 expression and its epigenetic regulation as biomarkers for insulin resistance treatment in diabetes mellitus. Biomark Med. 2020;14(6):413–416. doi:10.2217/bmm-2019-0481

192. Santin I, Castellanos-Rubio A, Aransay AM, Gutierrez G, Gaztambide S, Rica I, Vicario JL, Noble JA, Castaño L, Bilbao JR. Exploring the diabetogenicity of the HLA-B18-DR3 CEH: independent association with T1D genetic risk close to HLA-DOA. Genes Immun. 2009;10(6):596–600. doi:10.1038/gene.2009.41

193. Penfornis A, Tuomilehto-Wolf E, Faustman DL, Hitman GA; DiMe (Childhood Diabetes in Finland) Study Group. Analysis of TAP2 polymorphisms in Finnish individuals with type I diabetes. Hum Immunol. 2002;63(1):61–70. doi:10.1016/s0198-8859(01)00365-2

194. Varney MD, Valdes AM, Carlson JA, Noble JA, Tait BD, Bonella P, Lavant E, Fear AL, Louey A, Moonsamy P, et al. HLA DPA1, DPB1 alleles and haplotypes contribute to the risk associated with type 1 diabetes: analysis of the type 1 diabetes genetics consortium families. Diabetes. 2010;59(8):2055–2062. doi:10.2337/db09-0680

195. Payne F, Colnaghi R, Rocha N, Seth A, Harris J, Carpenter G, Bottomley WE, Wheeler E, Wong S, Saudek V, et al. Hypomorphism in human NSMCE2 linked to primordial dwarfism and insulin resistance. J Clin Invest. 2014;124(9):4028–4038. doi:10.1172/JCI73264

196. Yagil C, Varadi-Levi R, Yagil Y. A novel mutation in the NADH dehydrogenase (ubiquinone) 1 alpha subcomplex 4 (Ndufa4) gene links mitochondrial dysfunction to the development of diabetes in a rodent model. Dis Model Mech. 2018;11(11):dmm036699. doi:10.1242/dmm.036699

197. Huang T, Wang L, Bai M, Zheng J, Yuan D, He Y, Wang Y, Jin T, Cui W. Influence of IGF2BP2, HMG20A, and HNF1B genetic polymorphisms on the susceptibility to Type 2 diabetes mellitus in Chinese Han population. Biosci Rep. 2020;40(5):BSR20193955. doi:10.1042/BSR20193955

198. Arendt M, Fall T, Lindblad-Toh K, Axelsson E. Amylase activity is associated with AMY2B copy numbers in dog: implications for dog domestication, diet and diabetes. Anim Genet. 2014;45(5):716–722. doi:10.1111/age.12179

199. AlDehaini DMB, Al-Bustan SA, Malalla ZHA, Ali ME, Sater M, Giha HA. The influence of TERC, TERT and ACYP2 genes polymorphisms on plasma telomerase concentration, telomeres length and T2DM. Gene. 2021;766:145127. doi:10.1016/j.gene.2020.145127

200. Jiang DS, Zhang XF, Gao L, Zong J, Zhou H, Liu Y, Zhang Y, Bian ZY, Zhu LH, Fan GC, et al. Signal regulatory protein-αprotects against cardiac hypertrophy via the disruption of toll-like receptor 4 signaling. Hypertension. 2014;63(1):96–104. doi:10.1161/HYPERTENSIONAHA.113.01506

201. Yang D, Wang Y, Jiang M, Deng X, Pei Z, Li F, Xia K, Zhu L, Yang T, Chen M. Downregulation of Profilin-1 Expression Attenuates Cardiomyocytes Hypertrophy and Apoptosis Induced by Advanced Glycation End Products in H9c2 Cells. Biomed Res Int. 2017;2017:9716087. doi:10.1155/2017/9716087

202. Romano N, Ricciardi S, Gallo P, Ceci M. Upregulation of eIF6 inhibits cardiac hypertrophy induced by phenylephrine. Biochem Biophys Res Commun. 2018;495(1):601–606. doi:10.1016/j.bbrc.2017.11.046

203. Lund AK, Goens MB, Nuñez BA, Walker MK. Characterizing the role of endothelin-1 in the progression of cardiac hypertrophy in aryl hydrocarbon receptor (AhR) null mice. Toxicol Appl Pharmacol. 2006;212(2):127–135. doi:10.1016/j.taap.2005.07.005

204. Zhang D, Liang C, Li P, Yang L, Hao Z, Kong L, Tian X, Guo C, Dong J, Zhang Y, et al. Runt-related transcription factor 1 (Runx1) aggravates pathological cardiac hypertrophy by promoting p53 expression. J Cell Mol Med. 2021;10.1111/jcmm.16704. doi:10.1111/jcmm.16704

205. Liu Y, Li S, Gao Z, Li S, Tan Q, Li Y, Wang D, Wang Q. Indoleamine 2,3-Dioxygenase 1 (IDO1) Promotes Cardiac Hypertrophy via a PI3K-AKT-mTOR-Dependent Mechanism. Cardiovasc Toxicol. 2021;21(8):655–668. doi:10.1007/s12012-021-09657-y

206. Mitra A, Basak T, Ahmad S, Datta K, Datta R, Sengupta S, Sarkar S. Comparative Proteome Profiling during Cardiac Hypertrophy and Myocardial Infarction Reveals Altered Glucose Oxidation by Differential Activation of Pyruvate Dehydrogenase E1 Component Subunit β J Mol Biol. 2015;427(11):2104–2120. doi:10.1016/j.jmb.2014.10.026

207. Zou R, Tao J, Qiu J, Shi W, Zou M, Chen W, Li W, Zhou N, Wang S, Ma L, et al. Ndufs1 Deficiency Aggravates the Mitochondrial Membrane Potential Dysfunction in Pressure Overload-Induced Myocardial Hypertrophy. Oxid Med Cell Longev. 2021;2021:5545261. doi:10.1155/2021/5545261

208. Ito S, Asakura M, Liao Y, Min KD, Takahashi A, Shindo K, Yamazaki S, Tsukamoto O, Asanuma H, Mogi M, et al. Identification of the Mtus1 Splice Variant as a Novel Inhibitory Factor Against Cardiac Hypertrophy. J Am Heart Assoc. 2016;5(7):e003521. doi:10.1161/JAHA.116.003521

209. Pan L, Sheng M, Huang Z, Zhu Z, Xu C, Teng L, He L, Gu C, Yi C, Li J. Zinc-finger protein 418 overexpression protects against cardiac hypertrophy and fibrosis. PLoS One. 2017;12(10):e0186635. doi:10.1371/journal.pone.0186635

210. Zhang JL, Zhang DH, Li YP, Wu LM, Liang C, Yao R, Wang Z, Feng SD, Wang ZM, Zhang YZ. Myotubularin-related protein 14 suppresses cardiac hypertrophy by inhibiting Akt. Cell Death Dis. 2020;11(2):140. doi:10.1038/s41419-020-2330-6

211. Zhu J, Xu Y, Ren G, Hu X, Wang C, Yang Z, Li Z, Mao W, Lu D. Tanshinone IIA Sodium sulfonate regulates antioxidant system, inflammation, and endothelial dysfunction in atherosclerosis by downregulation of CLIC1. Eur J Pharmacol. 2017;815:427–436. doi:10.1016/j.ejphar.2017.09.047

212. Li J, Chen Y, Gao J, Chen Y, Zhou C, Lin X, Liu C, Zhao M, Xu Y, Ji L, et al. Eva1a ameliorates atherosclerosis by promoting re-endothelialization of injured arteries via Rac1/Cdc42/Arpc1b. Cardiovasc Res. 2021;117(2):450–461. doi:10.1093/cvr/cvaa011

213. Bandaru S, Ala C, Salimi R, Akula MK, Ekstrand M, Devarakonda S, Karlsson J, Van den Eynden J, Bergström G, Larsson E, et al. Targeting Filamin A Reduces Macrophage Activity and Atherosclerosis. Circulation. 2019;140(1):67–79. doi:10.1161/CIRCULATIONAHA.119.039697

214. Maier A, Wu H, Cordasic N, Oefner P, Dietel B, Thiele C, Weidemann A, Eckardt KU, Warnecke C. Hypoxia-inducible protein 2 Hig2/Hilpda mediates neutral lipid accumulation in macrophages and contributes to atherosclerosis in apolipoprotein E-deficient mice. FASEB J. 2017;31(11):4971–4984. doi:10.1096/fj.201700235R

215. Chen Y, Xiang R, Zhao S. The potential role of RTN3 in monocyte recruitment and atherosclerosis. Mol Cell Biochem. 2012;361(1-2):67–70. doi:10.1007/s11010-011-1089-6

216. Wang X, Meng H, Ruan J, Chen W, Meng F. Low G0S2 gene expression levels in peripheral blood may be a genetic marker of acute myocardial infarction in patients with stable coronary atherosclerotic disease: A retrospective clinical study. Medicine (Baltimore). 2021;100(3):e23468. doi:10.1097/MD.0000000000023468

217. Hernández-Romero D, Ruiz-Nodar JM, Marín F, Tello-Montoliu A, Roldán V, Mainar L, Pérez-Andreu V, Antón AI, Bonaque JC, Valdés M, et al. CALU A29809G polymorphism in coronary atherothrombosis: Implications for coronary calcification and prognosis. Ann Med. 2010;42(6):439–446. doi:10.3109/07853890.2010.499131

218. Meng B, Li Y, Ding Y, Xu X, Wang L, Guo B, Zhu B, Zhang J, Xiang L, Dong J, et al. Myeloid-derived growth factor inhibits inflammation and alleviates endothelial injury and atherosclerosis in mice. Sci Adv. 2021;7(21):eabe6903. doi:10.1126/sciadv.abe6903

219. Grandoch M, Feldmann K, Göthert JR, Dick LS, Homann S, Klatt C, Bayer JK, Waldheim JN, Rabausch B, Nagy N, et al. Deficiency in lymphotoxin β receptor protects from atherosclerosis in apoE-deficient mice. Circ Res. 2015;116(8):e57–e68. doi:10.1161/CIRCRESAHA.116.305723

220. Shyu HY, Fong CS, Fu YP, Shieh JC, Yin JH, Chang CY, Wang HW, Cheng CW. Genotype polymorphisms of GGCX, NQO1, and VKORC1 genes associated with risk susceptibility in patients with large-artery atherosclerotic stroke. Clin Chim Acta. 2010;411(11-12):840–845. doi:10.1016/j.cca.2010.02.071

221. Tian S, Cao Y, Wang J, Bi Y, Zhong J, Meng X, Sun W, Yang R, Gan L, Wang X, et al. The miR-378c-Samd1 circuit promotes phenotypic modulation of vascular smooth muscle cells and foam cells formation in atherosclerosis lesions. Sci Rep. 2021;11(1):10548. doi:10.1038/s41598-021-89981-z

222. Su YR, Dove DE, Major AS, Hasty AH, Boone B, Linton MF, Fazio S. Reduced ABCA1-mediated cholesterol efflux and accelerated atherosclerosis in apolipoprotein E-deficient mice lacking macrophage-derived ACAT1. Circulation. 2005;111(18):2373–2381. doi:10.1161/01.CIR.0000164236.19860.13

223. Vozenilek AE, Vetkoetter M, Green JM, Shen X, Traylor JG, Klein RL, Orr AW, Woolard MD, Krzywanski DM. Absence of Nicotinamide Nucleotide Transhydrogenase in C57BL/6J Mice Exacerbates Experimental Atherosclerosis. J Vasc Res. 2018;55(2):98–110. doi:10.1159/000486337

224. Zhang H, Ge S, Ni B, He K, Zhu P, Wu X, Shao Y. Augmenting ATG14 alleviates atherosclerosis and inhibits inflammation via promotion of autophagosome-lysosome fusion in macrophages. Autophagy. 2021;1–13. doi:10.1080/15548627.2021.1909833

225. Xiang R, Fan LL, Huang H, Chen YQ, He W, Guo S, Li JJ, Jin JY, Du R, Yan R, et al. Increased Reticulon 3 (RTN3) Leads to Obesity and Hypertriglyceridemia by Interacting With Heat Shock Protein Family A (Hsp70) Member 5 (HSPA5). Circulation. 2018;138(17):1828–1838. doi:10.1161/CIRCULATIONAHA.117.030718

226. Pigeyre M, Bokor S, Romon M, Gottrand F, Gilbert CC, Valtueña J, Gómez-Martínez S, Moreno LA, Amouyel P, Dallongeville J, et al. Influence of maternal educational level on the association between the rs3809508 neuromedin B gene polymorphism and the risk of obesity in the HELENA study. Int J Obes (Lond). 2010;34(3):478–486. doi:10.1038/ijo.2009.260

227. Belalcazar LM, Papandonatos GD, McCaffery JM, Peter I, Pajewski NM, Erar B, Allred ND, Balasubramanyam A, Bowden DW, Brautbar A, et al. A common variant in the CLDN7/ELP5 locus predicts adiponectin change with lifestyle intervention and improved fitness in obese individuals with diabetes. Physiol Genomics. 2015;47(6):215–224. doi:10.1152/physiolgenomics.00109.2014

228. Parente DJ, Garriga C, Baskin B, Douglas G, Cho MT, Araujo GC, Shinawi M. Neuroligin 2 nonsense variant associated with anxiety, autism, intellectual disability, hyperphagia, and obesity. Am J Med Genet A. 2017;173(1):213–216. doi:10.1002/ajmg.a.37977

229. Jenkins D, Seelow D, Jehee FS, Perlyn CA, Alonso LG, Bueno DF, Donnai D, Josifova D, Mathijssen IM, Morton JE, et al. RAB23 mutations in Carpenter syndrome imply an unexpected role for hedgehog signaling in cranial-suture development and obesity. Am J Hum Genet. 2007;80(6):1162–1170. doi:10.1086/518047

230. Zusi C, Morandi A, Maguolo A, Corradi M, Costantini S, Mosca A, Crudele A, Mantovani A, Alisi A, Miraglia Del Giudice E, et al. Association between MBOAT7 rs641738 polymorphism and non-alcoholic fatty liver in overweight or obese children. Nutr Metab Cardiovasc Dis. 2021;31(5):1548–1555. doi:10.1016/j.numecd.2021.01.020

231. Yıldız Bölükbaş E, Mumtaz S, Afzal M, Woehlbier U, Malik S, Tolun A. Homozygous mutation in CEP19, a gene mutated in morbid obesity, in Bardet-Biedl syndrome with predominant postaxial polydactyly. J Med Genet. 2018;55(3):189–197. doi:10.1136/jmedgenet-2017-104758

232. Andreasen CH, Mogensen MS, Borch-Johnsen K, Sandbaek A, Lauritzen T, Sørensen TI, Hansen L, Almind K, Jørgensen T, Pedersen O, et al. Non-replication of genome-wide based associations between common variants in INSIG2 and PFKP and obesity in studies of 18,014 Danes. PLoS One. 2008;3(8):e2872. doi:10.1371/journal.pone.0002872

233. Tian N, Liu Q, Li Y, Tong L, Lu Y, Zhu Y, Zhang P, Chen H, Hu L, Meng J, et al. Transketolase Deficiency in Adipose Tissues Protects Mice From Diet-Induced Obesity by Promoting Lipolysis. Diabetes. 2020;69(7):1355–1367. doi:10.2337/db19-1087

234. Jang I, Pottekat A, Poothong J, Yong J, Lagunas-Acosta J, Charbono A, Chen Z, Scheuner DL, Liu M, Itkin-Ansari P, et al. PDIA1/P4HB is required for efficient proinsulin maturation and ß cell health in response to diet induced obesity. Elife. 2019;8:e44528. doi:10.7554/eLife.44528

235. Ohkubo Y, Sekido T, Nishio SI, Sekido K, Kitahara J, Suzuki S, Komatsu M. Loss of μ ystallin causes PPARγ activation and obesity in high-fat diet-fed mice. Biochem Biophys Res Commun. 2019;508(3):914–920. doi:10.1016/j.bbrc.2018.12.038

236. Simmons RM, McKnight SM, Edwards AK, Wu G, Satterfield MC. Obesity increases hepatic glycine dehydrogenase and aminomethyltransferase expression while dietary glycine supplementation reduces white adipose tissue in Zucker diabetic fatty rats. Amino Acids. 2020;52(10):1413–1423. doi:10.1007/s00726-020-02901-9

237. Kim HR, Jin HS, Eom YB. Association of MACROD2 gene variants with obesity and physical activity in a Korean population. Mol Genet Genomic Med. 2021;9(4):e1635. doi:10.1002/mgg3.1635

238. Williams MJ, Eriksson A, Shaik M, Voisin S, Yamskova O, Paulsson J, Thombare K, Fredriksson R, Schiöth HB. The Obesity-Linked Gene Nudt3 Drosophila Homolog Aps Is Associated With Insulin Signaling. Mol Endocrinol. 2015;29(9):1303–1319. doi:10.1210/ME.2015-1077

239. Arrabal S, Lucena MA, Canduela MJ, Ramos-Uriarte A, Rivera P, Serrano A, Pavón FJ, Decara J, Vargas A, Baixeras E, et al. Pharmacological Blockade of Cannabinoid CB1 Receptors in Diet-Induced Obesity Regulates Mitochondrial Dihydrolipoamide Dehydrogenase in Muscle. PLoS One. 2015;10(12):e0145244. doi:10.1371/journal.pone.0145244

240. Lu Y, Habtetsion TG, Li Y, Zhang H, Qiao Y, Yu M, Tang Y, Zhen Q, Cheng Y, Liu Y. Association of NCOA2 gene polymorphisms with obesity and dyslipidemia in the Chinese Han population. Int J Clin Exp Pathol. 2015;8(6):7341–7349.

241. Bendlová B, Vaň vá M, Hill M, Vacínová G, Lukášová P, VejraŽková D, Šedová L, Šeda O, Včelák J. ZBTB16 gene variability influences obesity-related parameters and serum lipid levels in Czech adults. Physiol Res. 2017;66(Suppl 3):S425–S431. doi:10.33549/physiolres.933731

242. Vernochet C, Mourier A, Bezy O, Macotela Y, Boucher J, Rardin MJ, An D, Lee KY, Ilkayeva OR, Zingaretti CM, et al. Adipose-specific deletion of TFAM increases mitochondrial oxidation and protects mice against obesity and insulin resistance. Cell Metab. 2012;16(6):765–776. doi:10.1016/j.cmet.2012.10.016

243. Zhu X, Xie S, Xu T, Wu X, Han M. Rasal2 deficiency reduces adipogenesis and occurrence of obesity-related disorders. Mol Metab. 2017;6(6):494–502. doi:10.1016/j.molmet.2017.03.003

244. Lomba A, Milagro FI, García-Díaz DF, Campión J, Marzo F, Martínez JA. A high-sucrose isocaloric pair-fed model induces obesity and impairs NDUFB6 gene function in rat adipose tissue. J Nutrigenet Nutrigenomics. 2009;2(6):267–272. doi:10.1159/000308465

245. Yoshino S, Satoh T, Yamada M, Hashimoto K, Tomaru T, Katano-Toki A, Kakizaki S, Okada S, Shimizu H, Ozawa A, et al. Protection against high-fat diet-induced obesity in Helz2-deficient male mice due to enhanced expression of hepatic leptin receptor. Endocrinology. 2014;155(9):3459–3472. doi:10.1210/en.2013-2160

246. Deng T, Lyon CJ, Minze LJ, Lin J, Zou J, Liu JZ, Ren Y, Yin Z, Hamilton DJ, Reardon PR, et al. Class II major histocompatibility complex plays an essential role in obesity-induced adipose inflammation. Cell Metab. 2013;17(3):411–422. doi:10.1016/j.cmet.2013.02.009

247. Zheng W, Yan C, Wang X, Luo Z, Chen F, Yang Y, Liu D, Gai X, Hou J, Huang M. TheTGFB1 functional polymorphism rs1800469 and susceptibility to atrial fibrillation in two Chinese Han populations. PLoS One. 2013;8(12):e83033. doi:10.1371/journal.pone.0083033

248. Choudhury A, Chung I, Blann AD, Lip GY. Platelet surface CD62P and CD63, mean platelet volume, and soluble/platelet P-selectin as indexes of platelet function in atrial fibrillation: a comparison of “healthy control subjects” and “disease control subjects” in sinus rhythm. J Am Coll Cardiol. 2007;49(19):1957–1964. doi:10.1016/j.jacc.2007.02.038

249. Hou J, Yue Y, Hu B, Xu G, Su R, Lv L, Huang J, Yao J, Guan Y, Wang K, et al. DACT1 Involvement in the Cytoskeletal Arrangement of Cardiomyocytes in Atrial Fibrillation by Regulating Cx43. Braz J Cardiovasc Surg. 2019;34(6):711–722. doi:10.21470/1678-9741-2019-0033

250. Li J, Wang S, Bai J, Yang XL, Zhang YL, Che YL, Li HH, Yang YZ. Novel Role for the Immunoproteasome Subunit PSMB10 in Angiotensin II-Induced Atrial Fibrillation in Mice. Hypertension. 2018;71(5):866–876. doi:10.1161/HYPERTENSIONAHA.117.10390

251. Pollak AJ, Haghighi K, Kunduri S, Arvanitis DA, Bidwell PA, Liu GS, Singh VP, Gonzalez DJ, Sanoudou D, Wiley SE, et al. Phosphorylation of serine96 of histidine-rich calcium-binding protein by the Fam20C kinase functions to prevent cardiac arrhythmia. Proc Natl Acad Sci U S A. 2017;114(34):9098–9103. doi:10.1073/pnas.1706441114

252. Cheng WL, Kao YH, Chen YC, Lin YK, Chen SA, Chen YJ. Macrophage migration inhibitory factor increases atrial arrhythmogenesis through CD74 signaling. Transl Res. 2020;216:43–56. doi:10.1016/j.trsl.2019.10.002

253. Khalyfa A, Kheirandish-Gozal L, Bhattacharjee R, Khalyfa AA, Gozal D. Circulating microRNAs as Potential Biomarkers of Endothelial Dysfunction in Obese Children. Chest. 2016;149(3):786–800. doi:10.1378/chest.15-0799

254. Zamanillo R, Sánchez J, Serra F, Palou A. Breast Milk Supply of MicroRNA Associated with Leptin and Adiponectin Is Affected by Maternal Overweight/Obesity and Influences Infancy BMI. Nutrients. 2019;11(11):2589. doi:10.3390/nu11112589

255. Zhou Y, Hambly BD, Simmons D, McLachlan CS. RUNX1T1 rs34269950 is associated with obesity and metabolic syndrome. QJM. 2020;hcaa208. doi:10.1093/qjmed/hcaa208

256. Zhuo XZ, Bai K, Wang Y, Liu P, Xi W, She J, Liu J. Long-chain non-coding RNA-GAS5 / hsa-miR-138-5p attenuates high glucose-induced cardiomyocyte damage by targeting CYP11B2. Biosci Rep. 2021;BSR20202232. doi:10.1042/BSR20202232

257. Wu L, Archacki SR, Zhang T, Wang QK. Induction of high STAT1 expression in transgenic mice with LQTS and heart failure. Biochem Biophys Res Commun. 2007;358(2):449–454. doi:10.1016/j.bbrc.2007.04.119

258. Mirza AH, Kaur S, Nielsen LB, Størling J, Yarani R, Roursgaard M, Mathiesen ER, Damm P, Svare J, Mortensen HB, et al. Breast Milk-Derived Extracellular Vesicles Enriched in Exosomes From Mothers With Type 1 Diabetes Contain Aberrant Levels of microRNAs. Front Immunol. 2019;10:2543. doi:10.3389/fimmu.2019.02543

259. Brizzi MF, Dentelli P, Pavan M, Rosso A, Gambino R, Grazia De Cesaris M, Garbarino G, Camussi G, Pagano G, Pegoraro L. Diabetic LDL inhibits cell-cycle progression via STAT5B and p21(waf). J Clin Invest. 2002;109(1):111–119. doi:10.1172/JCI13617

260. Peng J, Xiang Y. Value analysis of CD69 combined with EGR1 in the diagnosis of coronary heart disease. Exp Ther Med. 2019;17(3):2047–2052. doi:10.3892/etm.2019.7175

261. Muiya NP, Wakil S, Al-Najai M, Tahir AI, Baz B, Andres E, Al-Boudari O, Al-Tassan N, Al-Shahid M, Meyer BF, et al. A study of the role of GATA2 gene polymorphism in coronary artery disease risk traits. Gene. 2014;544(2):152–158. doi:10.1016/j.gene.2014.04.064

262. Alfonso-Jaume MA, Bergman MR, Mahimkar R, Cheng S, Jin ZQ, Karliner JS, Lovett DH. Cardiac ischemia-reperfusion injury induces matrix metalloproteinase-2 expression through the AP-1 components FosB and JunB. Am J Physiol Heart Circ Physiol. 2006;291(4):H1838–H1846. doi:10.1152/ajpheart.00026.2006

263. Wang B, Xu A. Aryl hydrocarbon receptor pathway participates in myocardial ischemia reperfusion injury by regulating mitochondrial apoptosis. Med Hypotheses. 2019;123:2–5. doi:10.1016/j.mehy.2018.12.004

264. Mahmoud MM, Kim HR, Xing R, Hsiao S, Mammoto A, Chen J, Serbanovic-Canic J, Feng S, Bowden NP, Maguire R, et al. TWIST1 Integrates Endothelial Responses to Flow in Vascular Dysfunction and Atherosclerosis. Circ Res. 2016;119(3):450–462. doi:10.1161/CIRCRESAHA.116.308870

